# GABARAP membrane conjugation sequesters the FLCN-FNIP tumor suppressor complex to activate TFEB and lysosomal biogenesis

**DOI:** 10.1101/2021.02.22.432209

**Authors:** Jonathan M. Goodwin, Ward G. Walkup, Kirsty Hooper, Taoyingnan Li, Chieko Kishi-Itakura, Aylwin Ng, Timothy Lehmberg, Archana Jha, Sravya Kommineni, Katherine Fletcher, Jorge Garcia-Fortanet, Yaya Fan, Qing Tang, Menghao Wei, Asmita Agrawal, Sagar R. Budhe, Sreekanth R. Rouduri, Dan Baird, Jeff Saunders, Janna Kiselar, Mark R. Chance, Andrea Ballabio, Brent A. Appleton, John H. Brumell, Oliver Florey, Leon O. Murphy

## Abstract

Adaptive changes in lysosomal capacity are driven by the transcription factors TFEB and TFE3 in response to increased autophagic flux and endolysosomal stress, yet the molecular details of their activation are unclear. LC3 and GABARAP members of the ATG8 protein family are required for selective autophagy and sensing perturbation within the endolysosomal system. Here we show that during single membrane ATG8 conjugation (SMAC), Parkin-dependent mitophagy, and *Salmonella*-induced xenophagy, the membrane conjugation of GABARAP, but not LC3, is required for activation of TFEB/TFE3 to control lysosomal homeostasis and capacity. GABARAP directly binds to a novel LC3-interacting motif (LIR) in the FLCN/FNIP tumor suppressor complex with picomolar affinity and regulates its relocalization to these GABARAP-conjugated membrane compartments. This disrupts the regulation of RagC/D by the FLCN/FNIP GAP complex, resulting in impaired mTOR-dependent phosphorylation of TFEB without changing mTOR activity towards other substrates. Thus, the GABARAP-FLCN/FNIP-TFEB axis serves as a universal molecular sensor that coordinates lysosomal homeostasis with perturbations and cargo flux within the autophagy-lysosomal network.

## Introduction

Within the cell, the proteasome and the lysosome serve as the degradative hubs for both intracellular and extracellular cargo to maintain proteostasis and cellular fitness^1^. The lysosome is an acidic organelle that compartmentalizes hydrolytic enzymes capable of degrading complex organelles, protein aggregates, pathogens, complex lipids, as well as dysfunctional proteins^2^. Lysosomes can receive cargo from fusion with autophagosomes (macroautophagy), endosomes (endocytosis), phagosomes (phagocytosis and LC3-associated phagocytosis), engulfment through lysosomal microautophagy, or through import of proteins during chaperone-mediated autophagy^3–9^. Within the lysosome, cargo can be broken down and substituents are then recycled to help the cell adapt to stress, or processed to for antigen presentation or lysosomal exocytosis^4, 10, 11^. Apart from the enzymatic digestion of proteins and other cargo, lysosomes are important intracellular stores for cations such as calcium, iron, and sodium but also H^+^ protons which are critical for maintaining lysosomal pH. Ion channels and proton pumps present on the lysosomal membrane are essential for lysosomal function, positioning, membrane fusion, and host-pathogen responses^12^.

While an important pillar of proteostasis, the lysosome is also a critical regulator of cellular metabolism. The mechanistic target of rapamycin (mTOR) complex 1 (mTORC1) resides on the lysosomal membrane and coordinates anabolic processes in response to changes in nutrient availability^13^. Lysosomal localization of mTORC1 is controlled by a signaling platform consisting of the vATPase, the Ragulator complex (LAMTOR1-5), and the transmembrane protein SLC38A9, which together mediates the localization and activation state of the Rag GTPases^14, 15^. Functioning as obligate heterodimers, the GTP-loaded state of RagA/B and RagC/D is tightly controlled by the GATOR1 and FLCN GTPase-activating complexes, respectively. Active RagGTPases (where RagA/B is in the GTP-bound state) recruits mTORC1 to the lysosomal membrane and modulates substrate phosphorylation^15^.

In addition to regulation of cellular metabolism, mTORC1 exerts effects on the lysosomal network itself through the regulation of the TFE/MiTF bHLH transcription factor family^16^. The family members TFE3 and TFEB are well documented as master regulators of lysosomal biogenesis and autophagy^17–19^. In the case of macroautophagy, which is thought to have evolved as a stress response to starvation, mTORC1 functions to suppress both autophagosome biogenesis and TFE3/TFEB activation during nutrient availability. However, upon inhibition of mTORC1 by nutrient deprivation, increased autophagosome biogenesis is coupled with activation of TFE3/TFEB to drive lysosomal biogenesis and facilitate flux. Importantly, nutrient deprivation switches nucleotide binding on RagC/D to the GTP-bound state through formation of the lysosomal folliculin complex (LFC)^20, 21^. When GTP-Bound, RagC/D can no longer directly bind TFE3 and TFEB to present them as substrates to mTOR, a mechanism not required for other mTOR substrates such as S6K^22^. While this lysosome-to-nucleus signaling mechanism^23^ is documented during nutrient starvation, how TFE3/TFEB activation is coordinated with the lysosomal delivery pathways mentioned above, as well as specialized forms of selective autophagy, is unclear.

Conjugation of ATG8 homologs (e.g., LC3 and GABARAP proteins) to double membranes during autophagosome biogenesis mediates phagosome maturation and lysosomal delivery of cytosolic contents for degradation and recycling^3^. Alternatively, ATG8 proteins can also be conjugated to single-membrane organelles within the endocytic system, but the functional consequence of this is not well understood^24^. Single-membrane ATG8 conjugation, referred to here as SMAC, occurs during LC3-associated phagocytosis (LAP)^6^, LC3-associated endocytosis (LANDO)^8^, and upon perturbation of endolysosomal ion gradients during pathogen infection^25, 26^. SMAC can be pharmacologically induced by chemical probes that exhibit lysosomotropic and broad ionophore/protonophore-like properties but unfortunately these agents lack a molecular target^7^. Recently, it was reported that ATG8 proteins are directly conjugated to the lysosomal membrane upon disruption of lysosomal homeostasis by agents such as LLoMe, oxalate crystals, and membrane-permeabilizing pathogen virulence factors^27, 28^. This autophagy-independent ATG8 conjugation was required for the coordinated activation of TFEB, specifically uncoupling TFEB, but not other substrates, from regulation by mTOR. This prompts the question whether the commonality of ATG8 conjugation across lysosomal delivery pathways serves as a universal mechanism to coordinate TFE3/TFEB activation.

To uncover the novel mechanism by which ATG8 conjugation impacts TFEB activation, we characterized TRPML1 channel agonists as the first pharmacological probes with a defined molecular target that can be deployed as SMAC agonists, in contrast to non-specific ionophore/protonophores^7^. Regulating ATG8 conjugation in this way allowed us to reveal that GABARAP proteins directly bind and sequester the FLCN GAP complex on different membrane compartments. This disrupts regulation of RagC/D and liberates TFEB from control by mTORC1 to coordinate lysosomal biogenesis with organelle perturbations and elevations in autophagic cargo flux. This represents a TFEB transcriptional activation paradigm distinct from the setting of nutrient starvation, where broad inhibition of mTOR activity towards downstream substrates such as S6K and 4E-BP1 results in a concurrent block in protein translation.

## Results

### TRPML1 agonists stimulate single membrane ATG8 conjugation (SMAC)

Given the complexity of lysosomal membrane disrupting agents, we chose to acutely alter ion concentrations within the lysosome by harnessing pharmacological agonists of the lysosomal transient receptor potential mucolipin channel 1 (TRPML1). Treatment with the TRPML1 agonists MK6-83^29^, ML-SA1^30^ or a recently published, more potent channel agonist (designated as compound 8 “C8”)^31^ resulted in the rapid conversion of LC3 from its cytoplasmic “I” form to the lipidated, punctate “II” form in both wild type (WT) and autophagy-deficient cells (ATG13_KO). In contrast, the mTOR inhibitor AZD8055, a well-established agent to induce autophagosome biogenesis^32^, was able to regulate LC3 lipidation in WT but not autophagy-deficient cells (Fig. 1A, B, Fig. S1A, B, Movie S1, S2). AZD8055 or EBSS starvation induced conversion of LC3 which was sensitive to the VPS34 inhibitor PIK-III and potentiated with the vATPase inhibitor Bafilomycin A1 (BafA1) (Fig. 1C). Interestingly, treatment with C8 robustly induced VPS34-independent LC3 lipidation that was inhibited by BafA1 with no impact on mTOR activity (Fig. 1C). This rapid lipidation depends on TRPML1 (Fig. S2A) and is not accompanied by lysosomal alkalization or membrane damage (Fig. S3A, B). Together, these features are characteristic of SMAC where ATG8s are conjugated to endolysosomal membranes^33^. Consistent with this, TRPML1 agonist treatment also induced strong colocalization of ATG8s (LC3B or GABARAPL1) with the lysosomal marker LAMP1 (Fig. 1D-F).

**Figure 1.**
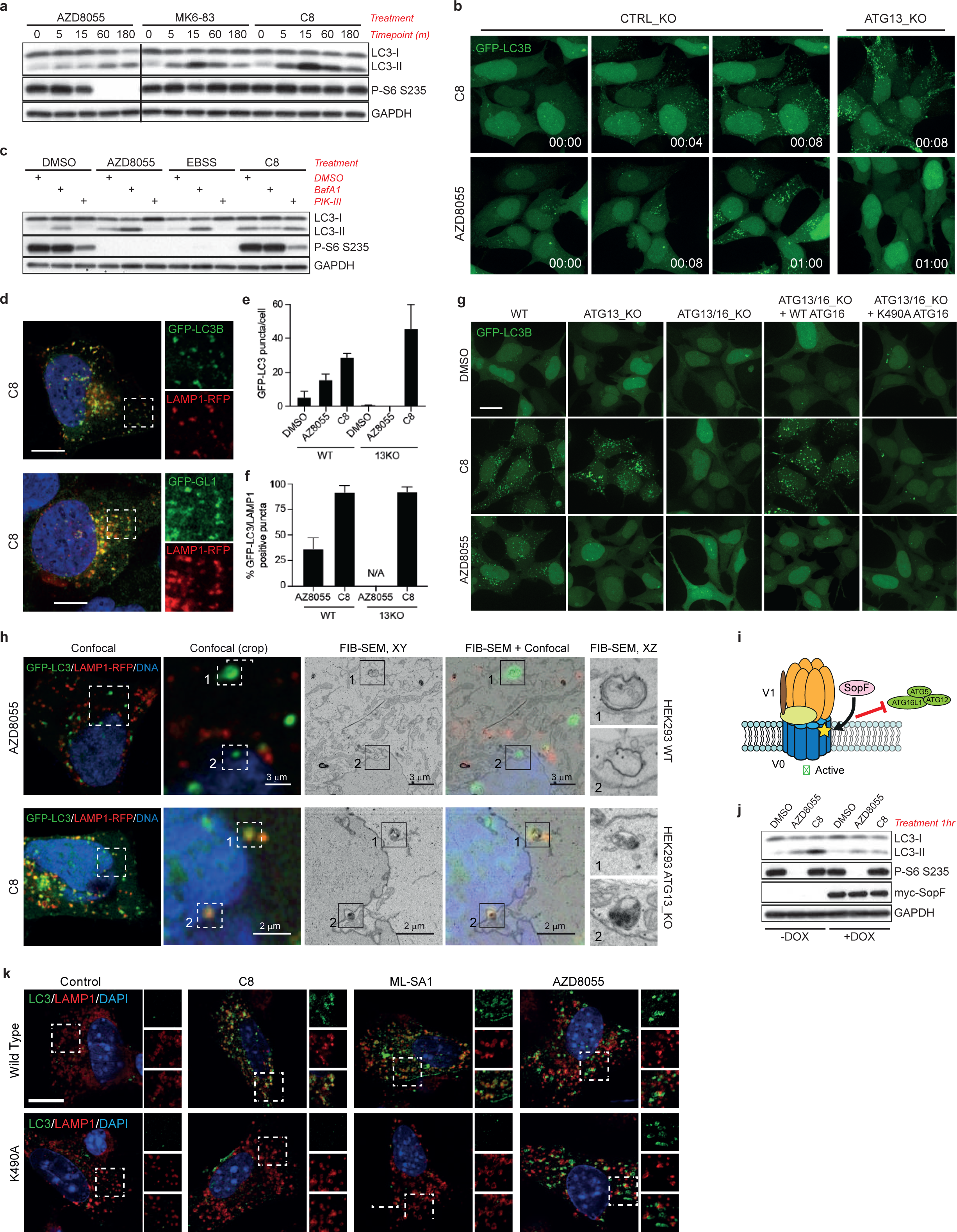
Activation of the lysosomal ion channel TRPML1 results in ATG8 conjugation to the lysosomal membrane independent of autophagy. **(a)** Western blot analysis of LC3 lipidation kinetics upon treatment with the mTOR inhibitor AZD8055 or the TRPML1 agonists MK6-83 (25uM) or C8 (2uM) for the indicated timepoints. **(b)** Time-lapse imaging of GFP-LC3B in WT or ATG13_KO HEK2393T cells. Cells were treated with C8 (2uM) or AZD8055 (1uM). Still images at the indicated timepoints (min) are shown. **(c)** Western blot analysis of LC3 lipidation sensitivity to either BafA1 (100nM) or PIK-III (5uM). Cells were cotreated with the indicated compounds for 2hr. **(d)** Colocalization analysis of GFP-tagged ATG8 homologs with the lysosomal marker LAMP1-RFP in WT HEK293T cells after treatment with C8 (2uM). **(e)** Quantification of GFP-LC3B puncta number in WT or ATG13_KO HEK293T cells treated with AZD8055 (1uM) or C8 (2uM). **(f)** Quantification of acute GFP-LC3B colocalization with RFP-LAMP1 in WT and ATG13_KO HEK293T cells. **(g)** Immunofluorescence analysis of GFP-LC3B puncta formation in WT HEK293T, ATG13_KO, ATG13/ATG16L1_DKO, and DKO cells rescued with either WT- or K490A-ATG16L1. Treatment with AZD8055 displayed puncta only in autophagy-competent cells, whereas the TRPML1 agonist treatment formed puncta in autophagy-deficient cells dependent on the ability of ATG16L1 to interact with single membrane structures. **(h)** Ultrastructural CLEM, FIB-SEM analysis of GFP-LC3B positive cells in HEK293T cells of the indicated genotype. CLEM representative images shown in optimal X/Y resolution. Zoom FIB-SEM images shown in X/Z plane. **(i)** Primary BMDM were treated with the indicated compounds for 1 hr and immunostained for LC3B and the lysosomal marker LAMP1. LC3 puncta generation with AZD8055 was preserved in both genotypes. **(j)** Diagram of SopF impairment of ATG16L1 recruitment by the vATPase. **(k)** Expression of SopF (+DOX) in HeLa cells blocks LC3A/B lipidation upon TRPML1 activation but does not impact autophagosome biogenesis stimulated by AZD8055.

ATG16L1-K490A is a recently discovered allele with a mutation in the C-terminal WD repeats of ATG16L1 required for SMAC but not autophagosome formation^26^. Expression of ATG16L1-K490A in autophagy-deficient cells blocked TRPML1 agonist-induced LC3 puncta formation (Fig. 1G). Using Focused Ion Beam (FIB-SEM) correlative light and electron microscopy (CLEM), TRPML1 agonist induced GFP-LC3 positive structures were identified as single-membrane late endosome/lysosomes as opposed to the double-membrane autophagosomes in AZD8055 treated cells (Fig. 1H, Movie S3, S4).

Next, we asked whether TRPML1 activation could induce SMAC in primary immune cells from a knockin ATG16L1-K490A mouse model. Treatment of ATG16L1^K490A^ or ATG16L1^WT^ primary bone marrow-derived macrophages with AZD8055 showed similar levels of LC3 puncta, but C8 or ML-SA1 were unable to induce LC3 puncta in ATG16L1^K490A^ expressing cells (Fig. 1K). Given the sensitivity to BafA1 and requirement of the ATG16L1 C-terminal WD repeats, we explored the role of vATPase in SMAC. We utilized the *S.* Typhimurium SopF effector protein, which has recently been shown to block interaction between ATG16L1 and the vATPase through ADP-ribosylation of the V0C subunit^25^. Indeed, expression of SopF was sufficient to block LC3-II formation upon TRPML1 activation but not AZD8055 treatment (Fig. 1I, J). Collectively, using genetic, pharmacological and ultrastructural methodologies, these data provide strong evidence that TRPML1 activation can induce SMAC directly on lysosomes.

### SMAC is required for TFEB activation downstream of TRPML1

The activation of TRPML1 upon nutrient starvation is known to result in the nuclear localization of the transcription factors TFEB and TFE3, in part due to local calcium-mediated activation of the phosphatase Calcineurin (CaN) to dephosphorylate TFEB/TFE3^34^. Interestingly, treatment of fed cells with TRPML1 agonists resulted in TFEB nuclear accumulation, with no impact of pharmacological or genetic inhibition of CaN (Fig. S4), suggesting a regulatory mechanism that differs from nutrient starvation^34^. To investigate whether SMAC was involved in TFEB activation, we focused first on a requirement for the vATPase. Acute treatment with BafA1 was sufficient to block TFEB activation by TRPML1 agonists, however it did not impact TFEB activation upon mTOR inhibition (Fig. 2A). Additionally, inhibition of SMAC through expression of SopF blocked TFEB nuclear localization induced by TRPML1 agonist but not AZD8055 treatment (Fig. S5). CRISPR mediated knockout of the ATG8 lipidation machinery such as ATG16L1, ATG5 or ATG7 but not FIP200, ATG9A or VPS34, which are required for autophagosome biogenesis, blocked TRPML1 agonist induced TFEB activation (Fig. 2B). Importantly, lysosomal calcium release stimulated by C8 or the activation of TFEB upon nutrient starvation were insensitive to BafA1 treatment or ATG16L1 knock-out (Fig. 2C, Fig. S6).

**Figure 2.**
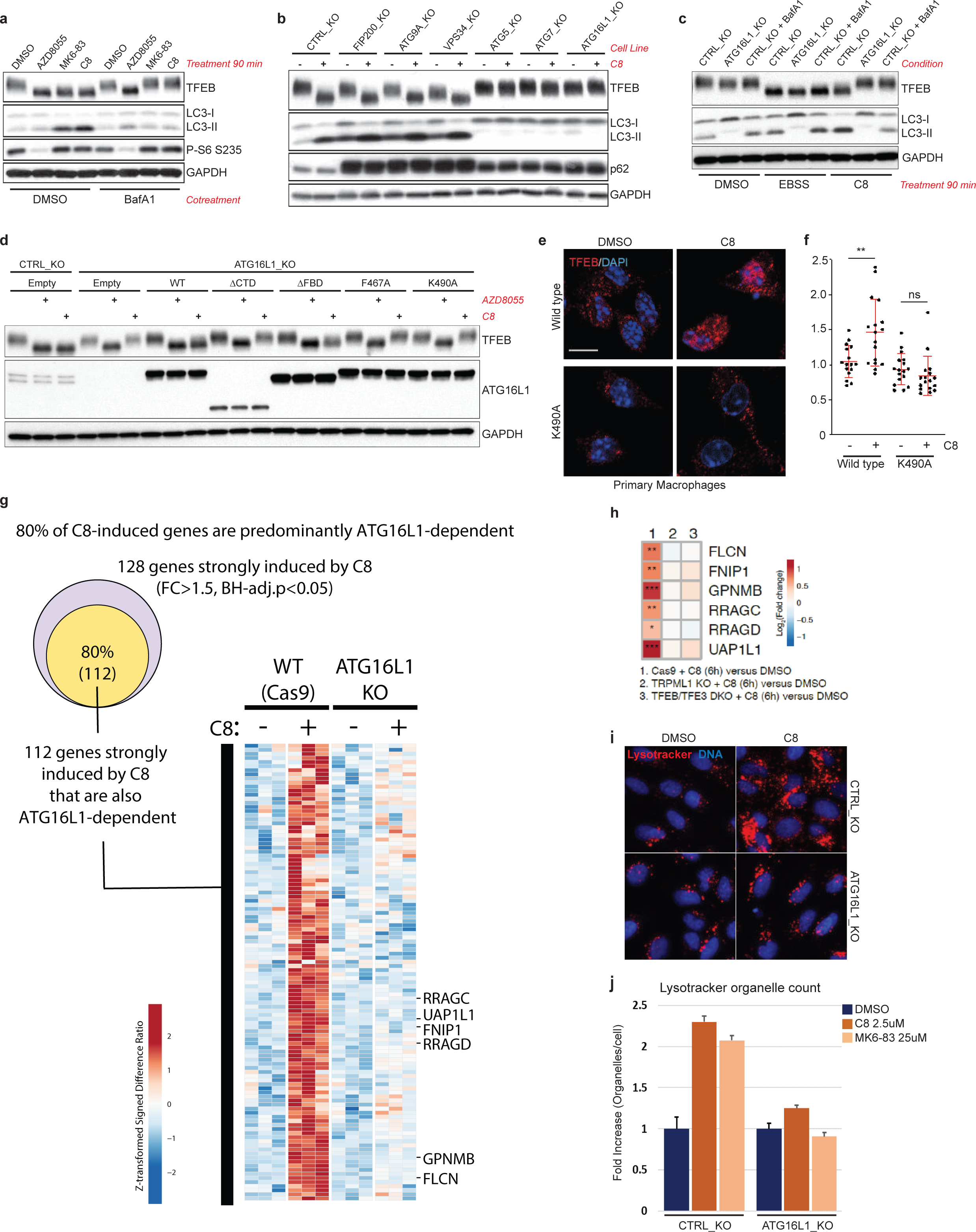
ATG16L1-dependent ATG8 conjugation to single membranes is required for TFEB activation and lysosomal biogenesis upon TRPML1 activation. **(a)** Western blot analysis of TFEB phosphorylation and active state reveals that TRPML1 activation of TFEB requires the vATPase. HeLa cells were treated with the indicated compounds for 90 min. **(b)** CRISPR knockout of the indicated genes in HeLa cells stably expressing Cas9. TRPML1 agonist induced TFEB activation in autophagy-deficient cells, yet was dependent upon the genes required for ATG8 conjugation. **(c)** TFEB activation by a TRPML1 agonist is sensitive to ATG16L1_KO or BafA1 cotreatment, whereas neither of these conditions affects TFEB activation upon nutrient starvation. HeLa.CTRL_KO and ATG16L1_KO cells were treated as indicated. **(d)** Western blot analysis of HeLa cells of the indicated genotype expressing variants of ATG16L1. TFEB activation was strictly dependent of the ability of the ATG16L1 C-terminal WD repeats to interact with single membranes. **(e)** Primary BMDM of the indicated genotype were treated with 2uM C8 for 1 hr and immunostained for endogenous TFEB localization. Nuclear localization was impaired in the ATG16L1^K490A^ cells. **(f)** Quantification of nuclear/cytosolic ratio of TFEB in primary macrophages of the indicated genotype upon C8 treatment. **(g)** RNAseq transcriptomics profiling of genes induced by C8 compound stimulation for 24h in ATG16L1 KO and WT HeLa cells (expressing Cas9). The Venn diagram shows a large proportion (80%) of C8-induced genes that were also ATG16L1 dependent. The heatmap shows Log2(CPM)-derived values for these genes, expressed as Z-transformed signed difference ratios (SDR) relative to their respective unstimulated baseline controls (either ATG16L1 KO or WT) and then scaled by normalizing to the maximum absolute deviation of each gene’s expression level from the unstimulated control. Differentially induced genes were identified by having a fold change (FC) >1.5 and Benjamini and Hochberg’s (BH)-adjusted p<0.05. **(h)** Expression analysis of TFEB target gene panel, which was induced upon TRPML1 agonist (C8) treatment but not in TRPML1_KO and TFEB/TFE3_DKO cells, which were used to highlight specificity of response. RNAseq was performed in triplicates. **(i)** Prolonged treatment with TRPML1 agonist induces lysosomal biogenesis dependent on ATG16L1. U2OS cells of the indicated genotype were treated for 24 hours before staining of the cells with Lysotracker dye. Representative images shown. **(j)** Quantification of Lysotracker staining organelle count after 24 hr treatment with the indicated compounds. Fold change organelles per cell ± SD.

Consistent with modulation of SMAC by SopF expression, ATG16L1_KO cells re-expressing WT or an autophagy deficient ATG16L1 FIP200 binding mutant (FBD) supported TRPML1 induced TFEB activation, while re-expression of a C-terminal domain truncation (ΔCTD) or F467A or K490A mutations deficient for SMAC^26^ did not (Fig. 2D, Fig. S7). In contrast, AZD8055 activated TFEB irrespective of ATG16L1 allele status. Furthermore, treatment of ATG16L1^K490A^ or ATG16L1^WT^ primary bone marrow-derived macrophages revealed TRPML1-mediated TFEB activation was abolished in ATG16L1^K490A^ expressing cells (Fig. 2E, F) further confirming the presence of a novel SMAC-regulated pathway that induces TFEB activity in diverse cell types. These data suggest that ATG8 conjugation to single membrane organelles can promote activation of the TFEB/TFE3 transcription factors through a vATPase-dependent, ATG8-dependent mechanism which is distinct to the regulation of TFEB in the context of nutrient starvation or mTOR inhibition.

Upon nuclear localization, TFEB serves as the primary transcription factor responsible for lysosomal biogenesis^16^. Remarkably, the TRPML1-dependent transcriptomic response was largely dependent on ATG16L1 and included numerous TFEB-target genes involved in lysosomal function (Fig. 2G, H, Fig. S8). Consistent with this profile, we found that TRPML1 activation by treatment with C8 for 24 hours increased the number of Lysotracker-positive organelles in an ATG16L1-dependent manner (Fig. 2I, J). Together, these observations demonstrate that following changes in lysosomal ion balance, the WD40 domain in ATG16L1 regulates lysosomal SMAC and that this is required for TFEB activation and lysosomal biogenesis.

### GABARAPs selectively bind and sequester the FLCN-FNIP tumor suppressor complex to lysosomes

Mammalian ATG8 homologs consist of 3 members of the MAP1LC3 family (LC3A/B/C) and 3 members of the GABA type A Receptor-Associated Protein family (GABARAP/L1/L2)^35^. Using a combinatorial CRISPR knockout approach, the GABARAP subfamily were found to be essential for the TRPML1-mediated activation of TFEB (Fig. 3A). In agreement with protein interaction databases linking GABARAP, but not LC3 proteins, with the TFEB regulators FLCN and FNIP1/2^36–38^, we confirmed that the FLCN-FNIP1 complex interacts with GABARAP through LIR-dependent binding (Fig. 3B-D, Fig. S9). The connection between GABARAP and the FLCN/FNIP complex was intriguing given the role that the FLCN/FNIP substrate RagC/D plays in regulating TFEB^39^.

**Figure 3.**
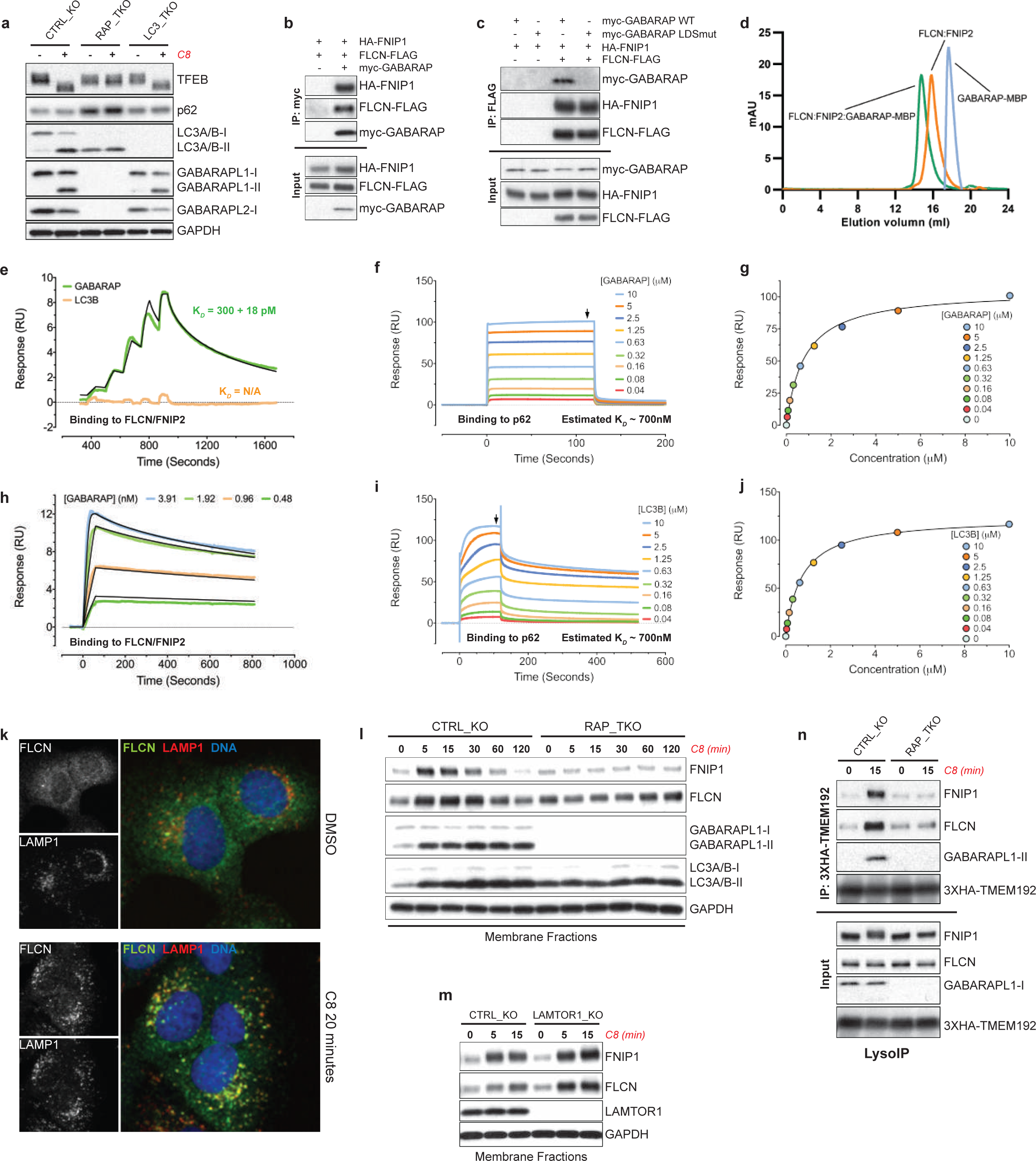
GABARAP is required for TFEB activation and FLCN-FNIP sequestration upon acute TRPML1 stimulation. **(a)** Combinatorial CRISPR KO of ATG8 homologs in HeLa cells stably expressing Cas9. RAP_TKO = GABARAP, GABARAPL1, and GABARAPL2 knockout. LC3_TKO = LC3A, LC3B, and LC3C knockout. Western blot analysis of cells treated with TRPML1 agonist reveals GABARAP proteins are specifically required for TFEB activation. **(b)** GABARAP binds the FLCN-FNIP1 complex. HEK293T cells transiently transfected with the indicated constructs for 20hr before lysis and immunoprecipitation. **(c)** Interaction of GABARAP with FLCN-FNIP1 complex requires LIR binding motif of GABARAP. HEK293T cells transiently transfected with the indicated constructs for 20hr before lysis and immunoprecipitation. LIR binding motif (LBM) mutant = K39Q/Y40H/Q74E/F75L. **(d)** GABARAP_MBP complexes with FLCN/FNIP2 over a Superose 6 column. Overlay of individual and complexed chromatograms. **(e)** GABARAP binds FLCN/FNIP2 with picomolar affinity (300 pM K_*D*_) in single cycle SPR, while LC3B does not show binding to FLCN/FNIP2. **(f)** GABARAP binds FLCN/FNIP2 with picomolar affinity in multi-cycle SPR. **(g, h)** GABARAP and **(i, j)** LC3B bind to the FIR domain of p62 with equivalent affinity (700 nM K_*D*_) in multi-cycle SPR. **(k)** Immunofluorescence analysis of U2OS cells treated with TRPML1 agonist for 20 minutes. Images show colocalization between endogenous FLCN and LAMP1. **(l)** Western blot analysis of HeLa WT or GABARAP_TKO membrane fractions after treatment with TRPML1 agonist for the indicated timepoints. **(m)** Western blot analysis of FLCN-FNIP1 membrane recruitment upon TRPML1 agonist treatment in LAMTOR1_KO cells. **(n)** Purification of lysosomes using LysoIP reveals acute GABARAP-dependent lysosomal FLCN-FNIP sequestration. U2OS cells of the indicated genotype were stably transduced with 3XHA-TMEM192 expression construct. LysoIP was performed after treatment with C8 (2uM, 15 min).

We next determined the binding affinity of the interaction between full length GABARAP and FLCN/FNIP2 by Surface Plasmon Resonance (SPR). GABARAP bound to immobilized FLCN/FNIP2 with a picomolar affinity in single cycle (300 ± 18 pM K_D_) and multi-cycle kinetics SPR formats (71 ± 10 pM K_D_), whereas LC3B did not bind to FLCN/FNIP2 under identical assay conditions (Fig. 3E, F). To confirm the functionality of the full length LC3B, we used multi-cycle SPR to measure the affinity of GABARAP and LC3B to immobilized p62 FIR domain. Both GABARAP (670 ± 80 nM K_*D*_) and LC3B (820 ± 40 nM K_*D*_) bound p62 with nanomolar affinities in steady state SPR measurements (Fig. 3G-J) which is consistent with previous reports^40^. Taken together, the specific binding of GABARAP to FLCN/FNIP2, with an unusually high affinity for an ATG8-binding event, is consistent with the important role GABARAP plays in regulating TFEB activation described above.

We hypothesized direct conjugation of GABARAPs to the lysosomal membrane could re-distribute the FLCN/FNIP complex. Indeed, following TRPML1 activation, there was a rapid and robust increase in membrane-associated FLCN and FNIP1 and this was dependent on the GABARAP proteins and vATPase function (Fig. 3L, Fig. S10). FLCN colocalization with the lysosomal marker LAMP-1 (Fig. 3K) and its presence on isolated lysosomes was dependent on GABARAP proteins (Fig. 3N). Lysosomal localization of FLCN also occurs upon nutrient starvation, where FLCN specifically binds to RagA^GDP^ and forms the inhibitory lysosomal folliculin complex (LFC)^20, 21^. However, in cells deficient for the Ragulator complex component LAMTOR1, which is part of the LFC, TRPML1 activation promoted FLCN membrane recruitment (Fig. 3M). Additionally, using NPRL2_KO cells which have constitutive RagA/B^GTP^ and defective lysosomal localization of FLCN upon starvation^41^, we found that the FLCN-FNIP1 complex distributed to membranes following TRPML1 activation (Fig. S11). These data indicate that the GABARAP-dependent sequestration of FLCN/FNIP1 is a process distinct from LFC formation.

We reasoned that GABARAP-dependent recruitment of FLCN to the lysosome could inhibit its GAP activity towards RagC/RagD, the heterodimeric partner of RagA/B, analogous to how lysosomal recruitment inhibits FLCN-FNIP GAP activity during LFC formation^20, 21^. In this model, FLCN-FNIP1 would normally exert its GAP function away from the lysosomal surface, thus promoting cytosolic RagC/D^GDP^ and subsequent TFEB cytosolic retention^22, 39^. Indeed, RagGTPase dimers have been shown to interact dynamically with the lysosome under fed conditions^42, 43^. Using NPRL2_KO cells mentioned above, additional knockout of FLCN resulted in complete nuclear localization of TFEB under nutrient rich conditions in both wild type and LFC-deficient NPRL2_KO cells, supporting a model where FLCN-FNIP1 GAP activity towards RagC/D can occur outside the context of the LFC (Fig. S12). In a complementary approach, we artificially tethered FLCN to the lysosomal surface (lyso-FLCN, see methods) and found this sufficient to activate TFEB and TFE3 in full nutrient conditions without impacting mTOR signaling to S6K1 (Fig. S13). Additionally, we found that in contrast to nutrient starvation, TRPML1 agonist treatment retained the ability to activate TFEB in NPRL2_KO cells, suggesting intact FLCN GAP activity toward RagGTPases (Fig. S14). However, expression of RagGTPases locked in the active state (RagB^Q99L^/RagD^S77L^), which are no longer regulated by FLCN-FNIP1, suppressed the mobility shift of TFEB and its homolog TFE3 following TRPML1 activation but not with AZD8055 (Fig. S14). Collectively, these data support a model in which the redistribution of the FLCN-FNIP1 complex to the lysosomal membrane can regulate the nucleotide binding state of RagC/D, thus resulting in a novel mechanism for TFEB activation.

### A novel LIR domain in FNIP1/2 mediates high affinity GABARAP interaction

Carboxyl group footprinting was employed as an unbiased approach to identify the molecular interface between GABARAP and the FLCN-FNIP complex^44, 45^. This method also leverages the observation that aspartate and/or glutamate residues are commonly observed upstream of LIR motifs^40, 46^. Analysis of covalent modification by LC-MS revealed three FNIP2 peptides showing the most significant protection (553-559, 560-573 and 564-573), which span a 21-residue segment containing a single LIR motif (YVVI) at positions 567-570 (Fig. 4B, Table S5).

**Figure 4.**
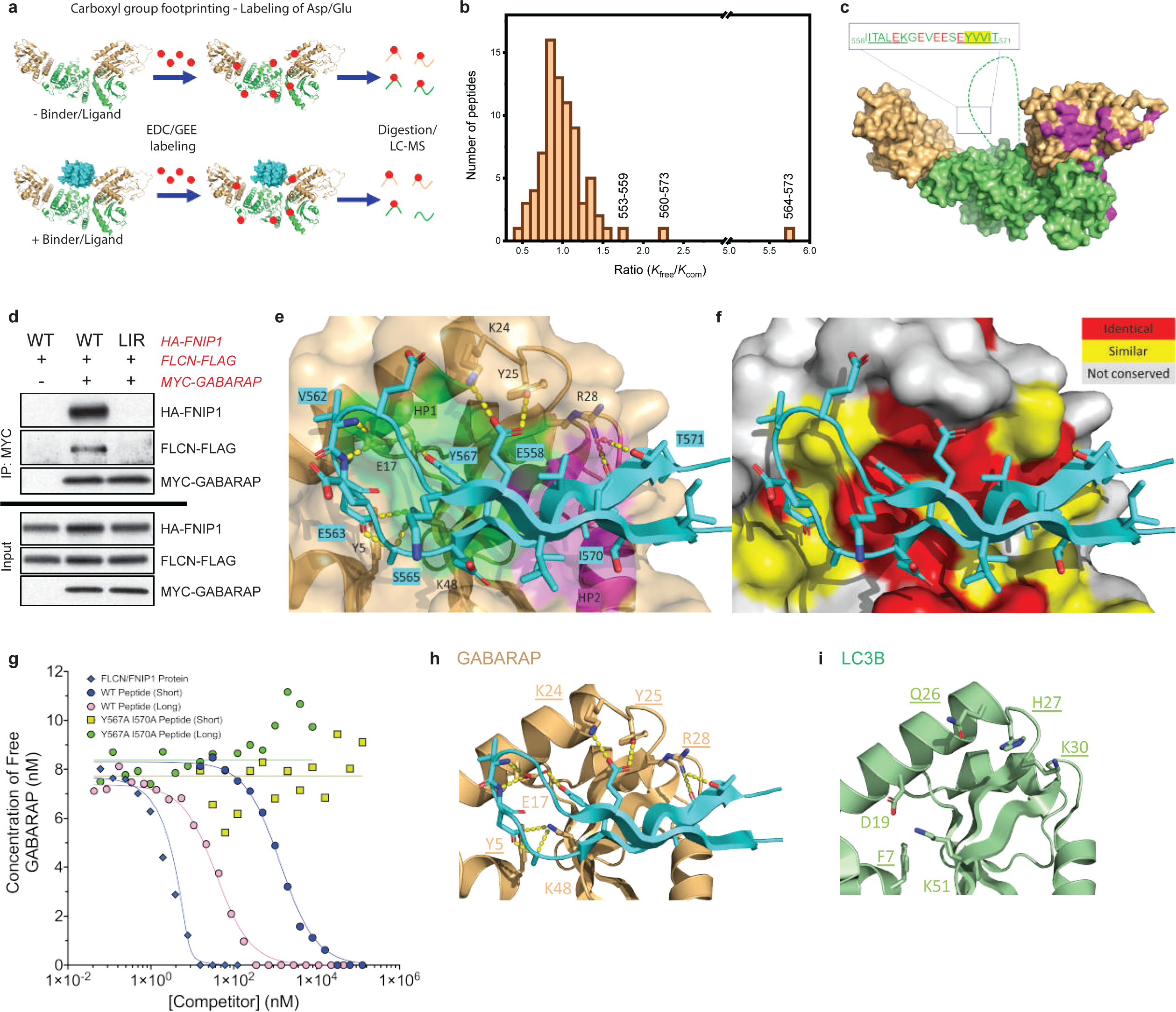
GABARAP binds the FLCN-FNIP complex through a novel LIR motif-driven interface. **(a)** Overview of chemical footprinting assay. GEE and EDC label carboxyl groups of Asp/Glu residues. **(b)** Significant protection observed for 3 overlapping peptides in FNIP2. **(c)** Location of putative LIR domain within reported FLCN-FNIP2 CryoEM structure. GABARAP binds a region located in an unresolved disordered loop, distinct from the RagGTPase binding interface (purple). **(d)** Identified LIR domain is required for GABARAP-FNIP1 interaction. HEK293T cells transiently transfected with the indicated constructs for 20hr before lysis and immunoprecipitation. LIR mutation = Y583A/V586A. **(e)** Crystal structure of FNIP2-GABARAP fusion protein. HP1 pocket shaded in green, HP2 shaded in purple. FNIP2 LIR motif forms a beta-sheet hairpin structure, with added interactions within the context of the hairpin N-terminal to the core LIR motif. **(f)** Representation of similarity between GABARAP and LC3B in the LIR docking site. **(g)** Competition SPR of immobilized FLCN/FNIP2 and GABARAP in solution with FLCN/FNIP1 protein and FNIP2 peptide competitors highlights the essential role of the LIR motif in driving the initial GABARAP-FLCN/FNIP2 interaction. Regions outside of the LIR result in stabilization and strengthening of the interaction. **(h)** Molecular interactions within the FNIP2 hairpin outside the core LIR motif. Key residues are underlined. **(i)** Sequence divergence of key underlined residues in LC3B.

Additional peptides with a lower level of protection were observed on FLCN (283-290 and 275- and FNIP2 (286-295 and 331-346) (Table S4), however these lacked an obvious LIR motif and may reflect potential conformational rearrangements upon complex formation. The putative FNIP2 LIR motif mapped to a >300-residue disordered loop within FLCN/FNIP2 from the LFC complex that was recently solved by CryoEM^20, 21^. This loop sits within the C shaped FLCN/FNIP2 cradle at a region distal to the RagA/RagC binding interface (Fig. 4C). Mutation of the orthologous LIR sequence in FNIP1 from YVLV to AVLA completely blocked GABARAP interaction with the FLCN-FNIP1 complex, whereas the FNIP1 LIR mutation did not impact its association with FLCN (Fig. 4D, Fig. S15).

We determined the crystal structure of GABARAP in complex with residues 558-576 of FNIP2 at 1.8 Å (Fig. S16, Table S1-S3). Tyr567 and Ile57 comprise the x0 and x3 positions of the 4-residue LIR motif, which occupy the canonical hydrophobic pockets 1 and 2, respectively, on GABARAP (Fig. 4E). Published co-structures of ATG8 family members often reveal additional interactions upstream and/or downstream of the LIR motif, which appear to contribute to affinity and selectivity. On the C-terminal side of the LIR, the GABARAP/FNIP2 co-structure contains a hydrogen bond between Thr571 (x_4_) and Arg28^GAB^, but no further downstream contacts in contrast to other reports showing interactions up to (x_10_)^40^. The lack of interaction on the C-terminal side is not unexpected given the lack of sequence conservation between FNIP1 and FNIP2 between x_5_ and x_10_ (Fig. S17).

On the N-terminal side of the LIR, GABARAP forms a beta-hairpin loop that contributes a number of side chain-mediated interactions (Fig. 4E, Fig. S18). In contrast to the C-terminal side, the N-terminal region is strongly conserved between FNIP1 and FNIP2. Glu558 (x-9) and Glu564 (x-3), which were protected by GEE-labeling (Fig. S17), form hydrogen bonds and salt bridges. Glu558 forms a bidentate engagement with Lys24^GAB^ and Gln25^GAB^ while Glu564 in combination with Ser565 (x-3) participate in a hydrogen bonding network that includes Tyr5^GAB^, Glu17^GAB^ and Lys48^GAB^ (Fig. 4E). Val562 occupies a shallow cleft that involves the hydrophobic portion of a trio of aliphatic residues.

To confirm the importance of the molecular interactions outside the core LIR motif, we developed a competition SPR assay using isolated FNIP2 peptides. A peptide spanning the LIR motif in FNIP2 (aa 558-576) was able to fully compete interaction of GABARAP with FLCN/FNIP2 (Fig. 4G). However, we noted a 10^4^ lower affinity of our FNIP2 LIR peptide for GABARAP (6.2 ± 3 uM) than full length FLCN/FNIP2. N-terminal extension of the FNIP2 peptide to incorporate the full stabilized hairpin structure (aa 550-576) showed markedly increased affinity for GABARAP (29 ± 2 nM K_*D*_) in competition SPR. Mutation of the candidate LIR sequence in the elongated peptide blocked competition with the GABARAP and FLCN/FNIP2, confirming the role of the LIR motif in driving the interaction between GABARAP and FLCN/FNIP2 (Fig. 4G).

It remains unclear how selectivity is established within the GABARAP and LC3B branches of the ATG8 family as universal rules have not yet been established for interaction partners. A comparison of LC3B and GABARAP shows that four of the five residues that form side chain:side chain-mediated interactions in the X-ray structure are not conserved between LC3B and GABARAP (Fig. 4H, I). These amino acid differences may contribute to the observed affinity differences of LC3B and GABARAP for the FLCN/FNIP complex (Fig. 3E).

### GABARAP sequestration of FLCN-FNIP complex to membranes is required for TFEB activation during SMAC and selective autophagy

To establish the functional requirement of the FNIP LIR domain, we reconstituted FNIP1/2 double knockout cells with WT or LIR-mutant (LIR) FNIP1. Both FNIP1-WT and FNIP1-LIR were able to rescue the constitutive TFEB activation in FNIP1/2_DKO cells as evidenced by suppression of GPNMB protein levels (Fig. 5A). TFEB activation upon acute TRPML1 stimulation was blocked specifically in cells expressing FNIP1-LIR, whereas TFEB activation in response to nutrient starvation was not impacted by FNIP1-LIR (Fig. 5A, B). Upon chronic treatment with the TRPML1 agonist, the functional TFEB transcriptional response can be measured by protein levels of the target gene *GPNMB*. GPNMB expression was completely blocked in FNIP1-LIR mutant cells (Fig. 5C). GPNMB protein levels were also largely suppressed upon AZD8055 treatment despite robust TFEB activation. This highlights how concurrent inhibition of protein translation may minimize the effective scope of the TFEB transcriptional activation (Fig. 5C). ATG8 conjugation was not altered by modulation of FNIP1 (Fig. 5A, C). Membrane sequestration of the FLCN-FNIP complex also required the identified FNIP1-LIR domain, confirming our hypothesis that GABARAP binding to the FLCN-FNIP complex is responsible for its relocalization (Fig. 5D). Other inducers of SMAC, such as the ionophore monensin, could induce FLCN-FNIP1 complex sequestration independently of TRPML1 (Fig. S19A) and regulated TFEB activation in a FNIP1-LIR domain dependent manner (Fig. S19B, C). This suggests that perturbation of lysosomal ion homeostasis, rather then TRPML1 activation specifically, serves as a trigger for GABARAP-dependent FLCN-FNIP relocalization.

**Figure 5.**
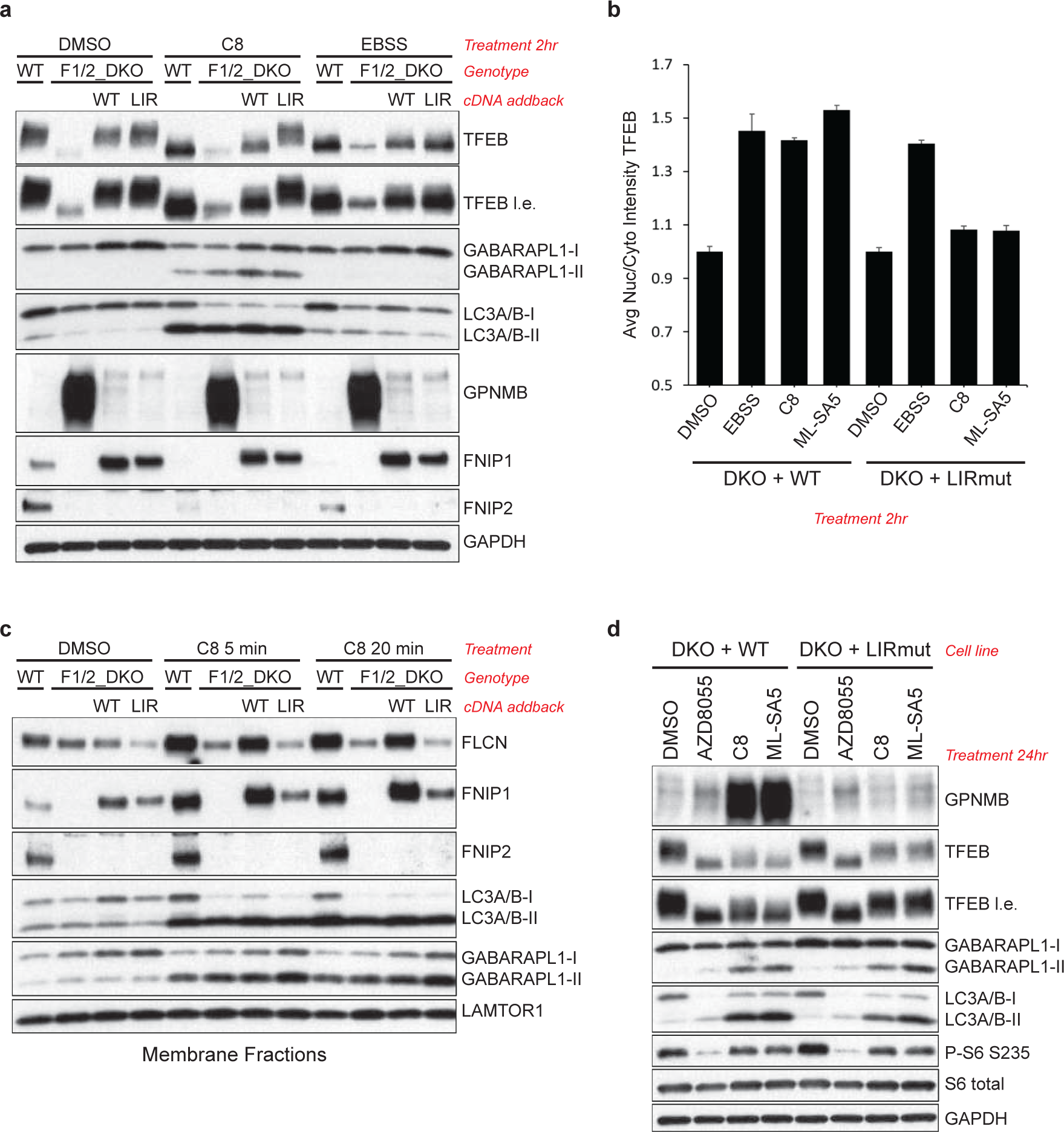
GABARAP-dependent sequestration of FLCN-FNIP complex is required to activate TFEB upon disruption of endolysosomal ion balance. **(a)** Reconstitution of FNIP1/2 double knockout (DKO) cells with either WT or LIR (LIR-mutant Y583A/V586A) FNIP1 reveals functional requirement of GABARAP interaction for TRPML1 agonist, but not EBSS, activation of TFEB. Western blot analysis of FNIP1 allele series treated with the indicated stimuli. l.e = long exposure. **(b)** Quantification of TFEB nuclear localization in WT or LIR expressing FNIP1/2_DKO HeLa cells treated with the indicated stimuli. Analysis was performed using high content imaging. Mean ± SD. Minimum of 1500 cells quantified per condition. C8 = 2uM. ML-SA5 = 1uM. **(c)** Functional TFEB response upon prolonged TRPML1 activation requires FNIP1 LIR domain. Western blot analysis of WT or LIR expressing FNIP1/2_DKO HeLa cells treated with the indicated stimuli. GPNMB is a validated TFEB/TFE3 transcriptional target. **(d)** Western blot analysis of membrane fractions from FNIP1 allele series after acute treatment with TRPML1 agonist for the indicated timepoints.

It is intriguing to consider that any instance where GABARAP proteins are conjugated to subcellular membranes might result in TFEB activation via the high affinity sequestration of FLCN/FNIP. Thus, we examined distinct forms of selective autophagy, mitophagy and xenophagy. It has been shown that TFEB activation occurs during parkin-dependent mitophagy^47^, and we found that this required GABARAP proteins (Fig. 6A, B, Fig. S15).

**Figure 6.**
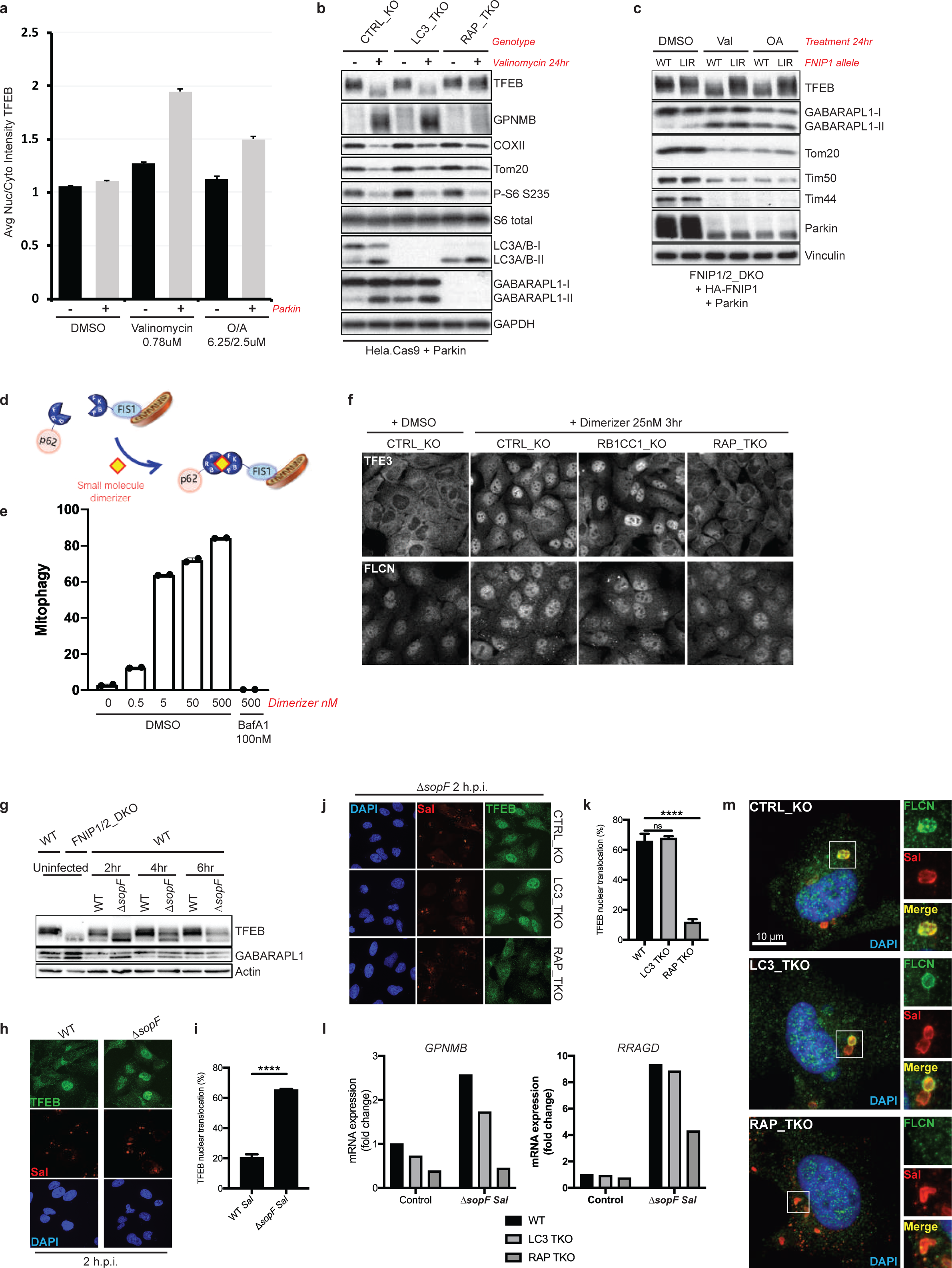
GABARAP regulates TFEB activation through FLCN relocalization during selective autophagy. **(a)** Parkin is required for robust TFEB activation upon stimulation of mitophagy. HeLa.Cas9 or HeLa.Cas9+ Parkin cells were treated with the indicated compounds for 4 hr and analyzed by immunofluorescence for endogenous TFEB. **(b)** GABARAPs, but not LC3s, are specifically required for TFEB transcriptional activity during mitophagy. HeLa cells expressing Parkin and CRISPR KO for the indicated ATG8 family members were treated with 0.78 uM valinomycin for 24 hr. **(c)** FNIP LIR motif is required for robust TFEB activation upon mitochondrial dysfunction. Cells of the indicated genotype were treated with mitophagy inducers for 24 hr. **(d)** Overview of proximity-based mitophagy induction model. Recruitment of p62 to mitochondria results in mitophagy independent of chemical disruption of mitochondrial function. **(e)** Quantification of mitophagy efficiency using the mKeima flow cytometry assay. U2OS cells expressing mKeima, FRB-p62, and FKBP-FIS1 were clonally isolated. Mitophagy = percentage of cells in the Keima_acidic_ quadrant. Keima_acidic_ signal can be fully blocked with BafA1 cotreatment. **(f)** Analysis of TFE3 and FLCN subcellular localization upon dimerizer induced mitophagy in U2OS cells. Cells of the indicated genotype were stimulated with 25nM AP21967 (dimerizer) for 3 hr. TFE3 nuclear localization and FLCN punctate structures are seen in CTRL and RB1CC1_KO (autophagy-deficient) cells but not in GABARAP_TKO cells. **(g)** Analysis of TFEB mobility shift upon challenge with WT or ΔsopF *Salmonella*. HeLa cells were infected for 30 min with the indicated strain and lysates taken an the indicated timepoints post infection. **(h)** Immunofluorescence analysis of nuclear TFEB accumulation upon *Salmonella* infection. Cells were infected with the indicated strain as above and analyzed at 2 hours post infection (h.p.i.). **(i)** Quantification of TFEB nuclear localization. A minimum of 100 cells were quantified per condition. **(j)** GABARAPs are specifically required for TFEB activation upon infection. Cells of the indicated genotype were infected with ΔsopF *Salmonella* as above and analyzed at 2 h.p.i.. **(k)** Quantification of TFEB nuclear localization. A minimum of 100 cells were quantified per condition. **(l)** Analysis of TFEB transcriptional activity in cells of the indicated genotype at 10 h.p.i with ΔsopF *Salmonella*. GPNMB and RRAGD were used as core TFEB target genes. **(m)** Analysis of FLCN recruitment to *Salmonella* vacuoles. Deletion of GABARAP proteins, but not LC3 family members, blocks the relocalization of FLCN.

Furthermore, TFEB activation was defective in cells stably expressing LIR-mutant FNIP1, confirming that GABARAP-dependent relocalization of FLCN to mitochondria mechanistically links TFEB activation to mitophagy (Fig. 6C). Interestingly, an earlier study observed that FLCN and FNIP could localize to mitochondria upon depolarization^48^ and the TFEB activation mechanism revealed in the current study explains the relevance of this. Importantly, using a proximity-regulated mitophagy system, where mitophagy is measured using the Keima fluorescence shift assay^49^ (Fig. 6D, E) we confirmed the GABARAP-dependence of TFE3 translocation and FLCN redistribution independent of using mitochondrial uncouplers (Fig. 6F).

Finally, we used a *Salmonella* infection model of xenophagy to determine if TFEB activation was regulated by GABARAP mediated FLCN sequestration. A portion of *Salmonella enterica* serovar Typhimurium (*S.* Typhimurium) are rapidly targeted by the autophagy machinery and become decorated with ATG8 homologs^50^. It was recently discovered that *S*. Typhimurium antagonize the ATG8 response through the bacterial effector SopF^25^. We hypothesized that if ATG8 proteins were involved in TFEB activation, Δ*sopF S.* Typhimurium would show a greater TFEB activation than WT due to increased ATG8 conjugation. Indeed, Δ*sopF S.* Typhimurium produced a robust activation of TFEB that occurred in a higher percentage of cells for a longer duration of time post-infection (Fig. 6G-I). Importantly, TFEB activation was blunted by deletion of GABARAP family members (RAP_TKO), but was not impacted by knockout of LC3 isoforms (LC3_TKO) (Fig. 6J-L). We next examined the localization of FLCN and found a striking relocalization of FLCN to coat the *S*. Typhimurium *Salmonella*-containing vacuole membrane (Fig. 6M). This relocalization required GABARAP proteins, indicating that GABARAP-dependent sequestration of FLCN to the *Salmonella* vacuoles results in TFEB activation upon infection (Fig. 6M). Taken together, the mitophagy and xenophagy examples of selective autophagy highlight that GABARAP-dependent sequestration of FLCN to distinct cellular membranes may serve as a universal mechanism to couple activation of TFE3/TFEB transcription factors to the initiation of selective autophagy.

## Discussion

We describe a previously unrecognized molecular mechanism orchestrated by GABARAP proteins independent from their role in substrate degradation and vesicle maturation. Changes in endolysosomal ion levels and induction of SMAC helped uncover a novel regulatory input to TFEB nuclear localization that is independent of nutrient status and requires the ATG5-ATG12-ATG16L1 conjugation machinery. GABARAP-dependent membrane sequestration of the FLCN-FNIP complex uncouples its regulation of RagC/D revealing a new paradigm for TFEB activation distinct from LFC formation during nutrient starvation. Moreover, GABARAP-dependent TFEB activation is permissive with mTORC1 activity offering new insights into strategies to enhance lysosomal biogenesis. Given the conjugation of ATG8 proteins to double membrane autophagosomes^51^, it is logical that GABARAPs would play a critical role in coordinating the activation of TFEB during the early stage of autophagy. Indeed, the GABARAP-FLCN-FNIP axis was required for TFEB activation during mitophagy and xenophagy suggesting that this newly identified mechanism can broadly serve to coordinate lysosomal capacity (Fig. 7). Conjugation of GABARAP and sequestration of FLCN-FNIP to the forming autophagosome can be viewed as a stage-gate ensuring tight coordination of autophagy with lysosomal capacity.

**Figure 7.**
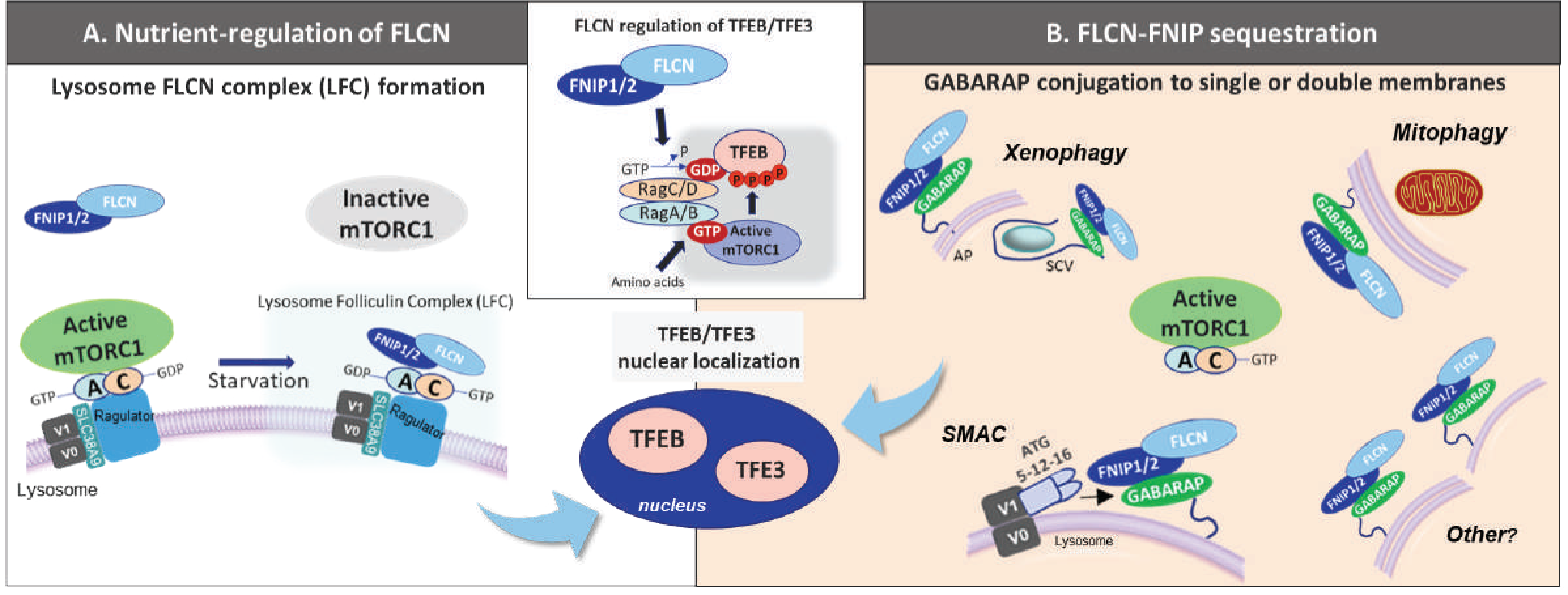
GABARAP-dependent membrane sequestration of the FLCN-FNIP complex represents a TFEB activation paradigm distinct from nutrient starvation. The FLCN-FNIP GAP complex critically regulates the mTOR-dependent phosphorylation and cytosolic retention of the TFEB/TFE3 transcription factors by promoting the GDP-bound state of RagC/D. GDP-bound RagC/D directly binds to and presents TFEB/TFE3 as a substrate to mTOR (center inset), as described previously. During nutrient starvation **(A)**, recruitment of FLCN-FNIP to the lysosomal membrane helps form the lysosomal folliculin complex (LFC), which has reduced GAP activity towards RagC/D. This is coincident with mTORC1 inhibition. Independently of LFC formation, GABARAP proteins bind directly to the FLCN-FNIP complex and sequester it at diverse intracellular membranes **(B)**. This membrane recruitment is required for TFEB activation in response to endolysosomal ion disruption (SMAC) and forms of selective autophagy (xenophagy and mitophagy). This suggests that FLCN-FNIP regulates cytosolic RagC-GTP and its sequestration on intracellular membranes reduces access to this substrate, allowing for nuclear retention of TFEB/TFE3 due to impaired Rag binding. Unlike (A), this novel TFEB activation pathway is permissive with mTORC1 activity. Subcellular redistribution of the FLCN-FNIP complex to both single and double membranes serves to broadly coordinate lysosomal capacity with homeostasis and perturbations within the endolysosomal network.

Germline mutations in *FLCN* underlie the FLCN-FNIP complex loss of function phenotype and TFEB-dependency in Birt-Hogg-Dube syndrome^22, 52^, a rare disorder that predisposes patients to kidney tumors. Interestingly, Ras-driven pancreatic adenocarcinoma cells (PDAC) show constitutive nuclear localization of TFEB/TFE3 and notable colocalization of LC3 with LAMP2-positive lysosomes^53, 54^. Moreover, elevated activity and permeability of lysosomes has been noted in PDAC^54^ and our study suggests a mechanism to explain TFEB nuclear localization despite nutrient replete and mTOR-active conditions^53^. Understanding the involvement of FLCN-FNIP membrane sequestration in this setting and whether oncogenic signals might take advantage of this mechanism to drive TFEB-dependent tumor growth may offer new therapeutic opportunities for lysosome-dependent tumors.

Linking TFEB activation to GABARAP membrane conjugation allows for sensitive detection of not only autophagy initiation and flux, but also dysfunction within the endolysosomal pathway, possibly as part of a host-pathogen response. Pathogens have evolved virulence factors to inhibit and evade SMAC, for example SopF of *S.* Typhimurium^25^, CpsA of *M. tuberculosis*^55^, and RavZ of *Legionella*^56^. Recently, it has been proposed that disruption of the phagosomal ion gradient triggered ATG8 modification of the Δ*sopF S.* Typhimurium-containing vacuole and that this precedes vacuole rupture and xenophagy^25^. Induction of SMAC could serve to couple TFEB-dependent transcription of cytoprotective/antimicrobial genes^57^ and lysosomal biogenesis to limit pathogen infection. While SMAC induction does result in the conjugation of both LC3 and GABARAP homologs to target membranes, our study highlights that different ATG8s serve distinct functions. While LC3 is proposed to regulate vesicle maturation and fusion with lysosomes^58, 59^, the primary role of GABARAP may be to coordinate lysosomal capacity to accommodate increased rates of lysosomal delivery.

Our data highlights the importance of the RagC nucleotide state in the regulation of TFEB by mTOR^22^. mTOR has been shown to mediate the nuclear export of TFEB/TFE3 transcription factors, with phosphorylation promoting cytosolic retention^60^. In the absence of FLCN-FNIP activity, the regulation of nuclear export is impaired due to a lack of mTOR access to TFEB/TFE3 as substrates, resulting in nuclear retention rather than active translocation of these transcription factors^22^. The GABARAP-dependent sequestration of FLCN-FNIP represents a new paradigm for the control of the RagC/D nucleotide state, previously thought to solely be regulated by nutrient levels. The contribution of this new mechanism to both the basal and induced adaptive TFEB/TFE3 responses will be interesting to test in future work. Interestingly, mice expressing a C-terminal ATG16L1 truncation, thus defective in SMAC, show an Alzheimer’s disease (AD) phenotype^61^ and deficiencies in lysosomal biogenesis/homeostasis are well-characterized in AD and other neurodegenerative disorders^62^. Further understanding the regulation of this GABARAP-dependent sequestration of FLCN-FNIP, and whether a homeostatic defect in TFEB/TFE3 activation contributes to neurodegeneration *in vivo* will provide additional insights into therapeutic opportunities.

## Materials and Methods

### Antibodies

Antibodies used in this study were as following: ATG16L1 (8089, human), Phospho-ATG14 S29 (92340), ATG14 (96752), Phospho-Beclin S30 (54101), FIP200 (12436), FLCN (3697), GABARAPL1 (26632), GABARAPL2 (14256), GAPDH (5174, 1:10000 for WB), DYKDDDDK tag (14793), HA tag (3724), myc tag (2278), LC3A/B (12741), LC3B (3868), LAMTOR1 (8975), LAMP1 (15665, 1:1000 for IF), NFAT1 (5861, 1:250 for IF), NPRL2 (37344), Phospho-S6K (9234), S6K (2708), Phospho-S6 S235/236 (4858, 1:3000 for WB), S6 (2217, 1:5000 for WB), TAX1BP1 (5105), TFEB (4240), TFEB (37785, 1:200 for IF), and Phospho-ULK S757 (14202) were from Cell Signaling Technologies. Mouse monoclonal anti-S. Typhimurium LPS (clone 1E6, ab8274) and FNIP1 (ab134969) were from Abcam. TFE3 (HPA023881) was from Millipore Sigma. p62 (GP62-C) was from Progen. Galectin-3 (sc-23938) was from Santa Cruz Biotechnology. TFEB (A303-673A, 1:200 for IF in murine cells) was from Bethyl Laboratories. All antibodies were used at a 1:1000 dilution for western blotting unless otherwise noted.

### Generation of knockout cell lines with CRISPR/Cas9

HeLa cells or U2OS cells were made to stably express Cas9 through lentiviral transduction (vector Cat# SVC9-PS-Hygro, Cellecta). Knockout cell lines were generated as pooled populations following subsequent lentiviral transduction with gRNA sequences as indicated (vector Cat# SVCRU6UP-L, Cellecta). Pooled populations were selected for 3 days with puromycin (2ug/ml, Life Technologies) and used for experiments 7-9 days post transduction with gRNA. Clones were isolated for ATG16L1_KO to use for reconstitution experiments. gRNA sequences were as follows (5’-3’):

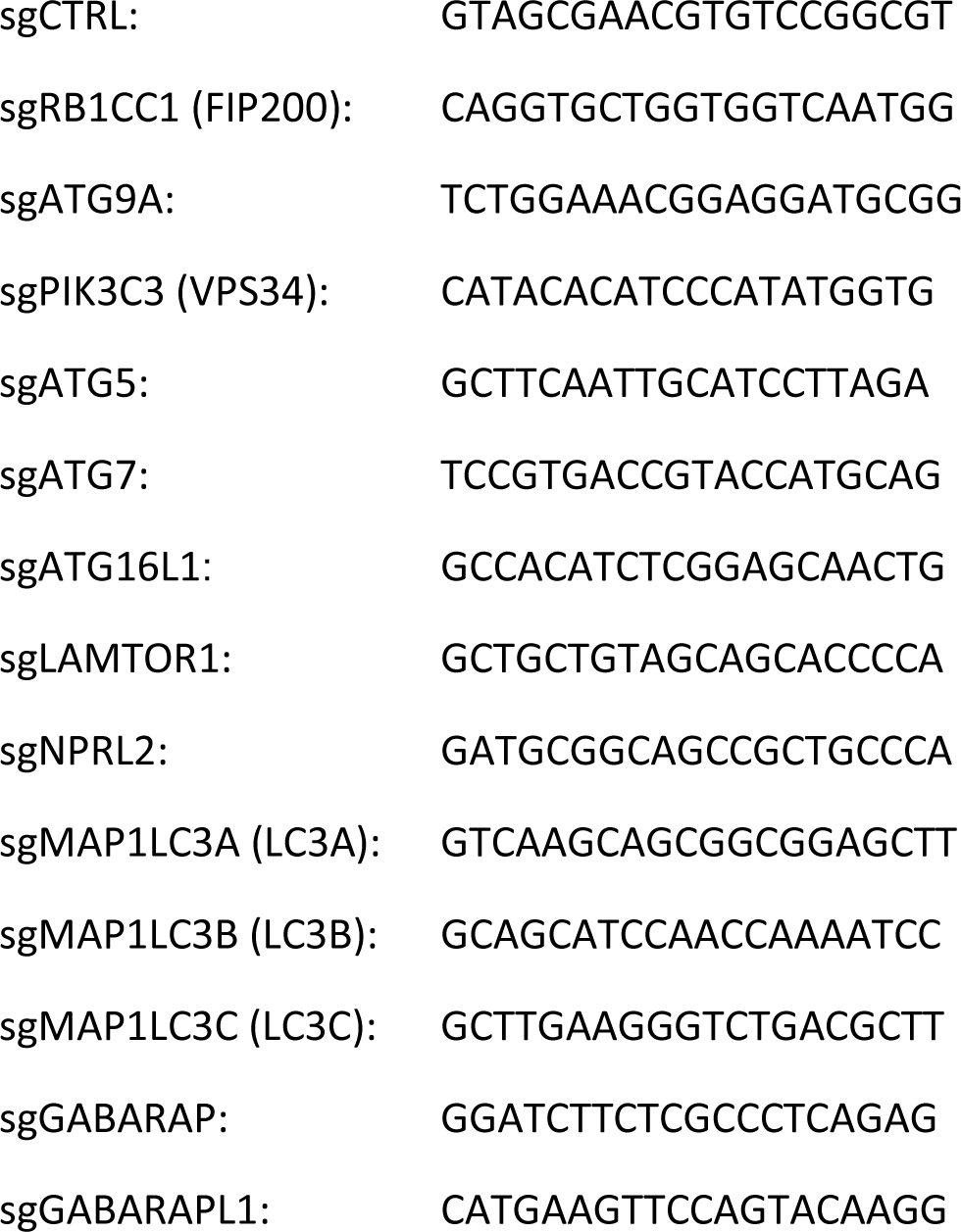
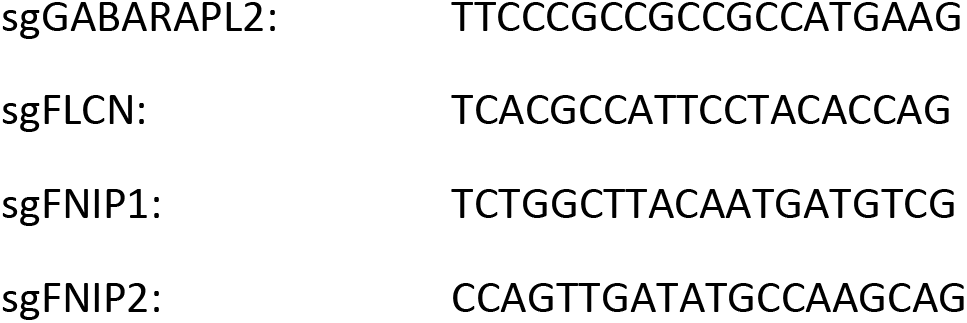

### cDNA expression constructs

Wild type and K490A mutant mouse ATG16L1 were cloned into pBabe-Puro-Flag-S-tag plasmids as previously described^26^. pBabe-Blast-GFP-LC3A has been previously described^5^.

The following constructs were generated for use in this study:

| Insert | Epitope Tag | Terminus | Expression vector |
| --- | --- | --- | --- |
| Human FNIP1 WT | 3X HA | N | pcDNA-DEST40 |
| Human FLCN WT | FLAG | C | pcDNA-DEST40<br>Tet Lenti |
| Human lyso-FLCN<br>(Nterm 39aa of<br>LAMTOR1 fusion | FLAG | C | Tet Lenti |
| Human GABARAP WT | myc | N | pcDNA-DEST40 |
| Human GABARAP<br>LBMmut | myc | N | pcDNA-DEST40 |
| Human ATG16L1 WT | FLAG | N | Tet Lenti |
| Human ATG16L1 ΔCTD | FLAG | N | Tet Lenti |
| Human ATG16L1 ΔFBD | FLAG | N | Tet Lenti |
| Human ATG16L1 F467A | FLAG | N | Tet Lenti |
| Human ATG16L1 K490A | FLAG | N | Tet Lenti |
| S. Typhimurium SopF | myc | N | Tet-Lenti |
| Human LAMP1 | RFP | C | pBabe |
| Human FNIP1 WT | 3XHA | N | Tet Lenti |
| Human FNIP1<br>Y583A/V586A | 3XHA | N | Tet Lenti |
| Human TMEM192 | 3XHA | C | Tet Lenti |

cDNA constructs with the indicated epitope tags were synthesized (Genscript, USA) and provided as entry clones. Gateway recombination was used to shuttle cassettes into pcDNA-DEST40 (Life Technologies) or a lentiviral vector allowing tetracycline inducible expression referred to as Tet-Lenti (synthesized by Genscript, USA).

### Cell Culture

Cell lines used in this study were U2OS, HeLa, and RAW264.7 and were obtained from the American Type Culture Collection (ATCC). HEK293FT were obtained from ThermoFisher Scientific. Cell lines were verified to be mycoplasma-free by routine testing. All cells were cultured in a humidified incubator at 37°C and 5% CO_2_. Cell culture reagents were obtained from Invitrogen unless otherwise specified. Cells were grown in Dulbecco’s Modified Eagle’s Medium (DMEM) supplemented with 10% fetal bovine serum and 1% penicillin/streptomycin.

### Reagents

Bafilomycin A1, PIK-III, and AZD8055 were purchased from Selleckchem. ML-SA1 and MK6-83 were purchased from Tocris. Monensin, nigericin, salinomycin, valinomycin, and LLoMe were purchased from Sigma Aldrich. C8 is available for purchase through Chemshuttle (Cat# 187417).

### Viral production and transduction

For lentiviral production of CRISPR gRNA or Cas9 virus and cDNA overexpression virus, 8 × 10^5^ 293FT cells were plated in 6-well plates. The next day, cells were transfected with lentiviral packaging mix (1 ug psPAX2 and 0.25 ug VSV-G) along with 1.5 ug of the lentiviral backbone using Lipofectamine 2000 (ThermoFisher). Supernatant was removed from 293FT cells after 48 hr, centrifuged at 2000 rpm for 5 min and then syringe filtered using a 0.45 um filter (Millipore). Polybrene was then added to a final concentration of 8 ug/ml and target cells were infected overnight. Cells were then allowed to recover for 24 hr in DMEM/10% FBS before being selected with 1 mg/ml neomycin (G418:Geneticin, ThermoFisher), 2ug/ml puromycin (ThermoFisher), or 500 ug/ml Hygromycin B (ThermoFisher) for 72 hr. Retroviral infection was performed using centrifugation. Stable populations were selected with puromycin (2 mg/ml) or blasticidin (10 mg/ml) for 3-5 days.

### Cell lysis and western blotting

For preparation of total cell lysates, cells were lysed in RIPA buffer (#9806, Cell Signaling Technology) supplemented with sodium dodecyl sulfate (SDS, Boston BioProducts) to 1% final concentration, and protease inhibitor tablets (Complete EDTA-free, Roche). Lysates were homogenized by sequential passaging through Qiashredder columns (Qiagen), and protein levels were quantified by Lowry DC protein assay (Bio-Rad). Proteins were denatured in 6X Laemmli SDS loading buffer (Boston BioProducts) at 100°C for 5 min.

For preparation of membrane fractions, 1.5 × 10^6^ cells were plated the day before in 6cm tissue culture dishes (BD Falcon). Cellular fractions were prepared using the MEM-PER kit (ThermoFisher) according to the manufacturer’s protocol. Protein levels were quantified by Lowry DC protein assay (Bio-Rad) and denatured in 6X Laemmli SDS loading buffer (Boston BioProducts).

For western blotting, equivalent amounts of total proteins were separated on Tris-Glycine TGx SDS-PAGE gels (Bio-Rad). Proteins were transferred to nitrocellulose using standard methods and membranes were blocked in 5% non-fat dry milk (Cell Signaling Technology) in TBS with 0.2% Tween-20 (Boston BioProducts). Primary antibodies were diluted in 5% bovine serum albumin (BSA, Cell Signaling Technology) in TBS with 0.2% Tween-20 and were incubated with membranes at 4°C overnight. HRP-conjugated secondary antibodies were diluted in blocking solution (1:20,000, ThermoFisher) and incubated with membranes at room temperature for 1 h. Western blots were developed using West PicoPLUS Super Signal ECL reagents (Pierce) and film (GEHealthcare).

### Immunoprecipitation

For immunoprecipitations, cells were lysed in IP CHAPS lysis buffer: 0.3% CHAPS, 10 mM β-glycerol phosphate, 10 mM pyrophosphate, 40 mM Hepes pH 7.4, 2.5 mM MgCl_2_, supplemented with protease inhibitor tablets (Roche) and Calyculin A (Cell Signaling Technology). Lysates were clarified by centrifugation and equilibrated as described above. For FLAG IP, lysates were incubated with anti-M2 FLAG-conjugated agarose beads (Sigma-Aldrich) at 10 ul bed volume per 1 mg of protein and incubated for 1 hour at 4°C with gentle rocking. For MYC IP, lysates were incubated with 10 ul bed volume per 1 mg protein of anti-myc 9E10-conjugated agarose beads (Sigma Aldrich). Beads were then centrifuged and washed 3X with lysis buffer supplemented with 300mM NaCl. Immunoprecipitate was eluted by addition of 6X Laemmli SDS loading buffer at 100°C for 5 min.

LysoIP was performed as described previously^63^. Briefly, U2OS cells stably expressing 3XHA-TMEM192 were washed twice with ice-cold PBS and then scraped into 1ml of ice-cold LysoIP buffer (136 mM KCl, 10 mM KH_2_PO_4_, pH 7.25 in Optima LC/MS water). 100ul of cell suspension was reserved for input sample. Cells were then homogenized using 35 strokes of a dounce homogenizer followed by centrifugation for 10 minutes at 1500 x g in a 4°C centrifuge. Clarified homogenates were then incubated with 50ul prewashed anti-HA magnetic beads for 30 minutes with constant rotation at 4°C. Beads were then washed 4X with LysoIP washing buffer (136 mM KCl, 10 mM KH_2_PO_4_, pH 7.25 in Optima LC/MS water, 300mM NaCl). Both input and immunoprecipitated samples were then lysed with LysoIP lysis buffer (40 mM HEPES pH 7.4, 1% Triton X-100, 10 mM β-glycerol phosphate, 10 mM pyrophosphate, 2.5 mM MgCl_2_ and complete EDTA-free Protease Inhibitor Cocktail (Roche)). 6X Laemmli SDS loading buffer was added and samples placed at 100°C for 5 min.

### Immunofluorescence and high content image analysis

Following indicated treatments, GFP-LC3 LAMP1-RFP expressing cells were fixed with ice cold methanol for 3 minutes at -20°C. Cells were washed in PBS and image acquisition was performed using a Confocal Zeiss LSM 780 microscope (Carl Zeiss Ltd) equipped with a 40x oil immersion 1.40 numerical aperture (NA) objective using Zen software (Carl Zeiss Ltd).

For quantification, the number of GFP-LC3 puncta were counted for >20 cells across multiple fields of view. For co-localization quantification, GFP-LC3 puncta were assessed for LAMP1-RFP.

For LC3 and LAMP1 staining in primary BMDMs, cells were plated on 18 mm coverslips. The next day cells were treated as indicated and cells fixed in ice cold methanol as above. Cells were then blocked in PBS + 5% BSA for 1 hour before addition of primary antibodies (anti-LC3A/B, CST #4108, 1:100; anti-LAMP1, BD #555798, 1:100) diluted in blocking buffer overnight at 4°C. Cells were washed and incubated with fluorescent secondary antibodies in blocking buffer for 1 hour at room temperature. Cells were washed in PBS incubated with DAPI and mounted on glass slides using Prolong anti-fade reagent (Life Technologies).

For endogenous TFEB staining in mouse macrophage, cells were fixed in 3.7% formaldehyde for 15 minutes at room temperature, washed in PBS and permeabilized in 0.2% triton/PBS for 5 mins. Cells were then processed as above for primary (anti-TFEB, Bethyl Laboratories, #A303-673A, 1:200) and secondary antibodies. Images were acquired using a Confocal Zeiss LSM 780 microscope (Carl Zeiss Ltd) equipped with a 40x oil immersion 1.40 numerical aperture (NA) objective using Zen software (Carl Zeiss Ltd). Analysis was performed using Image J. For nuclear cytosol quantification, the ratio of fluorescent intensity of TFEB within the DAPI was mask versus the cytosol of 30 cells across 2 independent experiments were measured.

For high content image acquisition, cells were plated in 96-well glass bottom, black wall plates (Greiner, #655892) or 384-well polystyrene, black-wall plates (Greiner, #781091) and grown overnight to 70% confluency. Treatments were performed as indicated. Cells were fixed for 10 min in either -20°C methanol or 4% paraformaldehyde (Electron Microscopy Sciences). Cells were blocked and permeabilized in a solution containing 1:1 Odyssey blocking buffer (LiCor)/PBS (Invitrogen) with 0.1% Triton X-100 (Sigma) and 1% normal goat serum (Invitrogen) for 1 h at room temperature. Primary antibodies were added overnight at 4°C in blocking buffer described above. After washing plates with PBS using an EL-406 plate washer (BioTek), secondary Alexa-conjugated antibodies (Life Technologies) were diluted 1:1,000 in blocking solution supplemented with DAPI and applied for 1 h at room temperature. Cells were then washed again in PBS as described and imaged using an INCELL 6500 high-content imager (GE Healthcare). Images were analyzed using the GE InCarta software.

For imaging in the context of *Salmonella* infection, cells were fixed with 4% paraformaldehyde in PBS at 37 °C for 10 min. Immunostaining was performed as previously described^64^. Cells were imaged using a Quorum spinning disk microscope with a 63Å∼ oil immersion objective (Leica DMIRE2 inverted fluorescence microscope equipped with a Hamamatsu Back-Thinned EM-CCD camera or Hamamatsu CMOS FL-400 camera, spinning disk confocal scan head) and Volocity 6.3 acquisition software (Improvision). Confocal z-stacks of 0.3 μm were acquired and images were analysed with Volocity 6.3 software.

### Correlative FIB-SEM

Cells were seeded in 35-mm glass-bottom dishes (MatTek Corp., USA, #P35G-2-14-CGRD). They were fixed with 4% formaldehyde (TAAB F017, 16% w/v solution formaldehyde-methanol free) in 0.1 M phosphate buffer pH 7.4 (PB) for 30 min at 4 °C. They were washed in PB and imaged with a 40x/1.4 NA objective on an inverted confocal microscope (Zeiss LSM780). Further fixation was carried out with 2% formaldehyde and 2.5% glutaraldehyde (TAAB G011/2, 25% solution glutaraldehyde) in PB for 2 hours, prior to further processing.

Samples were embedded using a protocol as described previously^65, 66^. The cells were washed in PB five times and post-fixed in 1% osmium tetroxide (Agar Scientific, R1023, 4% solution osmium tetroxide) and 1.5% potassium ferrocyanide (v/v) (SIGMA ALDRICH, P3289-100G, potassium hexacyanoferrate (II) trihydrate) for 1 hour on ice. And then dehydrated and embedded in Hard-Plus Resin812 (EMS, #14115). The samples were polymerized for 72 h at 60 °C. The coverslip was removed from the resin by dipping the block into liquid nitrogen. After locating the region of interest (ROI) on the block surface using the imprint of the grid, the block was cut to fit on an aluminium stub using a hacksaw, and trimmed with a razorblade. The block/stub was then coated with 20nm Pt using a Safematic CCU-010 sputter coater (Labtech) to create a conductive surface.

The block/stub was placed in the chamber of a Zeiss 550 CrossBeam FIB SEM and the surface imaged using the electron beam at 10 kV to locate the grid and underlying cells. Once the ROI had been identified, Atlas software (Fibics) was used to operate the system. A trench was cut into the resin to expose the target cell and serial SEM images were acquired with 7 nm isotropic resolution using a 1.5 kV electron beam. For 3D-image analysis, image stacks were processed using Atlas software and viewed using ImageJ.

### RNA isolation and RNAseq analysis

#### RNA isolation

Total RNA was prepared from cells treated with DMSO or 2uM C8 for 24 hours using Trizol extraction and RNAeasy Mini Kit (Qiagen). A total of 2ug of RNA with a RIN score of >9.8 was submitted for RNAseq analysis.

#### Library preparation, HiSeq sequencing and Analysis

RNA sequencing libraries were prepared using the NEBNext Ultra RNA Library Prep Kit for Illumina using manufacturer’s instructions (NEB, MA). mRNAs were enriched with Oligod(T) beads, and the enriched mRNAs fragmented at 94°C (15 minutes). This was followed by first strand and second strand cDNA synthesis. cDNA fragments were end-repaired and adenylated at 3’ends. Universal adapters were then ligated to cDNA fragments, followed by index addition and library enrichment by PCR with limited cycles. The sequencing library and RNA samples for RNAseq were quantified using Qubit 2.0 Fluorometer (Life Technologies, CA) and RNA integrity checked using Agilent TapeStation 4200 (Agilent Technologies, CA).

The sequencing libraries were clustered on a single lane of a flowcell on the Illumina HiSeq 4000 system according to manufacturer’s instructions. The samples were sequenced using a 2x150bp Paired End (PE) configuration. Image analysis and base calling were conducted by the HiSeq Control Software (HCS). Raw sequence data (.bcl files) generated from Illumina HiSeq was converted into fastq files and de-multiplexed using Illumina’s bcl2fastq 2.17 software. One mismatch was allowed for index sequence identification. Sequence reads were mapped to the Homo sapiens reference genome version GRCh38 available on ENSEMBL using the STAR aligner v.2.5.2b. Unique gene hit counts were calculated by using feature Counts from the Subread package v.1.5.2. Only unique reads that fell within exon regions were counted.

The count data was normalized by the trimmed mean of M-values normalization (TMM) method, followed by variance estimation and applying generalized linear models (GLMs), utilizing functions from empirical analysis of digital gene expression^67^ to identify differentially expressed genes as described previously^68, 69^. Factorial designs were incorporated into the analysis by fitting these linear models with the coefficient for each of the factor combinations and then simultaneously extracting contrasts for the respective ‘differential-of-differential’ analysis in the two experimental dimensions (C8 stimulation and genotype status: ATG16L1KO and WT). The associated p-values were adjusted to control the false discovery rate in multiple testing, using the Benjamini and Hochberg’s (BH) method (BH-adjusted p<0.05).

Pathway and biological process enrichment analysis were performed as previously described^69, 70^. Briefly, data were interrogated from KEGG pathways and gene ontology biological processes. Each module or category was assessed for statistical enrichment or over-representation among differentially expressed genes relative to their representation in the global set of genes in the genome. P-values were computed using the hypergeometric test.

### Quantitative PCR with reverse transcription analysis

RNA extractions were performed using the RNeasy Mini Kit (QIAGEN), and complementary DNA was subsequently generated using the iScript cDNA Synthesis Kit (Bio-Rad). Quantitative PCR analysis was performed on the QuantStudio 3 Real-time PCR system (ThermoFisher) using the SsoFast EvaGreen Supermix Kit (Bio-Rad) and the following primer sets: GPNMB human (5’ GCCTTTAAGGATGGCAAACA 3’ and 5’ TGCACGGTTGAGAAAGACAC 3’), RRAGD human (5’ TCCGGTGGATATGCAAACCT 3’ and 5’ ACAAAGCAAACGAGAGCCAG 3’) and GAPDH human (5’ GGAGCGAGATCCCTCCAAAAT 3’ and 5’ GGCTGTTGTCATACTTCTCATGG 3’). Data were normalized to that of the housekeeping gene GAPDH.

### Lysotracker staining

U2OS.Cas9 cells expressing a control gRNA or knocked out for ATG16L1 were treated for 24 hours with 2uM C8. Cells were then washed and incubated live for 20 min with 25nM Lysotracker Red DND-99 (ThermoFisher) and Hoecsht 33342 (ThermoFisher) diluted in warmed imaging buffer (20 mM HEPES (pH 7.4), 140 mM NaCl, 2.5 mM KCl, 1.8 mM CaCl_2_, 1 mM MgCl_2_, 10 mM D-glucose, and 5% v/v FBS). Staining solution was removed and cells were incubated in imaging buffer for an additional 30 min before image acquisition on the INCELL 6500. Images were analyzed using the GE InCarta software.

### Generation of ATG16L1 K490A knockin mouse model

The K490A point mutation was introduced into C57/BL6 mice via direct zygote injection of CRISPR/Cas9 reagents. Briefly, a single stranded guide sequence was designed and synthesised along with a tracerRNA from Dharmacon. A repair donor single stranded DNA sequence was designed to introduce the K490A point mutation and mutate the PAM sequence to stop re-targeting of the Cas9 complex to already edited DNA. These reagents along with recombinant Cas9, were injected into mouse zygotes. Pups born from these injections were genotyped via Transnetyx and heterozogous founders were bred with wild type mice to obtain pure heterozygote animals. Further breeding yielded mice homozygous for the K490A mutation. Mice were housed in the Biological Support Unit at the Babraham Institute under specific pathogen–free conditions. All animal experiments at The Babraham Insitute were reviewed and approved by The Animal Welfare and Ethics Review Body and performed under Home Office Project license PPL/PO302B91A.

K490A guide sequence-GUUAGGGGCCAUCACGGCUCGUUUUAGAGCUAUGCUGUUUUG Repair donor ssDNAGCTGTCTCCCTTAGGTCAGAGAGAGTG*TGG*TCCGAGAGATGGAACTGTTAGGGGCGATCACCGCTTTGGACCTAA ACCCTGAGAGAACTGAGCTCCTGAGCTGCTCCCGTGATGACCTG

### Bone Marrow Derived Macrophage isolation

C57/BL6 wild-type and ATG16L1 K490A mice, aged 13–15 weeks, were used to obtain BMDCs. Bone marrow cells were isolated by flushing tibias and femurs with PBS + 2% FBS. Cells were pelleted and resuspended in 1 ml Red Blood Cell lysis buffer (150 mM NH4Cl, 10 mM KHCO3, 0.1 mM EDTA) for 2 min at room temperature. Cells were pelleted and resuspended in RPMI 1640 (Invitrogen 22409-031), 10% FBS, 1% Pen/Strep, 50 μM 2-mercaptoethanol supplemented with 20 ng/ml M-CSF (Peprotech #AF-315-02) and 50ng/ml Fungizone (Amphotericin B) (Gibco #15290018). Media was refreshed on days 3 and 6 and cells plated for assays on day 8.

### HEK293 GFP-LC3B ATG13/ATG16L1_DKO cells

ATG13_KO HEK293 cells stably expressing GFP-LC3B maintained in DMEM, 10% FBS, 1% pen/strep, were used as previously described^7^. To generate *ATG16L1* KO, gRNA sequence (GTGGATACTCATCCTGGTTC) with overhangs for containing a BpiI site was annealed and cloned into the pSpCas9(BB)-2A-GFP plasmid (Addgene, 48138; deposited by Dr. Feng Zhang) digested with the BpiI restriction enzyme (Thermo Scientific, ER1011). The recombinant plasmid along with a pBabe-puro construct (Addgene, 1764; deposited by Dr. Hartmut Land) expressing mouse ATG16L1 variants was transfected into HEK293 ATG13_KO GFP-LC3B cells via Lipofectamine 2000 (Invitrogen). Cells were selected with 2.5 μg/ml puromycin (P8833, Sigma) for 48 h, and single cell clones were obtained by limiting dilution. After clonal expansion, *ATG16L1 KO* clones were selected based on the absence of ATG16L1 protein as detected by Western blot.

RAW264.7 wild type and ATG16L1_KO cells were kindly provided by Dr Anne Simonsen^71^ and maintained in DMEM 10% FBS, 1% pen/strep.

### Live imaging time-lapse confocal microscopy

HEK293 cells were plated on 35 mm glass-bottomed dishes (Mattek, Ashland, MA). Images were acquired every 2 minutes using a spinning disk confocal microscope, comprising Nikon Ti-E stand, Nikon 60x 1.45 NA oil immersion lens, Yokogawa CSU-X scanhead, Andor iXon 897 EM-CCD camera and Andor laser combiner. All imaging with live cells was performed within incubation chambers at 37°C and 5% CO_2_. Image acquisition and analysis was performed with Andor iQ3 (Andor Technology, UK) and ImageJ.

### Endogenous Calcium Imaging

Hela wild-type and Hela ATG16L1KO cells were trypsinized and seeded at 20000 per well of PDL coated Greiner Bio plates for 2 h. Cells were loaded with 10 µl of Calcium 6 dye solution for 1.5 h at room temperature. After incubation, the dye was removed from the plates and replaced with 10µl of low Ca^2+^ solution containing 145 mM NaCl, 5 mM KCl, 3 mM MgCl_2_, 10 mM glucose, 1 mM EGTA and 20 mM HEPES at pH 7.4. With 1 mM EGTA, the free Ca^2+^ concentration is estimated to be < 10 nM based on the Maxchelator software (http://maxchelator.stanford.edu/). Compounds plates were prepared with low calcium solution. Cell and compound plates were loaded onto the FLIPR and the 15 min protocol was run. The fluorescence intensity at 470nm was monitored. After an initial 10 second baseline read, compounds were added to the cells. Images were taken for 15 minutes to monitor effects on Ca^2+^ fluorescence. Data exported as max-min Relative Fluorescence Unit (RFU).

### Recombinant protein expression

#### Protein purification

For evaluation of the FLCN/FNIP2/GABARAP complex by size exclusion chromatography, FLCN and FNIP2 were subcloned as twin-strep-FLAG and GST fusion proteins, respectively, and purified as described^20^. Final purified complexes were snap frozen in liquid nitrogen in buffer A (25 mM HEPES pH 7.4, 130 mM NaCl, 2.5 mM MgCl_2_, 2 mM EGTA, and 0.5 mM TCEP). Full-length human GABARAP (1-117) was subcloned with a C-terminal MBP tag (GABARAP_MBP) separated by a GSSGSS linker in pET21b and expressed in *E. coli* following induction at 16°C for 16 hours in LB. Cells were lysed in 50 mM Tris pH 7.4, 500 mM NaCl, 0.5 mM TCEP, 0.1% Triton X-100, 1 mM PMSF, and 15 ug/ml benzamidine; sonicated; and clarified by centrifugation. GABARAP_MBP was purified using amylose resin equilibrated in wash buffer (50 mM Tris pH 7.4, 500 mM NaCl, 0.5 mM TCEP) and eluted with wash buffer plus 30 mM maltose. The protein was further purified by size exclusion chromatography using a Superdex 75 column equilibrated buffer A and snap frozen in liquid nitrogen. Purified GABARAP_MBP was mixed with FLCN/FNIP2 at a ratio of 1:0.8 and gently mixed for 2 hours at 4 °C. The sample was injected onto a Superose 6 Increase (GE) column (1CV = 24 mL) that was pre-equilibrated in Buffer A. The retention time of peak fractions were compared to FLCN/FNIP2 and GABARAP_MBP alone followed by evaluation of samples using 12% SDS-PAGE.

For GEE labelling studies, full-length human FLCN-PreScission-MBP and His8-TEV-FNIP2 were subcloned for co-expression in mammalian cells. Cells were lysed in 50 mM Tris pH 8.0, 200 mM NaCl, 10% glycerol, 1% Triton X-100, and 2 mM MgCl_2_ with Protease Inhibitor Cocktail (Roche) and clarified by centrifugation. Protease Inhibitor tablets were included throughout purification to prevent degradation. The FLCN/FNIP2 complex was purified over an amylose column equilibrated in wash buffer (50 mM Tris pH 8.0, 500 mM NaCl, 10% glycerol, 1% Triton X-100, and 2 mM MgCl_2_), washed and eluted in wash buffer plus 10 mM maltose. The complex was further purified by IMAC chromatography and eluted in buffer B (50 mM Tris pH 8.0, 500 mM NaCl, 10% glycerol, 0.11 mM NG311, and 2 mM MgCl_2_) plus 250 mM imidazole followed by overnight cleavage with PreScission Protease^TM^ at 4°C. The cleaved MBP tag was removed with a second amylose column, and the complex was concentrated prior to purification by size exclusion chromatography using a Superdex 200 column equilibrated in buffer B. The sample was snap frozen in liquid nitrogen. GABARAP (1-117) was cloned as GST-PreScission-His8-TEV-GABARAP for expression in *E. coli* and lysis as described for GABARAP_MBP. GABARAP was purified using glutathione resin equilibrated in buffer C (50 mM Tris pH 8.0, 500 mM NaCl, and 0.5 mM TCEP) and eluted using buffer C plus 10 mM glutathione. The GST tag was cleaved overnight at 4°C with PreScission Protease and purified by size exclusion chromatography using a Superdex 75 column equilibrated in buffer D (50 mM Tris pH 8.0, 150 mM NaCl, and 0.5 mM TCEP). His8-TEV-GABARAP was applied to an IMAC column equilibrated in buffer E (25 mM Tris pH 8.0, 150 mM NaCl, and 0.5 mM TCEP), eluted in buffer E plus 250 imidazole, and dialyzed into buffer E. Final samples were concentrated and snap frozen in liquid nitrogen.

For X-ray crystallography, a single polypeptide clone was designed using GST-PreScission-His8-TEV followed by residues 558-576 of FNIP2, a Gly-Ser spacer, and full-length GABARAP. Protein was expressed and purified using glutathione resin as described for GST-PreScission-His8-TEV-GABARAP. The tag was cleaved using TEV protease and purified by size exclusion chromatography using a Superdex 75 column in buffer D. Residual tag was removed by IMAC chromatography and the final protein was exchanged into buffer E by dialysis, concentrated to 18.7 mg/mL prior, and snap frozen in liquid nitrogen.

For evaluation of the binding affinity of GABARAP and LC3B for the FLCN/FNIP2 complex via SPR, full-length human FLCN-8xG-AviTag-PreScission-MBP and full-length human His8-TEV-FNIP2 were expressed in mammalian cells and purified above as described for GEE labelling studies. Biotinylation of the AviTag in full-length human FLCN-8xG-AviTag-PreScission-MBP was performed following according to the manufacturer’s (Avidity) suggested protocol and was performed following PreScission protease cleavage and prior to size exclusion chromatography. Full-length human GABARAP (1-117) was purified as described above for GEE studies. Full-length human LC3B (1-125) was cloned as GST-PreScission-His8-TEV-LC3B for expression in *E. coli* and lysis as described for GABARAP_MBP. LC3B was purified using glutathione resin equilibrated in buffer C (50 mM Tris pH 8.0, 500 mM NaCl, and 0.5 mM TCEP) and eluted using buffer C plus 10 mM glutathione. The GST tag was cleaved overnight at 4°C with PreScission Protease and purified by size exclusion chromatography using a Superdex 75 column equilibrated in buffer D (50 mM Tris pH 8.0, 150 mM NaCl, and 0.5 mM TCEP). His8-TEV-LC3B was applied to an IMAC column equilibrated in buffer E (25 mM Tris pH 8.0, 150 mM NaCl, and 0.5 mM TCEP), eluted in buffer E plus 250 imidazole, and dialyzed into buffer E. Final samples were concentrated and snap frozen in liquid nitrogen.

For evaluation of the binding affinity of GABARAP and LC3B for the p62 FIR domain via SPR, the FIR region of human p62 (326-380) with the 4P mutations (S349E, S365E, S366E, S370E)^72^ was cloned as GST-PreScission-p62 FIR 4P for expression in *E. coli* following induction at 18°C for 16 hours in TB. Cells were lysed in 50 mM HEPES pH 7.0, 300 mM NaCl, 5 mM MgCl_2_, 2 mM CaCl_2_, 1 mM TCEP and 1 U/mL DNase I; sonicated; and clarified by centrifugation. GST-PreScission-p62 FIR 4P was purified using glutathione resin equilibrated in 50 mM HEPES pH 7.0, 300 mM NaCl, 1 mM TCEP and eluted using the same buffer supplemented with 10 mM glutathione. The GST tag was left on the protein, and GST-PreScission-p62 FIR 4P was further purified by size exclusion chromatography using a Superdex 200 column equilibrated in 50 mM HEPES pH 7.0, 300 mM NaCl, 1 mM TCEP. Fractions containing GST-PreScission-p62 FIR 4P were pooled, concentrated and snap frozen in liquid nitrogen. GST protein used as a reference for SPR was purchased from Cytiva.

### Protein Crystallization

FNIP2-GABARAP chimera was crystallized by the vapor diffusion method using equal volumes protein and 0.1 M Magnesium acetate, 0.1 M MOPS pH 7.5, 12% v/v PEG 8000 in sitting drops. Crystals were frozen using well solution supplemented with 20% glycerol. Data were collected at beamline BL18U1 of the Shanghai Synchrotron Radiation Facility (SSRF) and processed with XDS^73^ and AIMLESS^74^. The structure was solved by molecular replacement with PHASER^75^ using 6hyo^40^ as a search model and refined with REFMAC5^76^ and PHENIX^77^. Model building was performed using COOT^78^.

### Peptide Synthesis

Wild Type and mutant peptides corresponding to residues 550-576 (Long) or 558-576 (Short) of FNIP2 were ordered from New England Peptide. All peptides were purified to >99% purity, with the exception of the WT Long FNIP2 peptide that was purified to 70% purity. 5 mg of each short peptide was resuspended in water to a concentration of 500 µM, whereas 5 mg aliquots of long peptides were resuspended in water supplemented with 3 uL of 1M ammonium bicarbonate to a final concentration of 500 µM.

### Measurement of *in vitro* binding kinetics of GABARAP and LC3B with FLCN/FNIP2 by Surface Plasmon Resonance (SPR)

Direct binding of GABARAP and LC3B to immobilized FLCN/FNIP2 was assayed on a Biacore S200 (Cytiva) using the Biotin CAPture Kit (Series S Sensor Chip CAP and Biotin CAPture reagent) (Cytiva). Prior to beginning each run, the S200 system was equilibrated using a 4x Multiprime of running buffer (1x HBS-P+ (Cytiva) supplemented with 1 mM TCEP). The chip surface was conditioned with three successive 60 second pulses of 6M Guanidine HCl / 50 mM NaOH at a flow rate of 30 µL/min. Prior to kinetic analysis, three startup cycles were performed using standard conditions for the CAPture Kit (Cytiva). For each cycle of kinetic analysis, undiluted Biotin CAPture reagent was injected onto all four flow cells for 150 seconds at a flow rate of 4 µL /min. 40 µg/mL Biotinylated FLCN-8xGlycine-AviTag/FNIP2 was injected onto flow cells 2 and 4 at 10 µL/min for 60 seconds to produce an immobilization level of 100-150 Response Units (RU). 0.06 – 31.25 nM GABARAP or 0.06 to 4000 nM LC3B protein was injected using “High performance Multi-cycle Kinetics” or “Single Cycle Kinetics” injections for 60 seconds at a flow rate of 100 µL/min, and were allowed to dissociate from FLCN/FNIP2 for 750 seconds prior to regeneration of the chip surface with 30 second pulses of 6M Guanidine HCl / 50 mM NaOH followed by 30% Acetonitrile / 250 mM NaOH.

Sensorgrams were processed with Cytiva Biacore S200 Evaluation Software (version 1.1, Build 28). Double referencing was performed by first subtracting the GABARAP/LC3B responses over the reference flow cells (1 and 3) from the corresponding responses in sample flow cells (2 and 4, respectively), followed by subtraction of averaged sensorgrams for 0 nM GABARAP/LC3B from sensorgrams corresponding to all other concentrations of GABARAP/LC3B. After double referencing kinetic data, injection and pump spikes were manually removed from sensorgrams and the data were fit globally by non-linear regression to a simple 1:1 Langmuir binding model without a Bulk RI term (RI = 0 RU) to determine association/dissociation rate constants (k_a_, K_d_), analyte binding capacity (R_max_) and the equilibrium dissociation constant (K_D_). R_max_ values obtained during kinetic analysis represented 50-80% of the theoretical R_max_ value indicating a surface activity of 50-80%. Sensorgrams and 1:1 binding model curve fits were exported from the S200 Evaluation Software and replotted in GraphPad Prism (v8.4.3).

### Measurement of *in vitro* binding kinetics of GABARAP and LC3B with the p62 FIR 4P Domain by Surface Plasmon Resonance (SPR)

Direct binding of GABARAP and LC3B to immobilized GST-p62 FIR 4P protein was assayed on a Biacore S200 (Cytiva) using a CM5 chip (Cytiva). Prior to beginning each run, the S200 system was equilibrated using a 4x Multiprime of 1x HBS-N buffer (Cytiva). The CM5 chip surface was preconditioned using five cycles of successive 60 second injections of 50 mM NaOH, 100 mM HCl, 0.2% SDS and nuclease free water at 30 µL/min. GST-p62 FIR 4P (Sample cells, 2 and 4) or GST (Reference cells, 1 and 3) were covalently coupled to the chip surface using NHS/EDC supplied in the Cytiva Amine Coupling Kit. GST-p62 and GST (Cytiva) were diluted to 0.1 µM in 10 mM Sodium Acetate, pH 4.5 and injected onto flow cells 2 and 4 (GST-p62) or 1 and 3 (GST) of the NHS/EDC activated chip until 250 RU of GST-p62 or 200 RU of GST was immobilized. Ethanolamine was then injected across all four flow cells according to the Amine Coupling Kit protocol to block remaining reactive sites on the chip surface.

The SPR system was then equilibrated using a 4x Multiprime of running buffer (1x HBS-P+ (Cytiva) supplemented with 1 mM TCEP). Prior to kinetic analysis, ten startup cycles were performed using 60 second sample injections of running buffer at 30 µL/min and 30 second dissociation times followed by two 30 second injections of pH 2.0 Glycine (Cytiva) at 30 µL/min and a “Carry-over control” injection. For each cycle of kinetic analysis, 40 – 10,000 nM GABARAP or LC3B protein was injected across all four flow cells using “High performance Multi-cycle Kinetics” injections for 60 seconds at a flow rate of 100 µL/min, and were allowed to dissociate from GST-p62 FIR 4P and GST surfaces for 1000 seconds prior to regeneration of the chip surface with two 30 second injections of pH 2.0 Glycine (Cytiva) at 30 µL/min followed by a 300 second stabilization period and a “Carry-over control” injection.

Sensorgrams were processed with Cytiva Biacore S200 Evaluation Software as described above. While sensorgrams for LC3B binding to GST-p62 FIR 4P domain showed high affinity binding with a very slow off rate (Estimated 80 nM K_D_ with 1200 second half life), they were more complicated than could be accurately described using a simple 1:1 Langmuir binding model with a Bulk RI term. In spite of their complexity the LC3B sensorgrams and sensorgrams for GABARAP binding to GST-p62 reached a steady state during injection onto the chip surface and were fit using a “Steady State Affinity Model” with the offset parameter set to zero. R_max_ values obtained during steady state affinity analysis represented ∼85% of the theoretical R_max_ value indicating a surface activity of ∼85%. Sensorgrams and steady state affinity binding isotherm fits were exported from the S200 Evaluation Software and replotted in GraphPad Prism (v8.4.3).

### Measurement of the Affinity of p62, FLCN/FNIP1 and FNIP2 Peptides for GABARAP by Competition in Solution / Affinity in Solution Surface Plasmon Resonance (SPR)

We used the ‘Competition in solution’ (also called ‘affinity in solution’) method^79–81^ to measure the affinity of the GST-p62 FIR 4P domain, full length FLCN/FNIP1 protein and FNIP2 Peptide (New England Peptide) competitors for GABARAP. Full length AviTag-FLCN/FNIP2 heterodimers were immobilized on the Series S Sensor Chip CAP surface (as described above for FLCN/FNIP binding kinetics with GABARAP and LC3B) and used to capture and measure the concentration of free GABRAP in pre-equilibrated mixtures containing a constant amount of GABARAP with varying amounts of soluble p62, FLCN/FNIP1 and FNIP2 peptide competitors.

Calibration curves of 0 - 15.6 nM GABRAP were prepared by two-fold serial dilution of 31.2 nM GABARAP into running buffer (1x HBS-P+ buffer (Cytiva) supplemented with 1 mM TCEP). Calibration curves of GABARAP were injected onto the chip surface for 60 seconds at 30 µL/min and the maximum RU for each concentration of GABARAP was recorded. Following sample injections, the chip surface was regenerated by injecting one 30 second pulse of 6M Guanidine Hydrochloride with 0.25 M NaOH and a second 30 second pulse of 30% acetonitrile with 0.25 M NaOH, followed by a ’carry over control injection’.

Mixtures containing a fixed amount of GABARAP (7.8 nM) and soluble GST-p62 FIR, FLCN/FNIP1 protein and FNIP2 peptide competitors were prepared by 1:1 dilution of 15.6 nM GABARAP with two-fold serial dilutions of competitors in running buffer. Each mixture of GABARAP and competitor was injected to produce a series of sensorgrams that were recorded as described for the GABARAP calibration curve.

Sensorgrams were processed with Cytiva Biacore S200 Evaluation Software (version 1.1, Build 28). Double referencing was performed by first subtracting the GABARAP/LC3B responses over the reference flow cells (1 and 3) from the corresponding responses in sample flow cells (2 and 4, respectively), followed by subtraction of averaged sensorgrams for 0 nM GABARAP/LC3B from sensorgrams corresponding to all other concentrations of GABARAP/LC3B.

Sensorgrams were processed with the Cytiva Biacore S200 Evaluation Software. The y-axes were zeroed at the baseline for each cycle and x-axes were aligned at the injection start. Bulk refractive index changes and systematic deviations in sensorgrams were removed by double referencing as described above. The concentration of free GABARAP in each mixture with competitors was determined from the calibration curve, exported into Prism, plotted against the log of competitor concentration and fit to the equation detailed on page 192 of the Biacore S200 Software Handbook (Revision 29-1431-08 AA). The equation assumes the existence of a single binding site between FLCN/FNIP2 competitors and GABARAP.

### Bacterial strains and infections

Infections were performed with wild-type *S*. Typhimurium SL1344 and isogenic mutant lacking the SopF effector. Mutation in the *S*. Typhimurium SL1344 background lacking SopF (Δ*sopF*) was a kind gift from Dr. Feng Shao and described previously^25^. A previously established approach was used for infection of epithelial cells, using late-log *S*. Typhimurium cultures as inocula^82^. Briefly, subcultured *Salmonella* strains were pelleted at 10,000g for 2 min, resuspended and diluted 1:50 in PBS, pH 7.2, and added to cells for 10 min at 37 °C. Selection for intracellular bacteria was performed at 30 min p.i. using 100 μg ml^−1^ gentamicin, a concentration that was decreased to 10 μg ml−1 at 2 h p.i. for maintenance purposes. If applicable, cells were fixed with 4% paraformaldehyde in PBS at 37 °C for 10 min.

## Acknowledgements

We would like the thank Drs. James Hurley, Sascha Martens, Keith Dionne, Frank Gentile, Bob Tepper, and the employees of Casma Therapeutics for critical feedback in preparation of this manuscript. We thank all staff in the Biological Support Unit at the Babraham Institute for their outstanding care of our mouse colony and Dr Simon Walker in the Imaging Facility for his support. We thank Dr Domink Spensberger for his role in generating the K490A mouse and Dr Elise Jacquin for early work on vATPase. We thank Dr Anne Simonsen for providing the ATG16L1 KO RAW264.7 cell line. This work was supported by grants from the BBSRC, BB/P013384/1 (BBS/E/B/000C0432 and BBS/E/B/000C0434), BB/R019258/1, Cancer Research UK Career Development award C47718/A16337 and the Canadian Institutes of Health Research (FDN#154329 to J.H.B). J.H.B. holds the Pitblado Chair in Cell Biology.

## Author Contributions

J.M.G, L.O.M, and O.F designed all experiments and wrote the manuscript. J.M.G., W.G.W. IV, K.H., T.L., and S.K performed experiments. K.F. and O.F. generated the ATG16L1 K490A knockin mouse model. A.N. performed data analysis of RNAseq profiling, A.J, A.A., S.R.B., and S.R.R. performed calcium imaging experiments. C.K-I. performed ultrastructural analysis. J.G.F. provided chemistry support. M.W., B.A., and D.B. performed protein purification and complex formation assays. A.B. provided valuable scientific guidance throughout the study.

## SUPPLEMENTARY MATERIAL

**Supplementary movie S1. Time lapse imaging of GFP-LC3B puncta formation in HEK293T WT cells upon AZD8055 treatment. Images taken at 2 minute intervals. AZD8055 = 1uM.**

**Supplementary movie S2. Time lapse imaging of GFP-LC3B puncta formation in HEK293T WT cells upon C8 treatment.** Images taken at 2 minute intervals. C8 = 2uM.

**Supplementary movie S3. Reconstruction of FIB-SEM imaging in HEK293T WT cells upon AZD8055 treatment.** Ultrastructural analysis of colocalization events HEK293T WT cells stably expressing GFP-LC3B and LAMP1-RFP were imaged by correlative light electron microscopy. Image compilation for zoomed ROI.

**Supplementary movie S4. Reconstruction of FIB-SEM imaging in HEK293T ATG13_KO cells upon C8 treatment.** Ultrastructural analysis of colocalization events HEK293T ATG13_KO cells stably expressing GFP-LC3B and LAMP1-RFP were imaged by correlative light electron microscopy. Image compilation for zoomed ROI.

---

**Supplementary figure 1.**
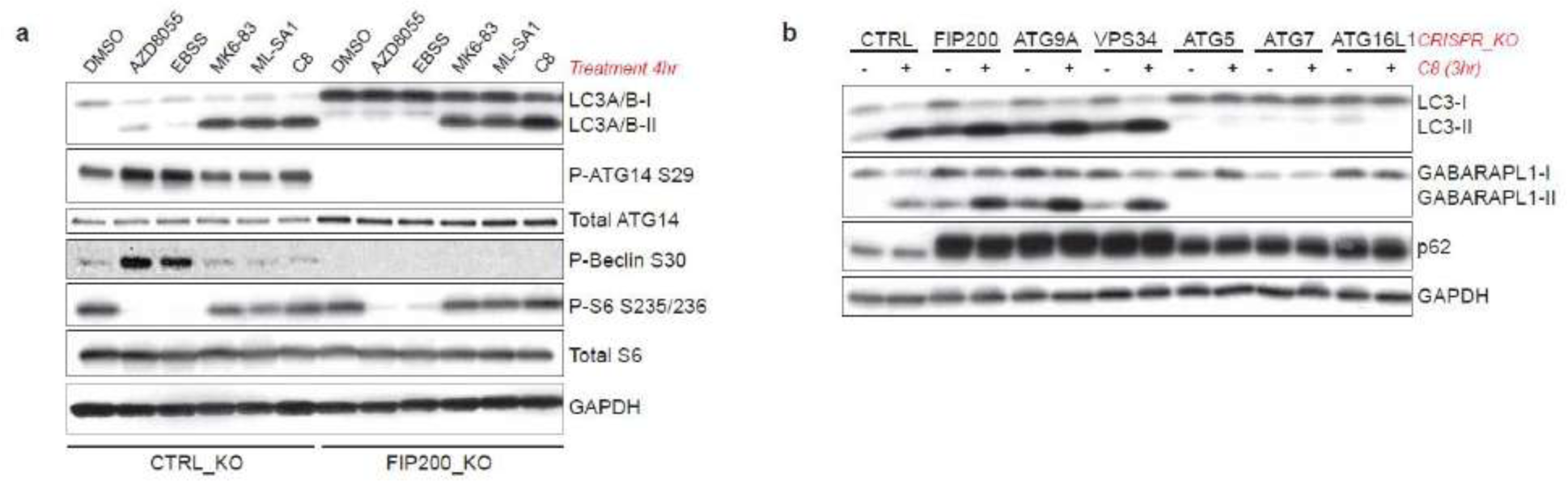
TRPML1 agonists stimulate ATG8 lipidation in autophagy-deficient cells. **(a)** U2OS cells stably expressing Cas9 were deleted for the ULK1 complex component FIP200. Cells were then treated with the indicated compounds for 4 hr. P-ATG14 S29 and P-Beclin S30 are specific ULK1-dependent phosphorylation sites. **(b)** HeLa cells stably expressing Cas9 were deleted for the indicated autophagy components and then treated with compound C8 (2uM) for 3hr. Lipidation of the ATG8 homologs LC3A/B and GABARAPL1 could be observed irrespective of autophagosome biogenesis.

**Supplementary figure 2.**
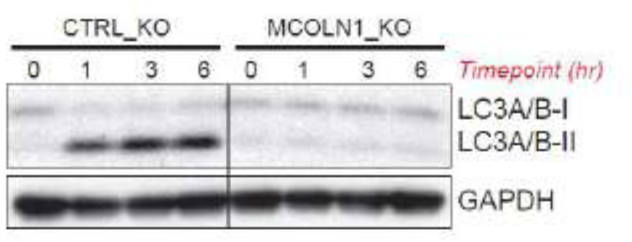
TRPML1 agonists induce rapid lipidation of LC3 through an on-target mechanism. CRISPR knockout of TRPML1 in HeLa cells stably expressing Cas9 abolishes LC3-II induction upon TRPML1 agonist (C8) treatment.

**Supplementary figure 3.**
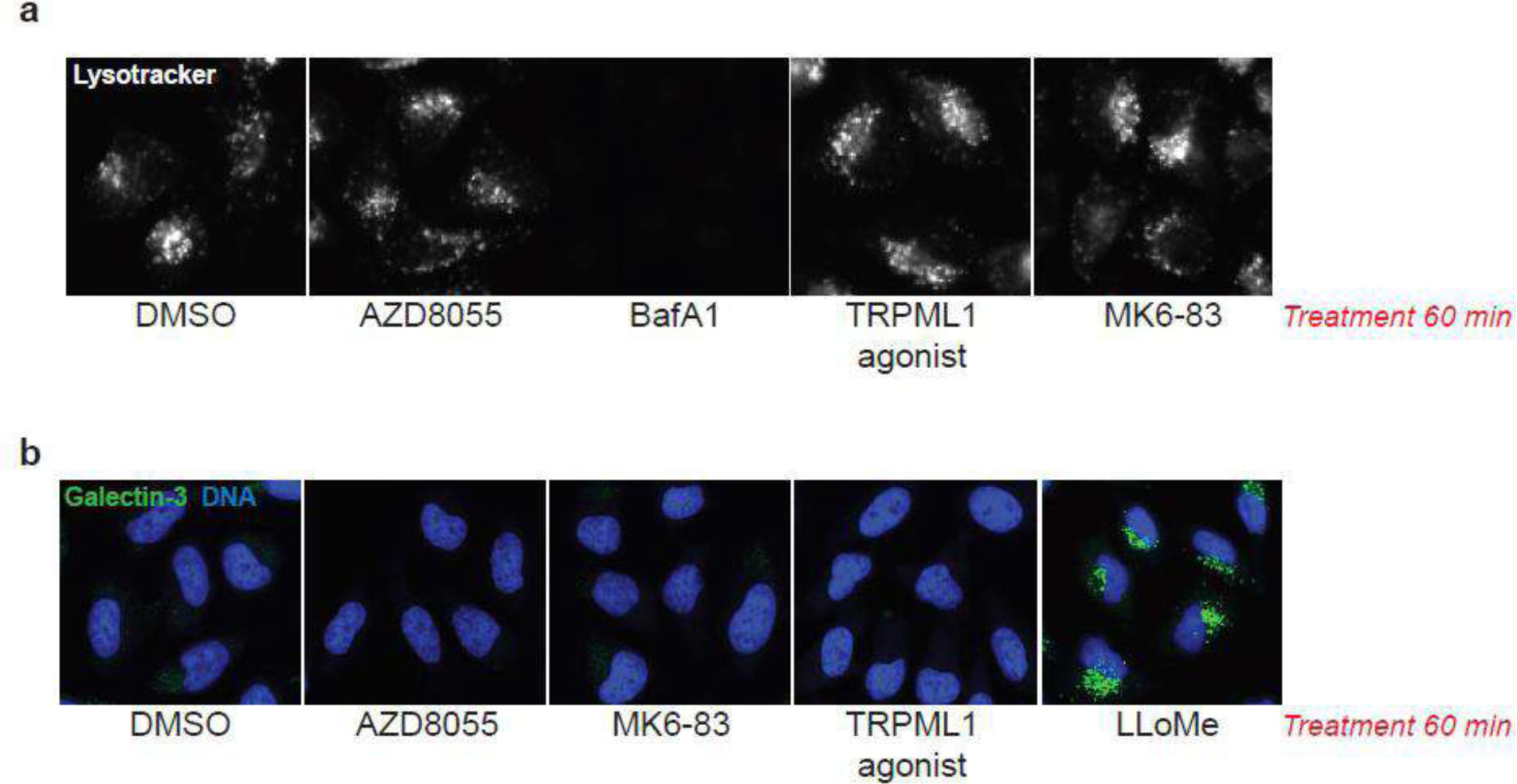
TRPML1 activation does not result in loss of the lysosomal pH gradient or membrane damage. **(a)** HeLa cells were treated for 60 min with the indicated compounds and stained with the pH-sensitive Lysotracker dye. BafA1 serves as a positive control. Representative images are shown. **(b)** HeLa cells were treated for 60 min with the indicated compounds, fixed, and stained for immunofluorescence analysis of Galectin-3 puncta. LLoMe (1mM) served as a positive control. Representative images are shown.

**Supplementary figure 4.**
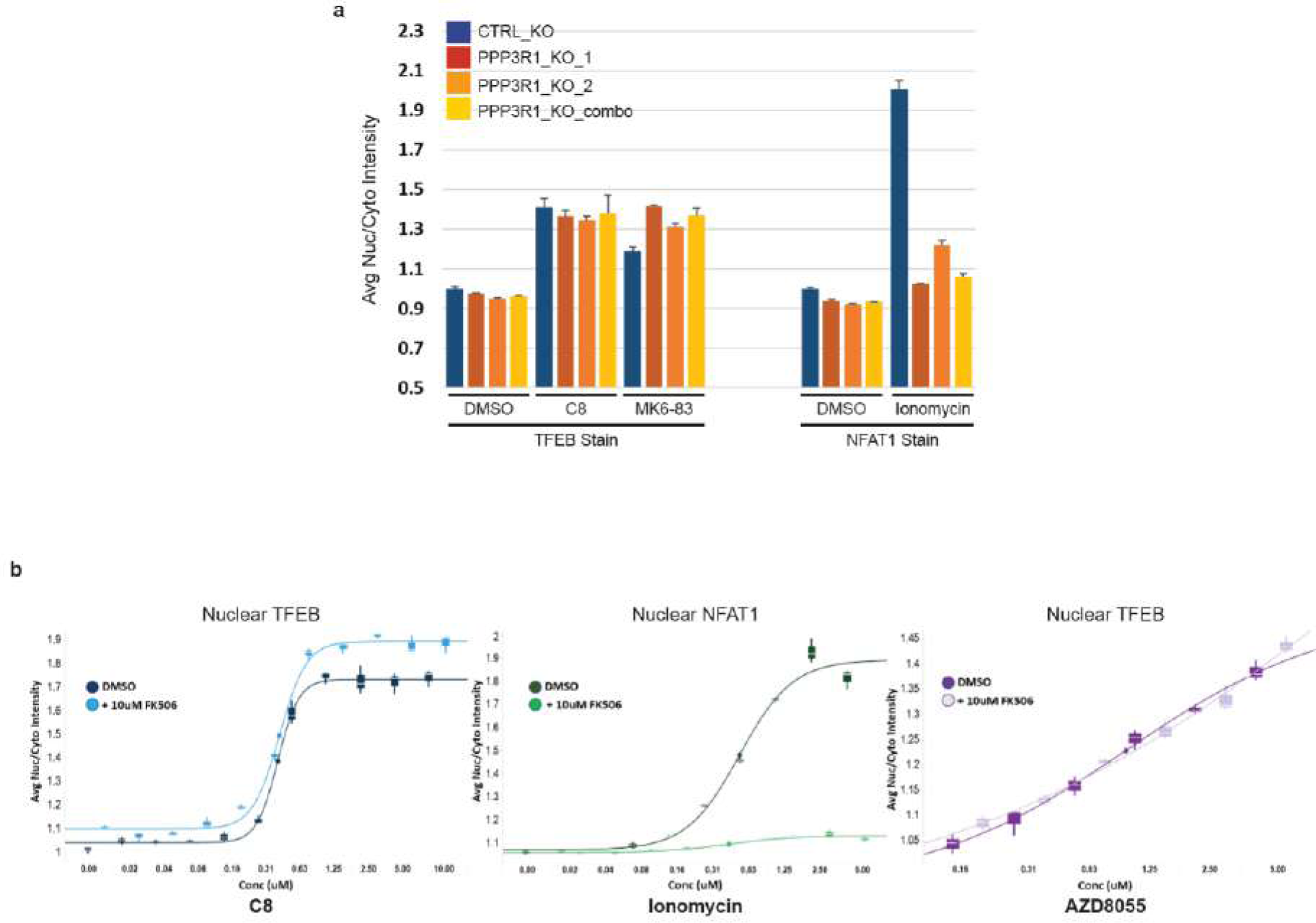
CRISPR knockout of the essential calcineurin (CaN) regulatory subunit PPP3R1 does not impact TFEB nuclear localization upon TRPML1 activation, yet does block activation of the canonical CaN substrate NFAT1 by ionomycin. **(a)** HeLa cells stably expressing Cas9 were knocked out for PPP3R1 using CRISPR. PPP3R1_KO_1 and PPP3R1_KO_2 represent pooled population derived from distinct gRNA sequences. PPP3R1_KO_combo represents cell infected with both gRNA sequences. Cells were treated with the indicated compounds for 3 hours, fixed, and stained for endogenous TFEB or NFAT1. Analysis was performed using high content imaging. Mean ± SD. Minimum of 1500 cells quantified per condition. **(b)** Dose response analysis of TFEB nuclear localization upon cotreatment with the CaN inhibitor FK-506. FK-506 did not impact nuclear TFEB upon TRPML1 agonist (C8) or AZD8055 treatment despite effective inhibition of CaN-mediated NFAT translocation upon ionomycin treatment. Cells were treated as in (A). Mean ± SD. Minimum of 1500 cells quantified per condition.

**Supplementary figure 5.**
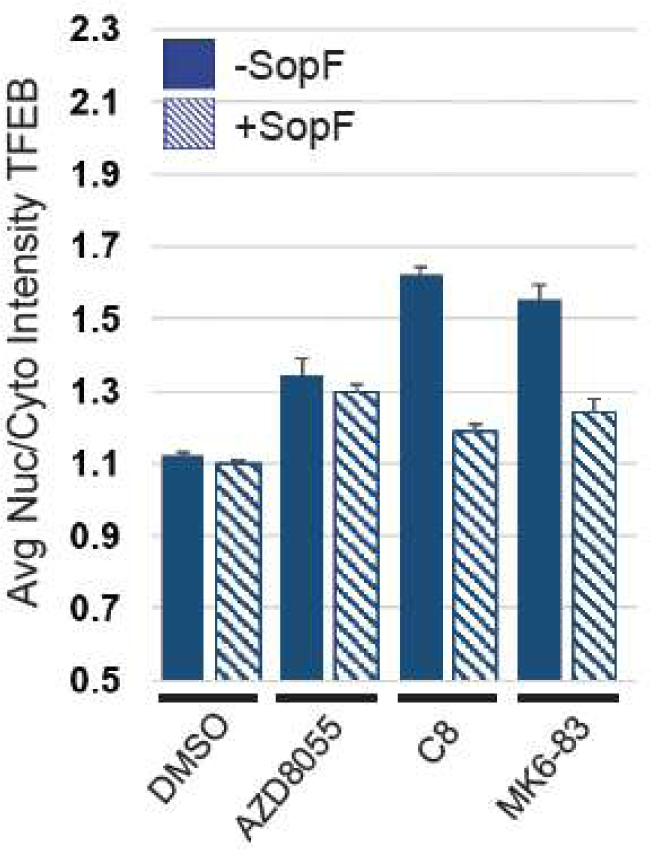
HeLa cells expressing inducible-SopF were treated with the indicated compounds for 3 hr. DOX induction of SopF was begun 24 hr prior to compound treatment. Mean ± SD. Minimum of 1500 cells quantified per condition.

**Supplementary figure 6.**
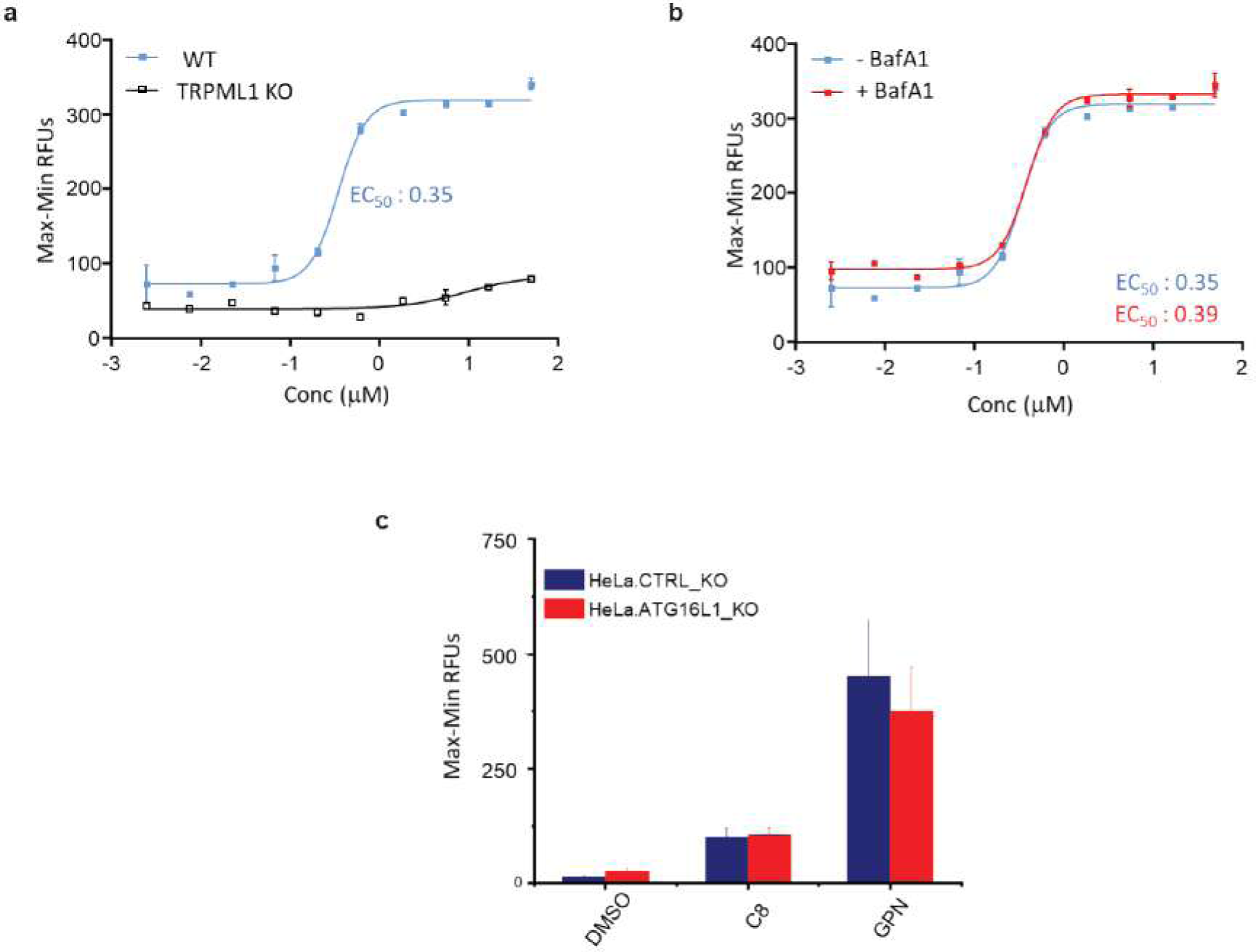
BafA1 treatment or ATG16L1 knockout does not affect TRPML1-dependent calcium release. **(a)** Whole-cell calcium imaging in wild-type or TRPML1_KO HeLa cells upon dose-response treatment with C8. Mean ± SD plotted from three wells per datapoint. **(b)** BafA1 cotreatment (100nM) in wild-type HeLa cells upon dose response treatment with C8. Mean ± SD plotted from three wells per datapoint. **(c)** ATG16L1 knockout does not impact C8 induced calcium release. GPN treatment shown to examine total lysosomal calcium content. Mean ± SD plotted from three wells per datapoint.

**Supplementary figure 7.**
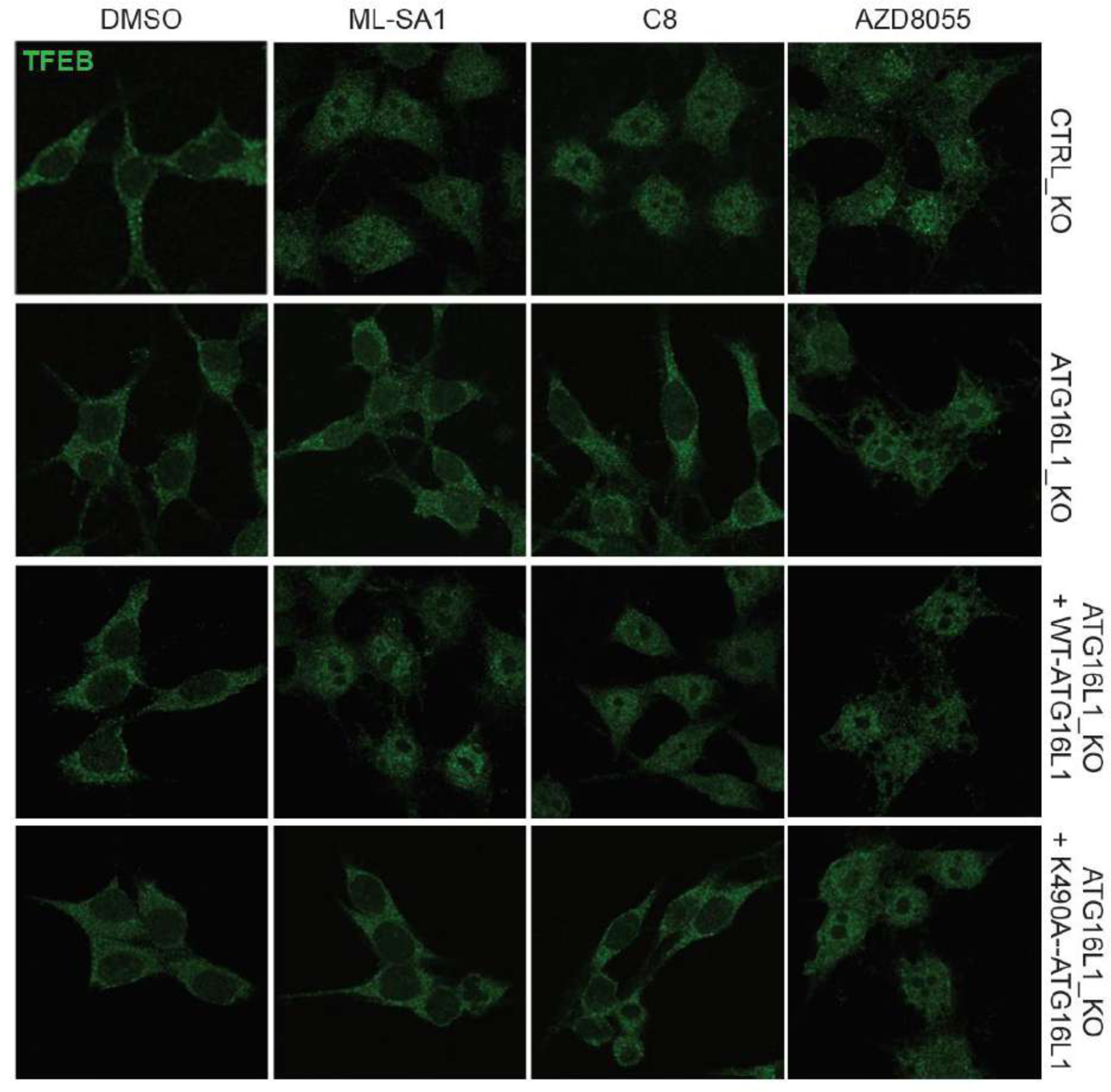
Single membrane ATG8 conjugation is required for TRPML1 agonist induced nuclear localization of TFEB. RAW264.7 murine macrophages were genetically engineered to knockout endogenous ATG16L1 and reconstituted with either WT-ATG16L1 or K490A-ATG16L1. Cells were treated with the indicated compounds for 2 hours, fixed and stained for endogenous TFEB. Representative images are shown. AZD8055 effectively translocates TFEB in all lines, however the TRPML1 agonists only retain activity with competent ATG8 conjugation to single membranes.

**Supplementary figure 8.**
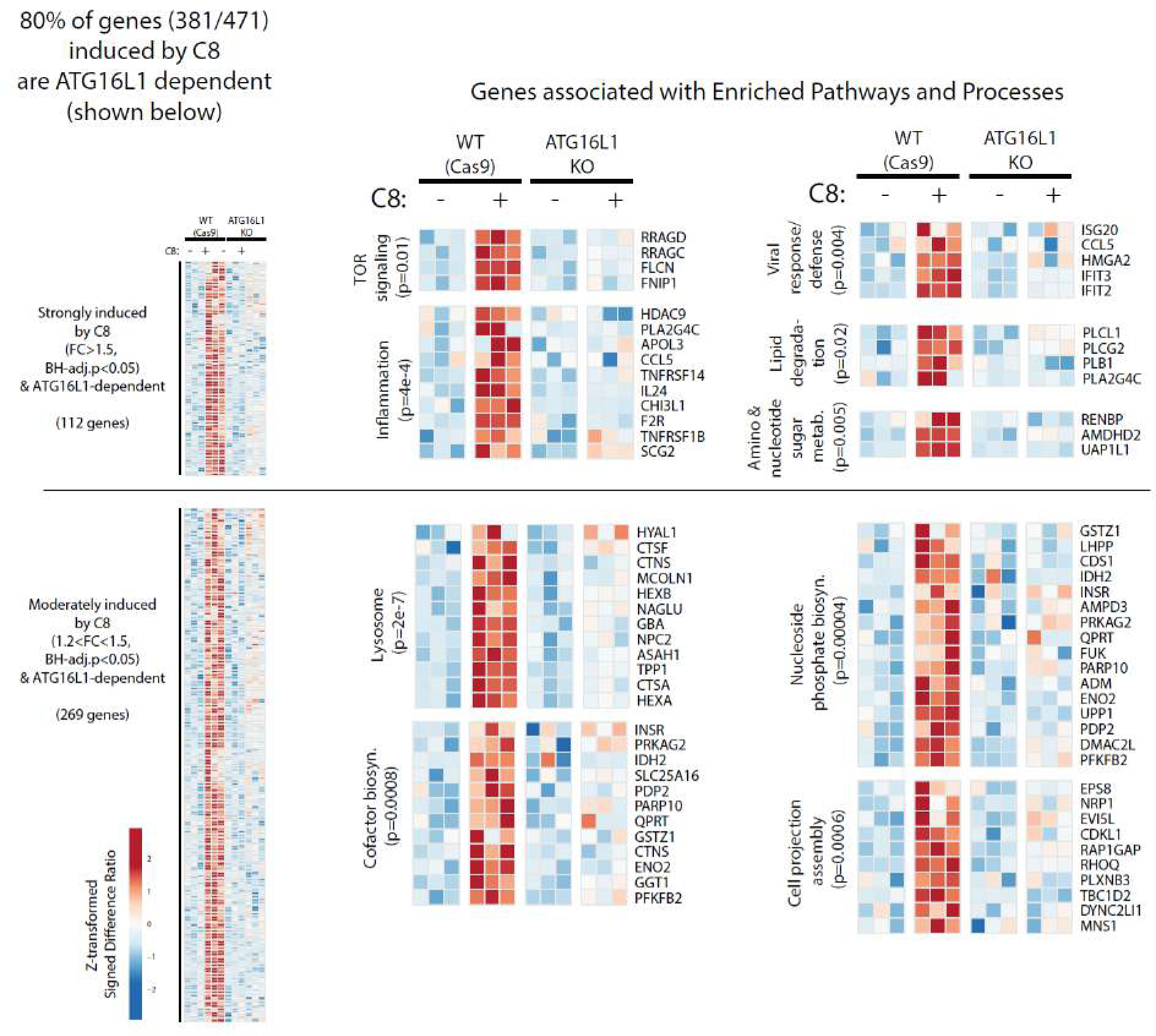
Representation of ATG16L1-dependent differentially expressed genes upon treatment with C8. RNAseq transcriptomics profiling of genes induced by C8 compound stimulation for 24h in ATG16L1 KO and WT HeLa cells (expressing Cas9). The heatmap shows Log_2_(CPM)-derived values for these genes, expressed as Z-transformed signed difference ratios (SDR) relative to their respective unstimulated baseline controls (either ATG16L1 KO or WT) and then scaled by normalizing to the maximum absolute deviation of each gene’s expression level from the unstimulated control. Differentially induced genes were identified by having a fold change (FC) >1.5 and Benjamini and Hochberg’s (BH)-adjusted p<0.05. Genes were partitioned based of fold change. Strongly induced genes >1.5 FC. Moderately induced genes 1.2-1.5 FC. Differentially induced genes were identified by having a Benjamini and Hochberg’s (BH)-adjusted p<0.05. Pathway enrichment analysis shows lysosomal genes are significantly increased in the moderate fraction upon C8 treatment (2uM for 24hr).

**Supplementary figure 9.**
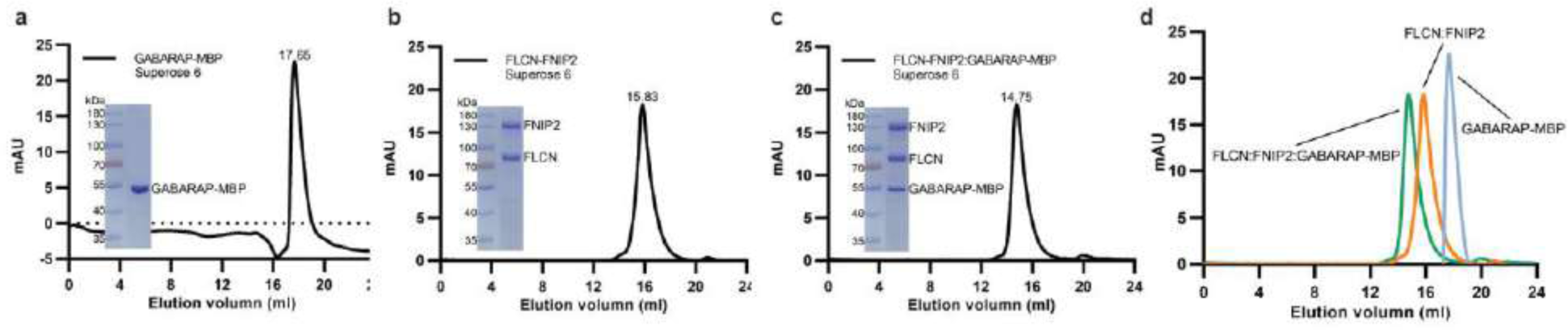
GABARAP_MBP complexes with FLCN/FNIP2 over a Superose 6 column. **(a)** Size exclusion chromatography elution profile of GABARAP_MBP. **(b)** Size exclusion chromatography of Strep/Flag tagged FLCN/FNIP2. **(c)** Complexing and size exclusion profile of GABARAP_MBP and Strep/Flag_FLCN/FNIP2. **(d)** Overlay of individual and complexed chromatograms.

**Supplementary figure 10.**
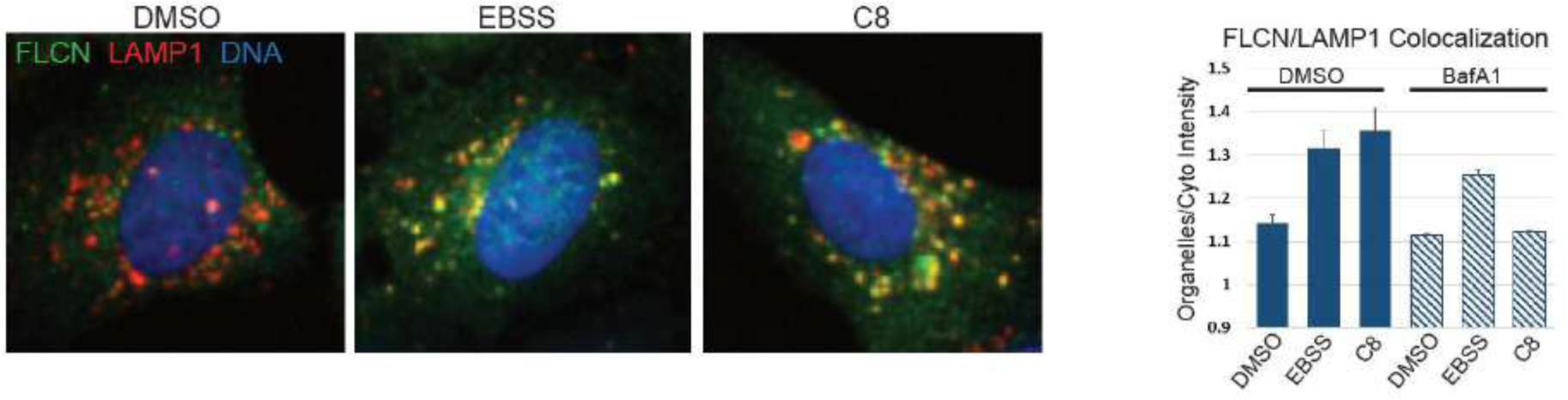
FLCN recruitment to lysosomes upon TRPML1 activation is sensitive to vATPase inhibition, unlike recruitment stimulated by nutrient starvation. **(a)** Representative images of FLCN/LAMP1 colocalization upon EBSS or TRPML1 agonist treatment. **(b)** Quantification of FLCN/LAMP1 colocalization. Organelle masks were established on the endogenous LAMP1 staining and the intensity ratio of endogenous FLCN in the organelle/cytoplasm was measured using high content imaging. Mean ± SD. Minimum of 1500 cells quantified per condition.

**Supplementary figure 11.**
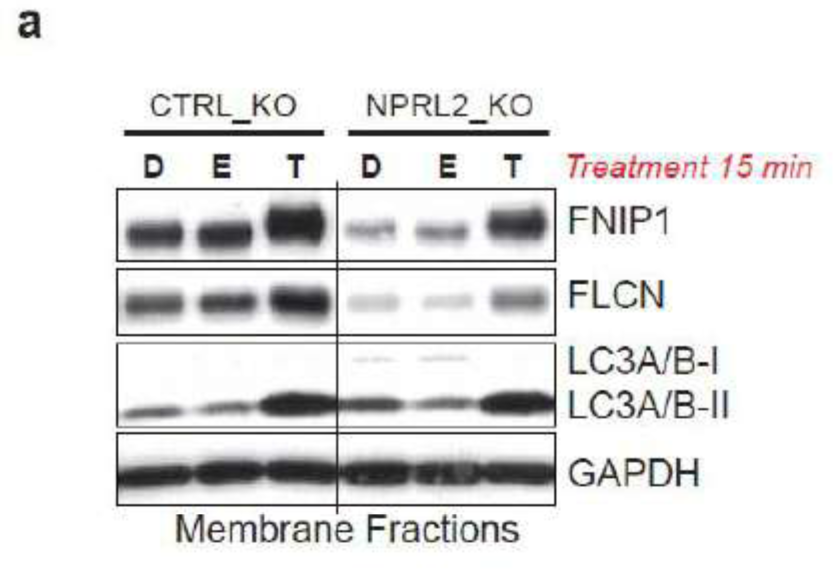
TRPML1 agonist regulates FLCN-FNIP membrane sequestration in LFC-deficient NPRL2_KO cells (const. RagA^GTP^). **(a)** Western blot analysis of FLCN-FNIP1 membrane recruitment upon TRPML1 agonist treatment or EBSS starvation in NPRL2_KO cells.

**Supplementary figure 12.**
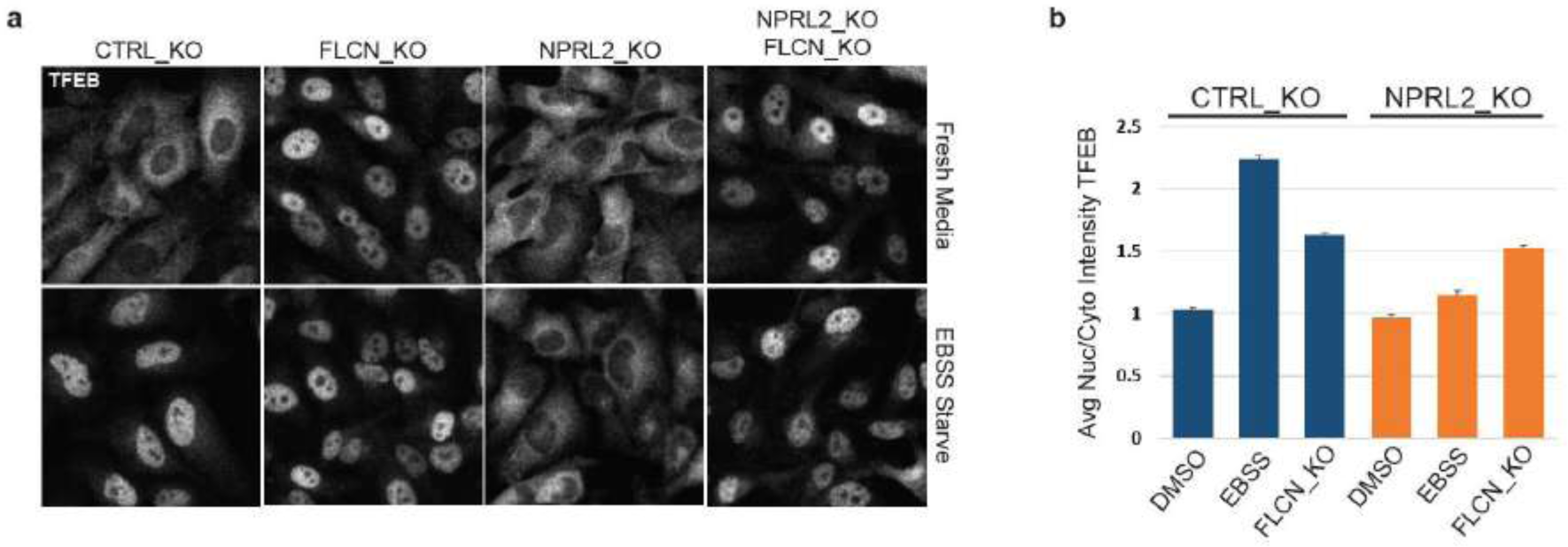
FLCN loss of function results in TFEB nuclear accumulation in WT and NPRL2_KO cells. **(a)** Immunofluorescence of TFEB localization in HeLa cells of the indicated genotypes with or without nutrient starvation (EBSS). **(b)** Quantification of immunofluorescence data from Figure 4I. Cells were starved with EBSS for 60 minutes. Nuclear localization of endogenous TFEB was analyzed by immunofluorescence and high content imaging. Mean ± SD. Minimum of 1500 cells quantified per condition.

**Supplementary figure 13.**
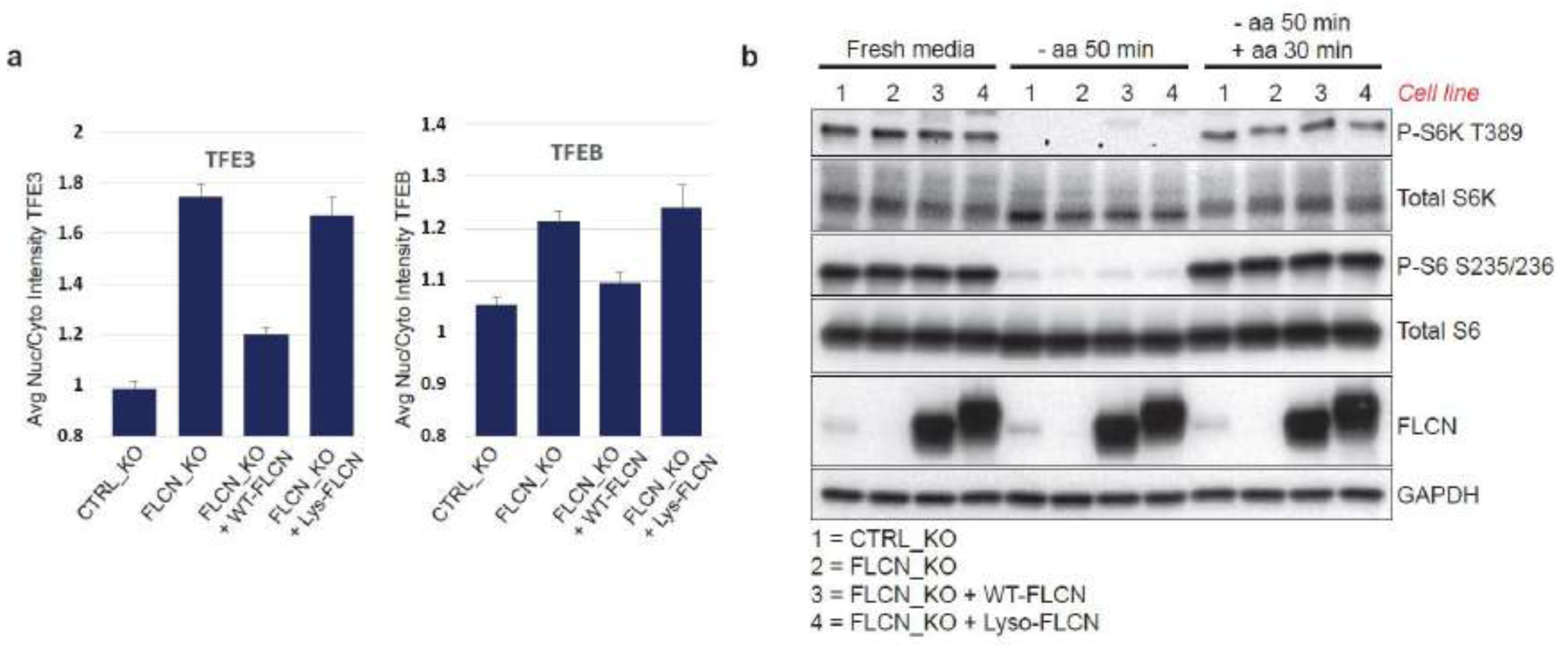
Forced lysosomal recruitment of FLCN activates TFEB/TFE3 in full nutrient conditions independent of mTOR. **(a)** U2OS cells stably expressing Cas9 were genetically engineered to knockout endogenous FLCN and reconstituted with either WT- or lysosome-targeted-FLCN (N-terminal 39aa of LAMTOR1 fused to FLCN N-terminus). Expression of WT-FLCN, but not lyso-FLCN, is able to rescue TFEB/TFE3 cytosolic retention. Mean ± SD. Minimum of 1500 cells quantified per condition. **(b)** FLCN status or localization does not impact mTOR signaling to canonical substrates in response to nutrient modulation, despite regulation of TFEB/TFE3 localization.

**Supplementary figure 14.**
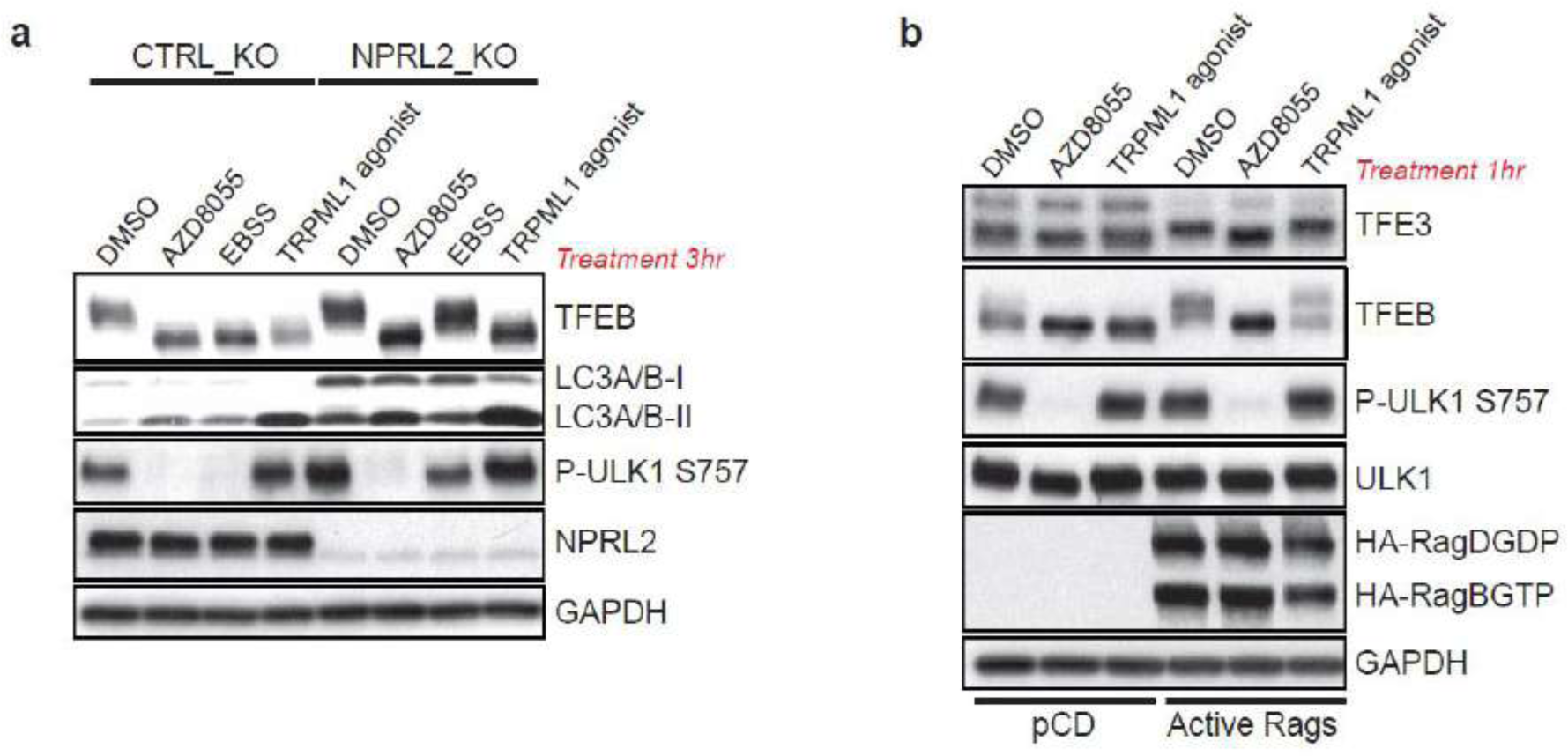
Constitutive RagD^GDP^, but not constitutive RagA^GTP^, suppresses TFEB activation by TRPML1 agonist. **(a)** Western blot analysis of HeLa WT or NPRL2_KO cells treated with the indicated stimuli. Constitutive RagA^GTP^ (LFC-deficiency) suppresses TFEB activation by starvation but not TRPML1 agonist. **(b)** Western blot analysis of HEK293T cells transiently expressing constitutively active RagGTPases (RagB^GTP^/RagD^GDP^) and treated with the indicated compounds for 1hr.

**Supplementary figure 15.**
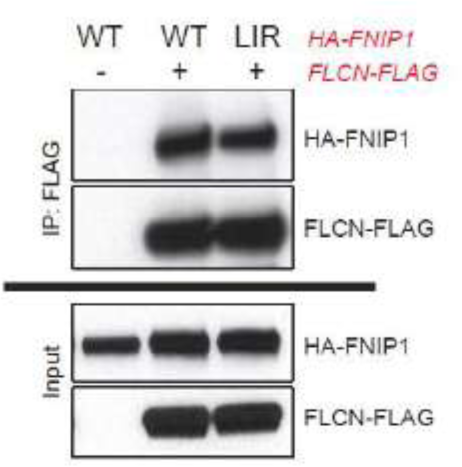
Mutation of FNIP1 LIR motif does not impact binding to FLCN. HEK293T cells were transiently transfected with the indicated constructs 20 hr prior to immunoprecipitation.

**Supplemental figure 16.**
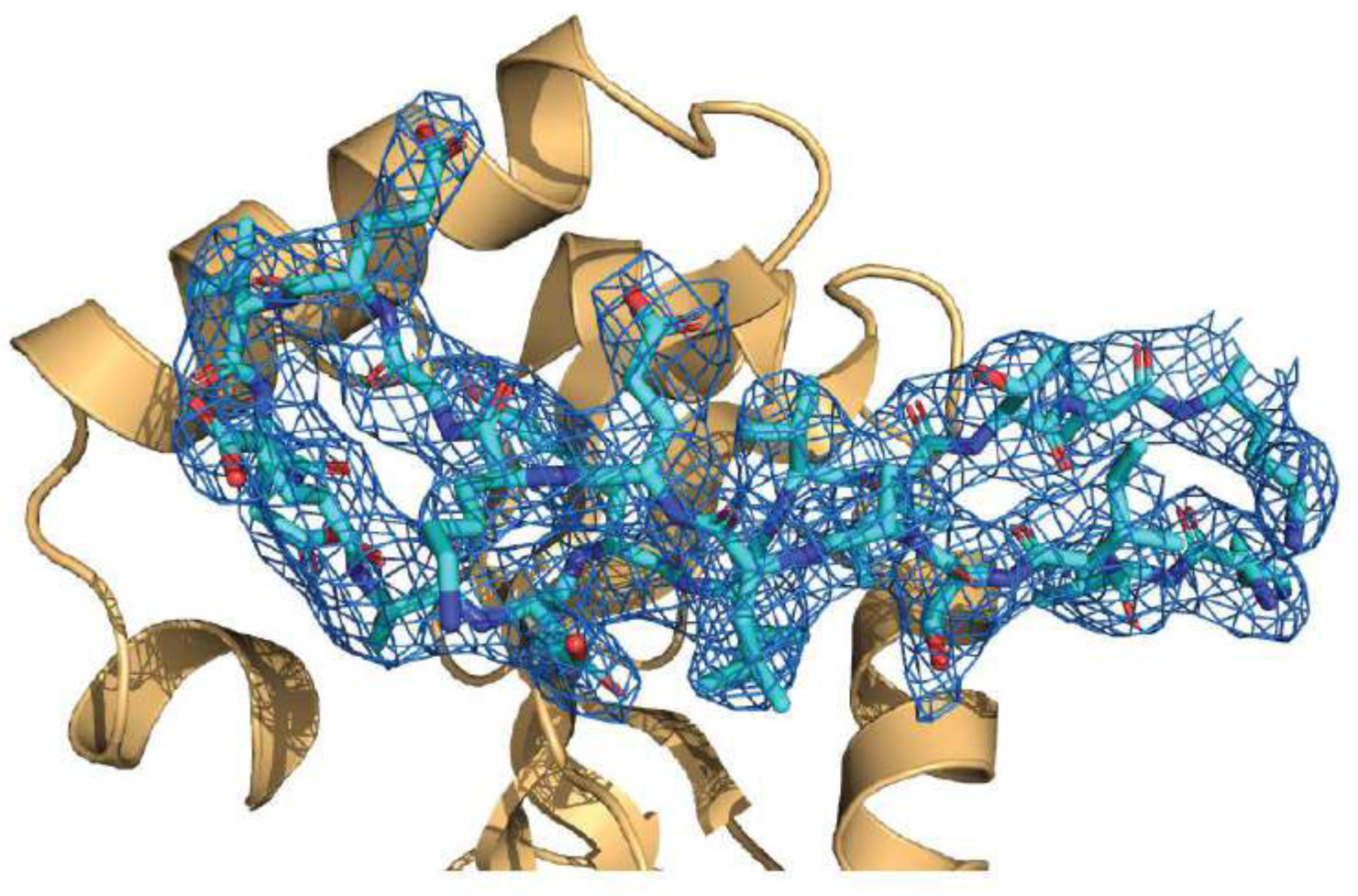
The electron density for the region of FNIP2 is contoured at 1 sigma from the 2Fo-Fc map.

**Supplemental figure 17.**
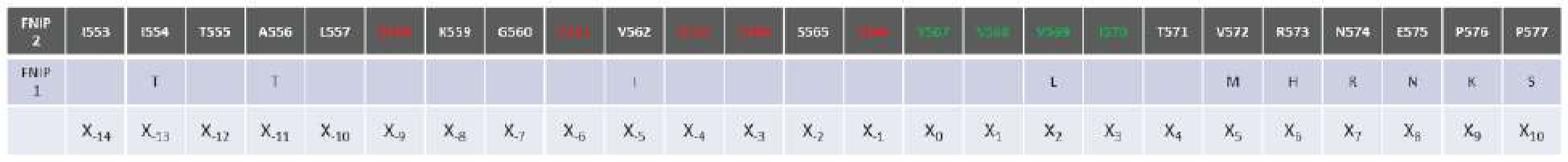
Sequence conservation between FNIP1 and FNIP2 surrounding the identified LIR motif.

**Supplemental figure 18.**
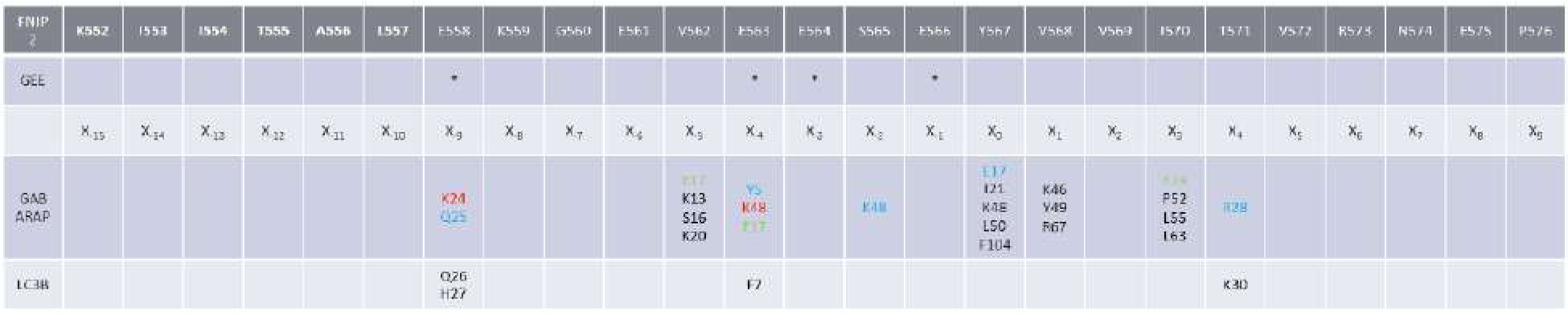
The region spanning 552-576 of FNIP2 was crystallized in complex with GABARAP. Glutamates protected by GEE-labelling are indicated with asterisks. Side chain-mediated interactions on GABARAP are indicated (red, salt bridge; blue, side chain:side chain hydrogen bond; green, main chain: side chain hydrogen bond; black, hydrophobic interactions). Key amino acid differences with LC3B as highlighted in Figure 4G, H are listed.

**Supplemental figure 19.**
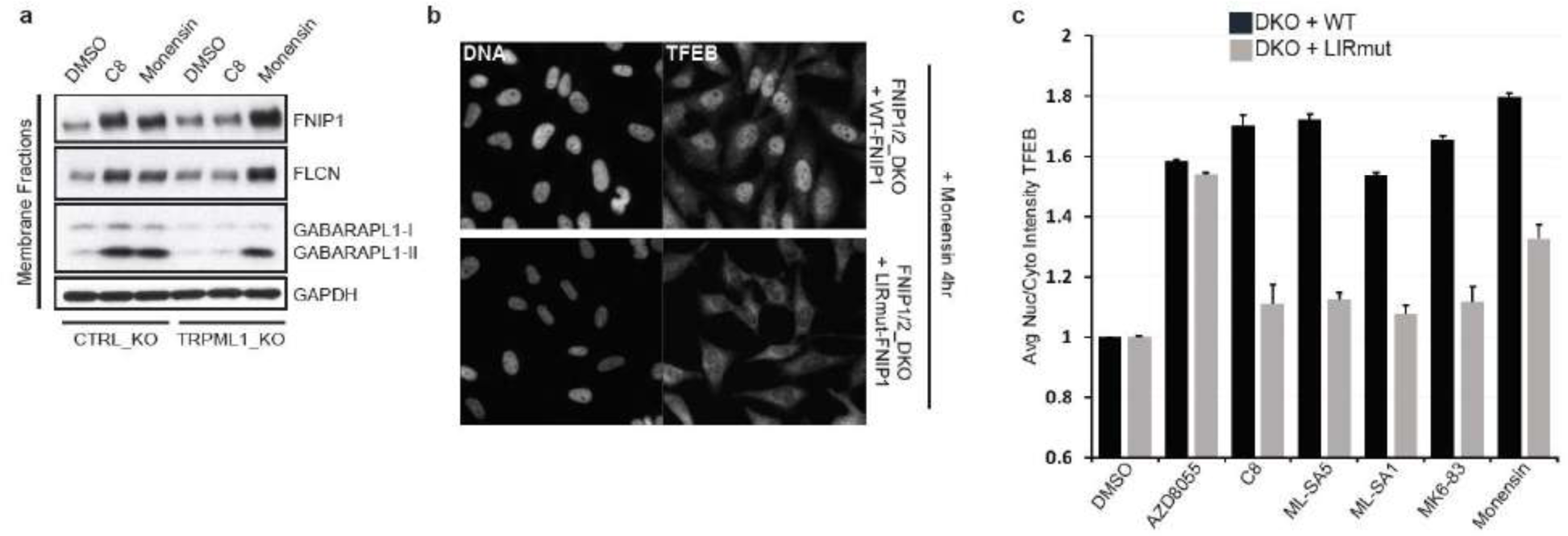
**(a)** Monensin treatment stimulates FLCN-FNIP recruitment to membrane fractions independent of TRPML1. WT or TRPML1_KO HeLa cells were treated with the indicated compounds for 30 min and membrane fractions were analyzed by western blot with the indicated antibodies. C8 (2uM). Monensin (1uM). **(b)** TFEB nuclear localization upon monensin treatment requires FNIP1 LIR-mediated membrane sequestration. Representative images of TFEB nuclear localization upon treatment of the indicated cell lines with monensin (1uM) for 4 hr. (c) Quantification of TFEB nuclear localization upon treatment of the indicated cell lines for 4 hr with the annotated compounds. AZD8055 (2uM), C8 (2uM), ML-SA5 (1uM), MK6-83 (25uM), and monensin (1uM). Mean ± SD. Minimum of 1500 cells quantified per condition.

**Supplementary table 1.** Data collection and refinement statistics (molecular replacement)

|  | Crystal 1 | Crystal 2 |
| --- | --- | --- |
| **Data collection** |  |  |
| Space group | P2(1) |  |
| Cell dimensions |  |  |
| *a*, *b*, *c* (Å) | 75.91, 44.75, 78.48 |  |
| $\alpha$, $\beta$, $\gamma$ (°) | 90.0, 118.5, 90.0 | |
| Resolution (Å) | 39.4 – 1.80 (1.84 – 1.80) * |  |
| *R*<sub>sym</sub> or *R*<sub>merge</sub> | 0.230 (0.716) |  |
| *I* / σ*I* | 10.7 (2.0) | |
| Completeness (%) | 97.5 (98.4) |  |
| Redundancy | 2.8 (2.7) |  |
| **Refinement** |  |  |
| Resolution (Å) | 39.5 – 1.80 |  |
| No. reflections | 39930 |  |
| *R*<sub>work</sub> / *R*<sub>free</sub> | 0.229 / 0.273 |  |
| No. atoms |  |  |
| Protein | 3464 |  |
| Ligand/ion | 0 |  |
| Water | 408 |  |
| *B*-factors |  |  |
| Protein | 18.9 |  |
| Ligand/ion | 0 |  |
| Water | 35.2 |  |
| R.m.s. deviations |  |  |
| Bond lengths (Å) | 0.012 |  |
| Bond angles (°) | 1.69 |  |

\*Values in parentheses are for highest-resolution shell.

**Supplementary table 2.**
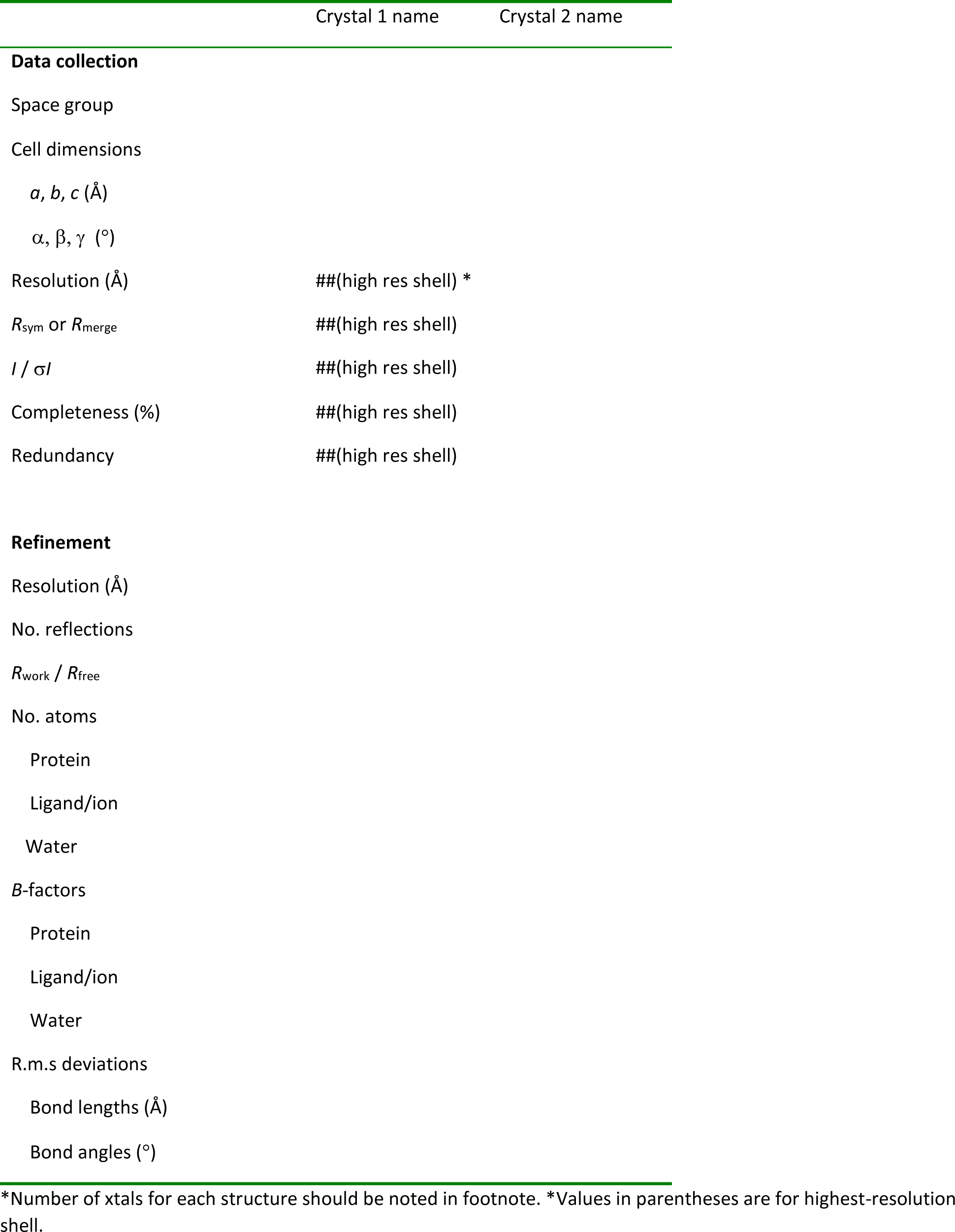
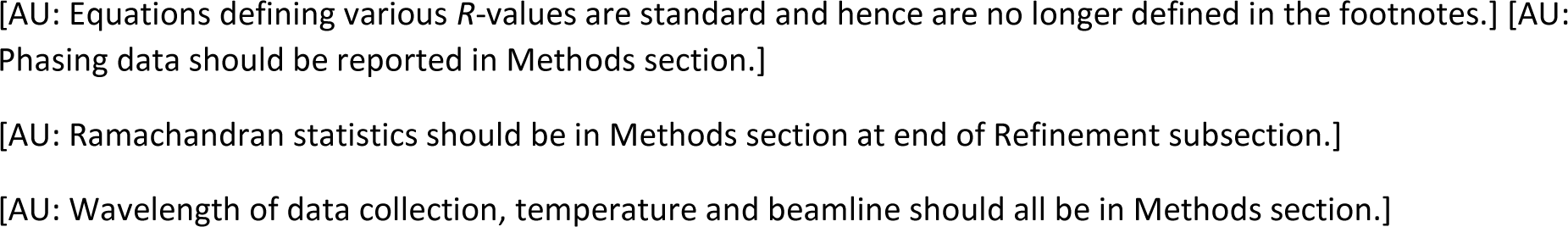
Data collection, phasing and refinement statistics (MIR)

**Supplementary table 3.**
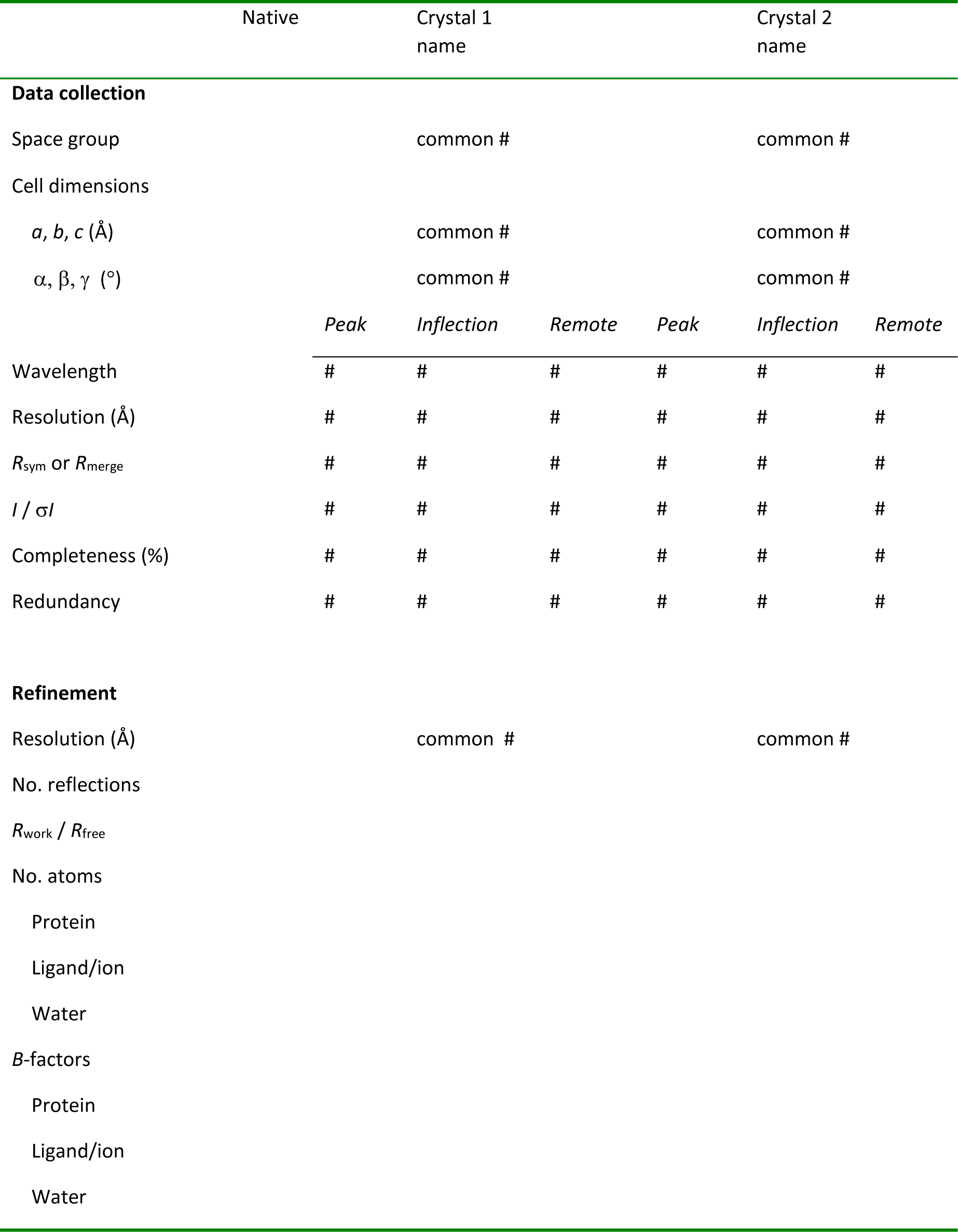
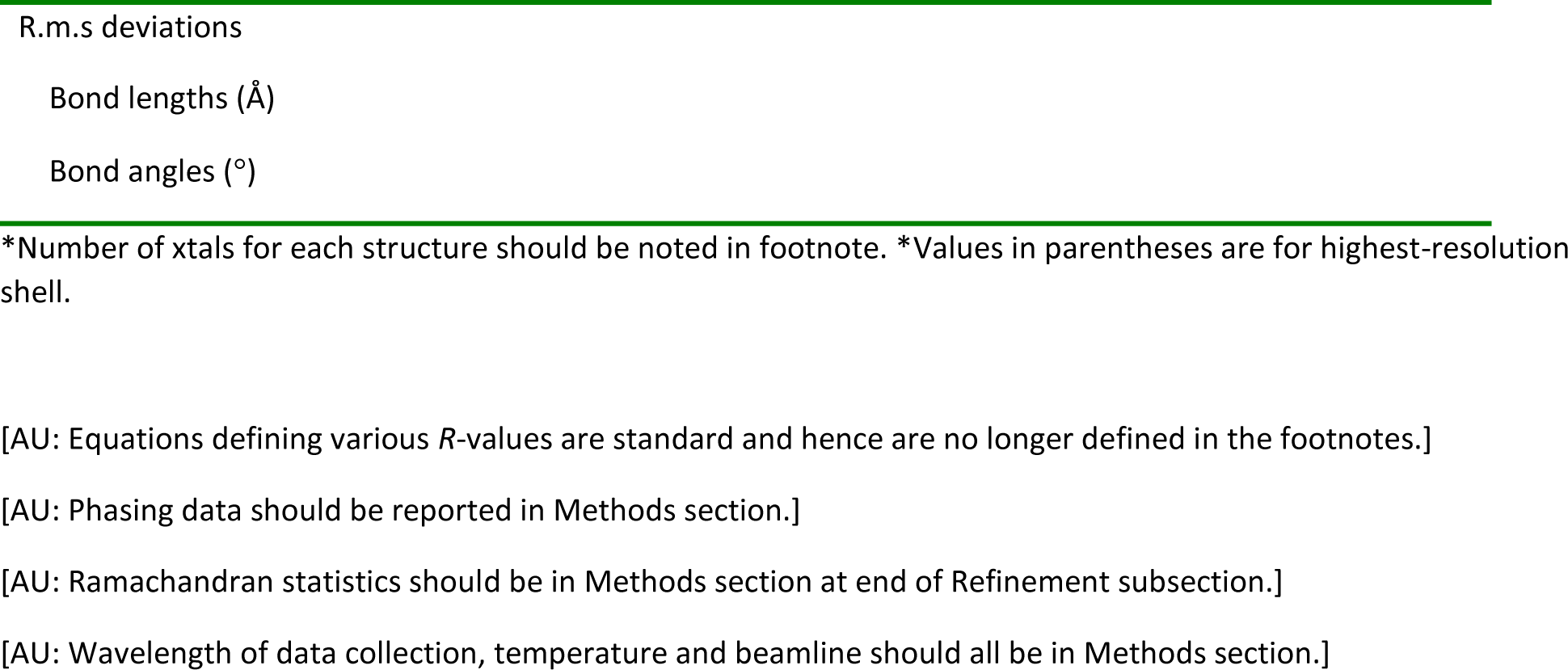
Data collection, phasing and refinement statistics for MAD (SeMet) structures.

**Supplementary table 4.**
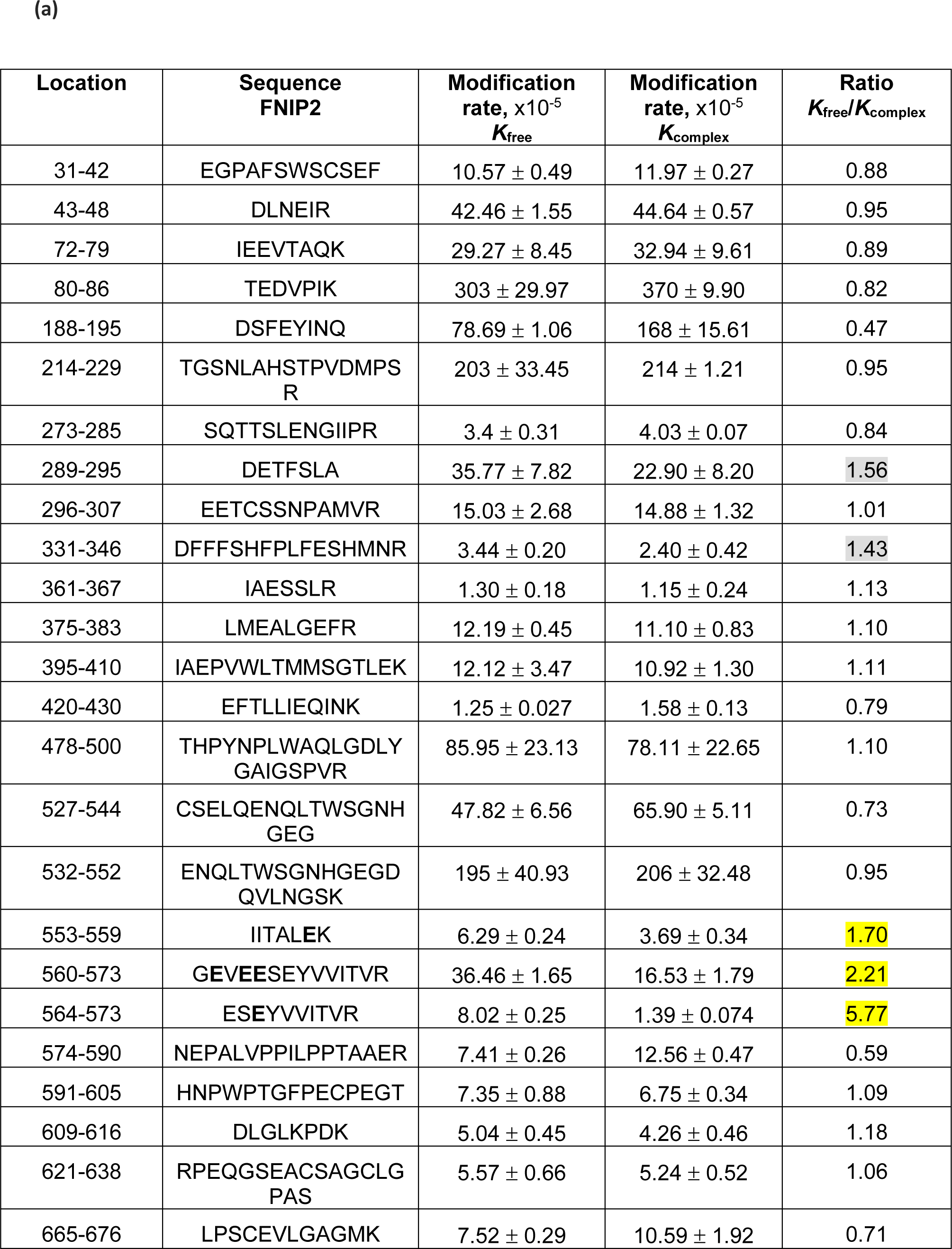
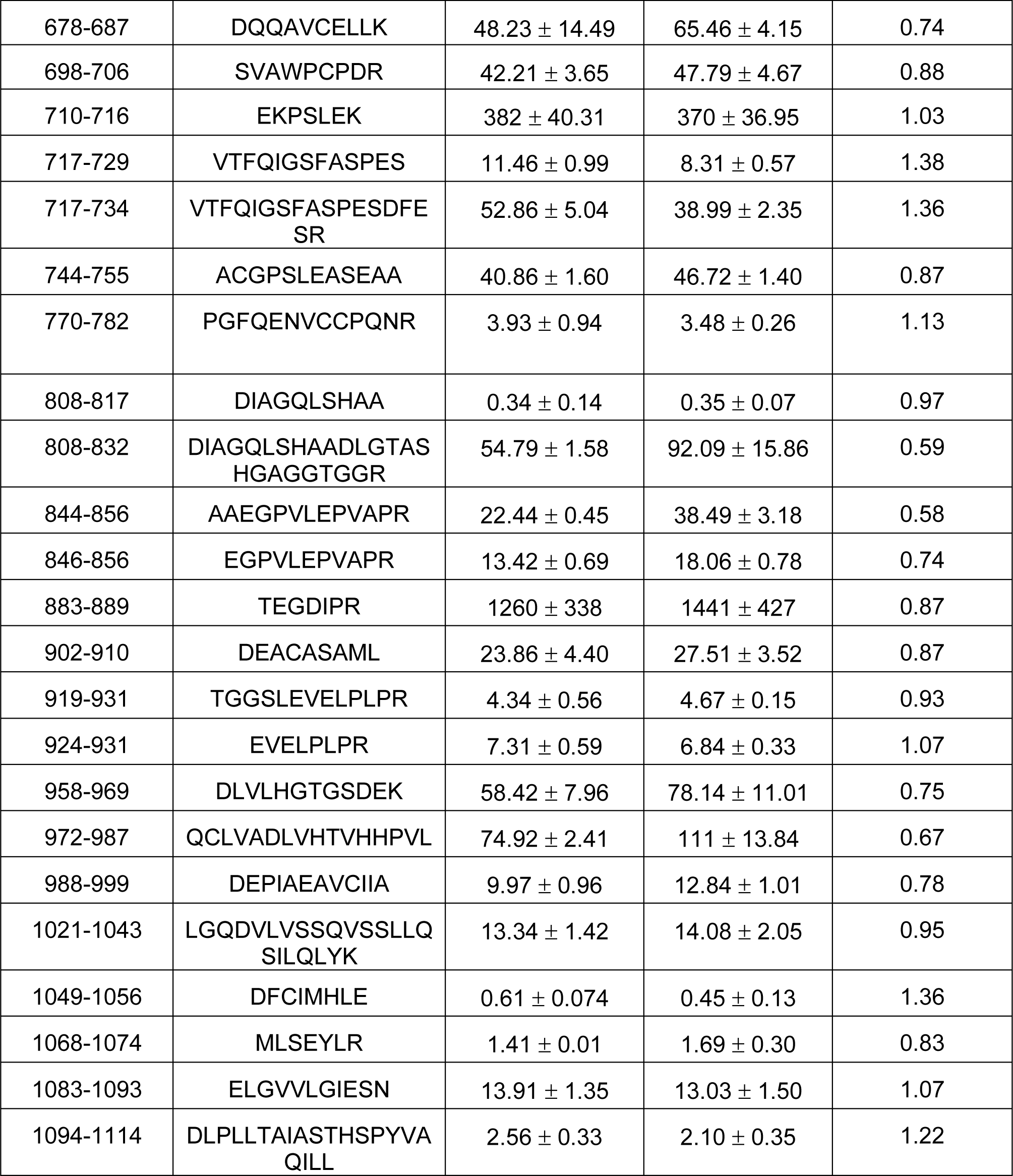
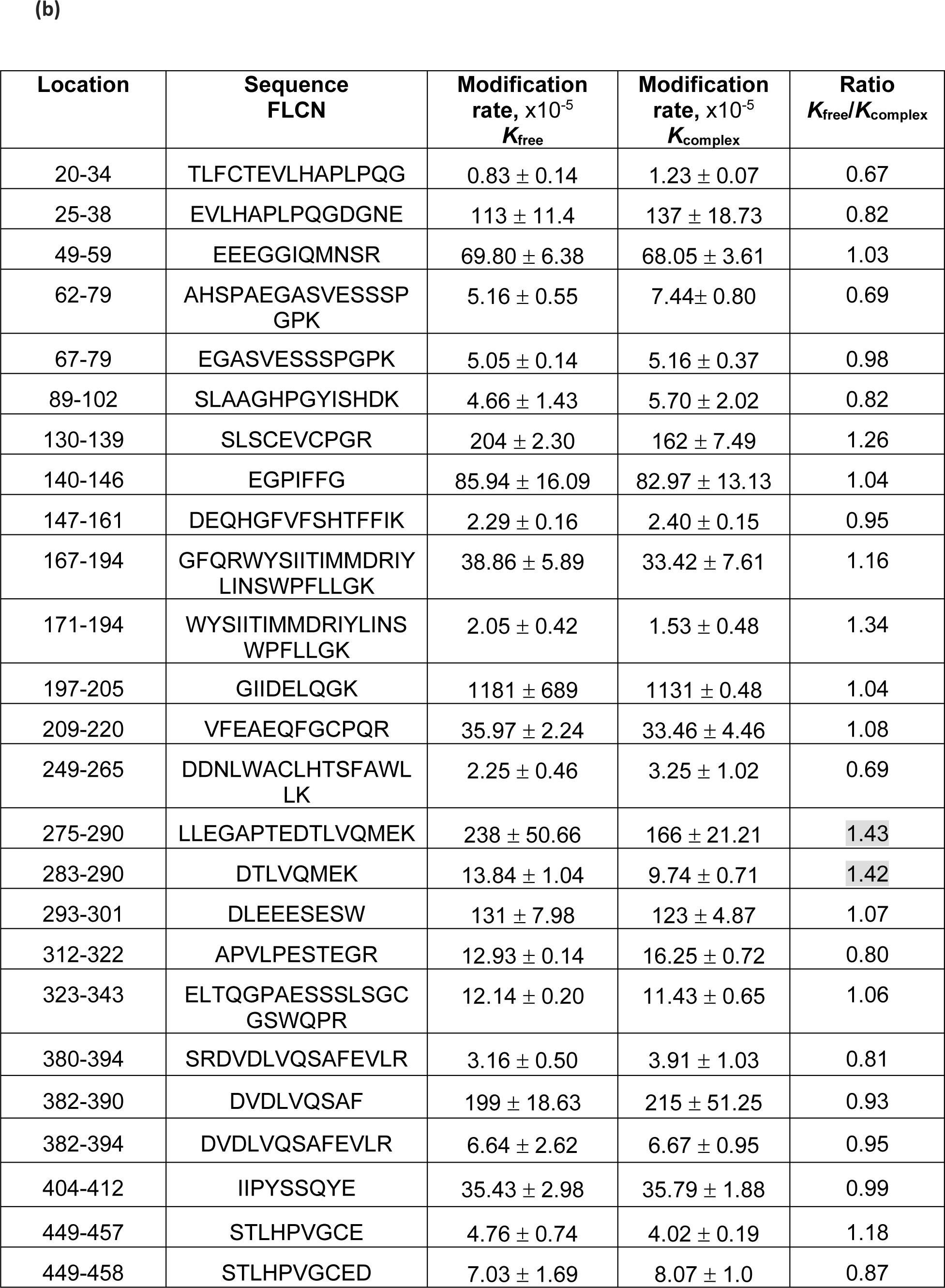
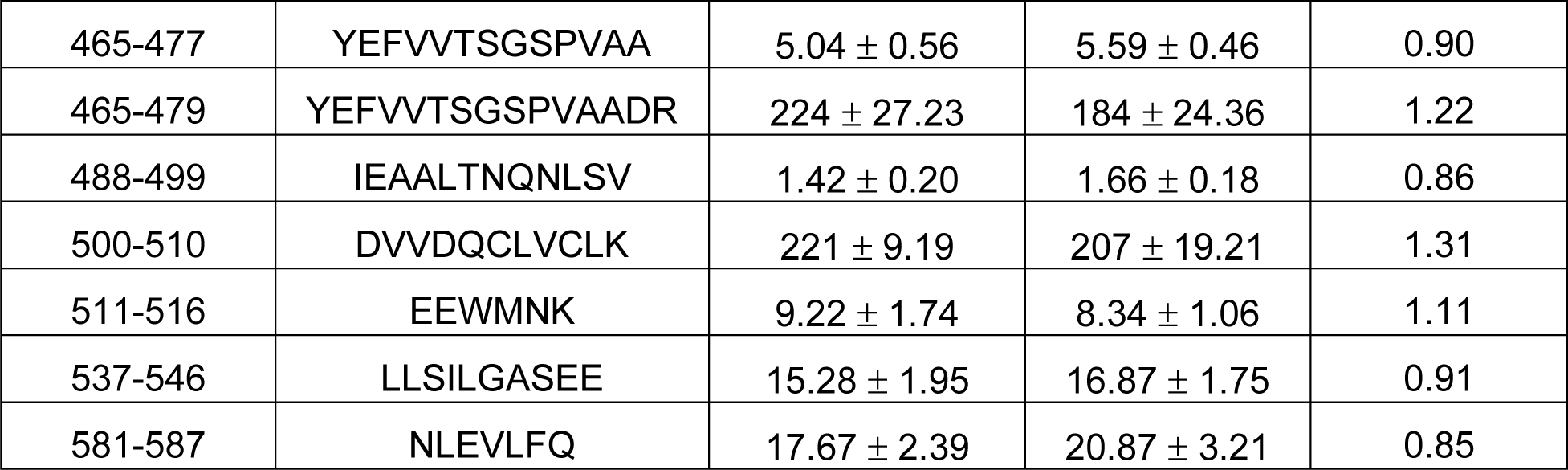
Rate constants for the modified peptides of FNIP2 and FLCN identified by carboxyl group Footprinting. The highest protection ratios (R) are highlighted in yellow for peptides derived from (a) FNIP2 and (b) FLCN. Modestly protected peptides are highlighted in grey. Three peptides from FNIP2 that cover 553-559, 560-573 and 564-573 exhibited the highest protection ratios of 1.70, 2.21 and 5.77, respectively, while two peptides covering 286-295 and 331-346 showed more modest protection ratios of 1.56 and 1.43, respectively. For FLCN, two peptides covering 275-290 and 283-290 showed modest protection ratios of 1.43 and 1.42, respectively. MS data from one replicate experiment were used to calculate *K* values for each peptide. The overall fit results for all detected peptides within (a) FNIP2 and (b) FLCN are shown. Peptide locations and their corresponding sequences are shown in columns 1 and 2. The third and fourth columns denote the *K* values for free FLCN/FNIP2 (*K*_free_) and the complex with GABARAP (*K*_complex_), respectively. The fifth column shows the ratio, R = *K*_free_/*K*_complex_. For a given peptide, R<1 suggests that the corresponding region experienced a gain in solvent accessibility due to structural changes introduced during complex formation. A R value close to 1 indicates that the solvent accessibility of the region remains unchanged, while a R>1 suggests that the corresponding region exhibits protection from the solvent as a function of the complex formation. The R values for all of the peptides (column five) fell between 0.47 and 5.77 with a mean value of 1.06 and a median value of 0.95. The histogram of the distribution of R values (Fig. 3B) for all peptides also indicated that a majority of the peptides within FLCN/FNIP2 exhibited changes in modification upon complex formation close to 1. Using a strategy similar to one used in metabolomics to correct for non-biological variations between samples (mean scaling, division by central tendency), we used the average of the mean and the median to normalize the ratios to 1. A normalization factor for these studies was calculated to be 1 = ((1.05+0.95)/2).

**Supplementary table 5.**
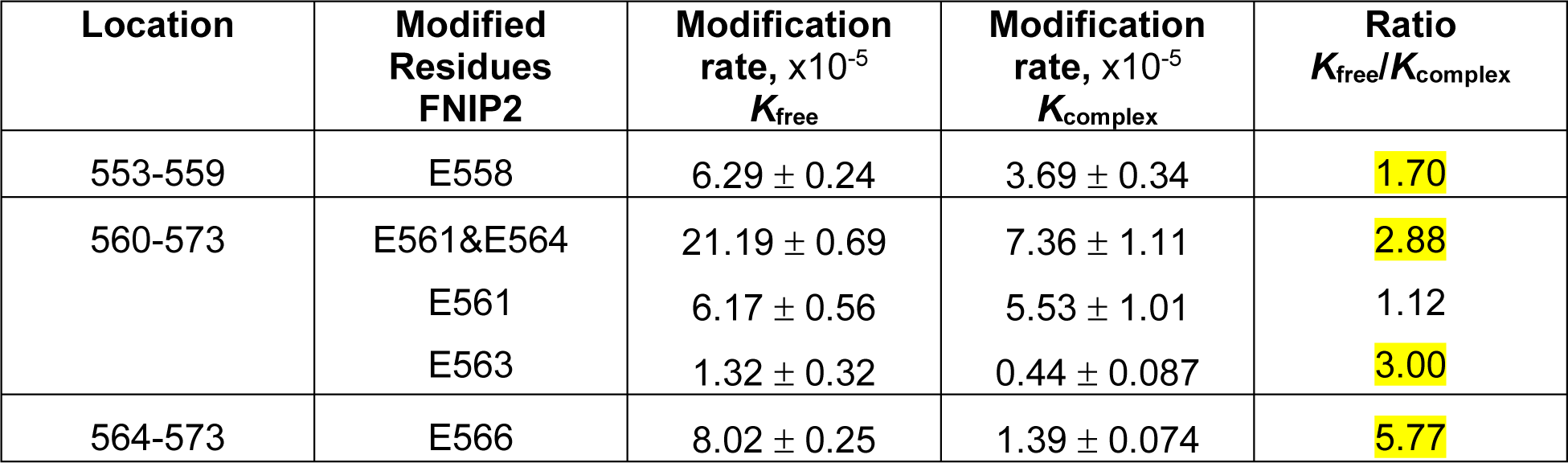
Rate constants for the three most modified residues in FNIP2. Residues E558, E564, E563 and E566 (within peptides 553-559, 560-573 and 564-573) showed the highest protection of 1.70, 2.88, 3.0 and 5,77, respectively, upon complex formation Peptides were derived from trypsin/Asp-N digestion.

## References and notes

1. Dikic, I. Proteasomal and autophagic degradation systems. Ann Rev Biochem. 86, 193– 224 (2017).

2. Lawrence, R.E., and R. Zoncu. The lysosome as a cellular centre for signalling, metabolism and quality control. Nat Cell Biol. 21, 133–142 (2019).

3. Kaur, J., and J. Debnath, Autophagy at the crossroads of catabolism and anabolism. Nat Rev Mol Cell Biol. 16, 461–72 (2015).

4. Ballabio, A., and J.S. Bonifacino. Lysosomes as dynamic regulators of cell and organismal homeostasis. Nat Rev Mol Cell Biol. 21, 101–118 (2020).

5. Florey, O., Kim, S.E., Sandoval, C.P., Haynes, C.M., and M. Overholtzer. Autophagy machinery mediates macroendocytic processing and entotic cell death by targeting single membranes. Nat Cell Biol. 13, 1335–43 (2011).

6. Martinez, J.S., Almendinger, J., Oberst, A., Ness, R., Dillon, C.P., Fitzgerald, P., Hengartner, M.O., and D.R. Green. Microtubule-associated protein 1 light chain 3 alpha (LC3)-associated phagocytosis is required for the efficient clearance of dead cells*. Proc. Nat* Acad. Sci. 108, 17396–401 (2011).

7. Jacquin, E., Leclerc-Mercier, S., Judon, C., Blanchard, E., Fraitag, S., and O. Florey. Pharmocological modulators of autophagy activate a parallel noncanonical pathway driving unconventional LC3 lipidation. Autophagy. 13, 854–867 (2017).

8. Heckmann, B.L., Teubner, B.J.W., Tummers, B., Boada-Romero, E., Harris, L., Yang, M., Guy, C.S., Zakharenko, S.S., and D.R. Green. LC3-associated endocytosis facilitates β-amyloid clearance and mitigates neurodegeneration in murine Alzheimer’s disease. Cell. 178, 536–51 (2019).

9. Kaushik, S., and A.M. Cuervo. The coming of age of chaperone-mediated autophagy. Nat Rev Mol Cell Biol. 19, 365–381 (2018).

10. Honey, K., and A.Y. Rudensky. Lysosomal cysteine proteases regulate antigen presentation. Nat Rev Immunol. 3, 472–482 (2003).

11. Buratta, S., Tancini, B., Sagini, K., Delo, F., Chiaradia, E., Urbanelli, L., and C. Emiliani. Lysosomal exocytosis, exosome release, and secretory autophagy: the autophagic- and endo-lysosomal systems go extracellular. Int J Mol Sci. 21, 2576 (2020).

12. Xu, H. and D. Ren. Lysosomal physiology. Annu Rev Physiol. 77, 57–80 (2015).

13. Liu, G.Y., and D.M. Sabatini. mTOR at the nexus of nutrition, growth, ageing and disease. Nat Rev Mol Cell Biol. 21, 183–203 (2020).

14. Fromm, S.A., Lawrence, R.E., and J.H. Hurley. Structural mechanism for amino acid-dependent Rag GTPase nucleotide state switching by SLC38A9. Nat Struct Mol Biol. 27, 1017–1023 (2020).

15. Kim, J., and K-L. Guan. mTOR as a central hub of nutrient signalling and cell growth. Nat Cell Biol. 21, 63–71 (2019).

16. Settembre, C., Fraldi, A., Medina, D.L., and A. Ballabio. Signals from the lysosome: a control centre for cellular clearance and energy metabolism. Nat Rev Mol Cell Biol. 14, 283–96 (2013).

17. Sardiello, M., Palmieri, M., di Ronza, A., Medina, D.L., Valenza, M., Gennarino, V.A., Di Malta, C., Donaudy, F., Embrione, V., Polishchuk, R.S., et al., A gene network regulating lysosomal biogenesis and function. Science. 325, 473–7 (2009).

18. Martina, J.A., Diab, H.I., Lishu, L., Jeong-A, L., Patange, S., Raben, N., and R. Puertollano. The nutrient-responsive transcription factor TFE3 promotes autophagy, lysosomal biogenesis, and clearance of cellular debris. Sci Signal. 7, ra9 (2014).

19. Settembre, C., Di Malta, C., Polito, V.A., Garcia Arencibia, M., Vetrini, F., Erdin, S., Erdin, S.U., Huynh, T., Medina, D., Colella, P., et al. TFEB links autophagy to lysosomal biogenesis. Science. 332, 1429–33 (2011).

20. Lawrence, R.E., Fromm, S.A., Fu, Y., Yokom, A.L., Kim, D.J., Thelen, A.M., Young, L.N., Lim, C-Y., Samelson, A.J., Hurley, J.H., and R. Zoncu. Structural mechanism of a Rag GTPase activation checkpoint by the lysosomal folliculin complex. Science. 366, 971–977 (2019).

21. Shen, K., Rogala, K.B., Chou, H-T., Huang, R., Yu, Z., and D.M. Sabatini. Cryo-EM structure of the human FLCN-FNIP2-Rag-Ragulator complex. Cell. 179, 1319–1329 (2019).

22. Napolitano, G., Di Malta, C., Esposito, A., de Araujo, M.E.G., Pece, S., Bertalot, G., Matarese, M., Benedetti, V., Zampelli, A., Stasyk, T., et al., A substrate-specific mTORC1 pathway underlies Birt-Hogg-Dube syndrome. Nature. 585, 597–602 (2020).

23. Settembre, C., Zoncu, R., Medina, D.L., Vetrini, F., Erdin, S., Erdin, SU., Huynh, T., Ferron, M., Karsenty, G., Vellard, M.C., et al., A lysosome-to-nucleus signalling mechanism senses and regulates the lysosome via mTOR and TFEB. EMBO J. 31, 1095–1108 (2012).

24. Galluzzi, L., and D.R., Green, Autophagy-independent functions of the autophagy machinery. Cell. 177, 1682–99 (2019).

25. Xu, Y., Zhou, P., Cheng, S., Lu, Q., Nowak, K., Hopp, A-K., Li, L., Shi, X., Zhou, Z., Gao, W., et al., A bacterial effector reveals the V-ATPase-ATG16L1 axis that initiates xenophagy. Cell. 178, 1–15 (2019).

26. Fletcher, K., Ulferts, R., Jacquin, E., Veith, T., Gammoh, N., Arasteh, J.M., Mayer, U., Carding. S.R., Wileman, T., Beale, R., and O. Florey. The WD40 domain of ATG16L1 is required for its non-canonical role in lipidation of LC3 at single membranes. EMBO J. 37, e97840 (2018).

27. Nakamura, S., Shigeyama, S., Minami, S., Shima, T., Akayama, S., Matsuda, T., Esposito, A., Napolitano, G., Kuma, A., Namba-Humano, T., et al., LC3 lipidation is essential for TFEB activation during the lysosomal damage response to kidney injury. Nat Cell Biol. 22, 1252–1263 (2020).

28. Kumar, S., Jain, A., Choi, S.W., Duarte da Silva, G.P., Allers, L., Mudd, M.H., Peters, R.S., Anonsen, J.H., Rusten, T-E., Lazarou, M., and V. Deretic. Mammalian Atg8 proteins and the autophagy factor IRGM control mTOR and TFEB at a regulatory node critical for responses to pathogens. Nat Cell Biol. 22: 973–985 (2020).

29. Chen, C-C., Keller, M., Hess, M., Schiffmann, R., Urban, N., Wolfgardt, A., Schaefer, M., Bracher, F., Biel, M., Wahl-Scott, C., and C. Grimm. A small molecule restores function to TRPML1 mutant isoforms responsible for mucolipidosis type IV. Nat Comm., 5, 1–10 (2014).

30. Shen, D., Wang, X., Li, X., Zhang, X., Yao, Z., Dibble, S., Dong, X-P., Yu, T., Lieberman, A.P., Showalter, H.D., and H. Xu. Lipid storage disorders block lysosomal trafficking by inhibiting a TRP channel and lyosomal calcium release. Nat Comm. 3, 1–11 (2012).

31. Liang., C., Piperazine derivatives as trpml modulators. WO 2018/005713 A1. Jan 4 (2018).

32. Chresta, C.M., Davies, B.R., Hickson, I., Harding, T., Cosulich, S., Critchlow, S.E., Vincent, J.P., Ellston, R., Jones, D., Sini, P., et al., AZD8055 is a potent, selective, and orally bioavailable ATP-competitive mammalian target of rapamycin kinase inhibitor with in vitro and in vivo antitumor activity. Cancer Res. 70, 288–98 (2010).

33. Florey, O., Gammoh, N., Kim, S.E., Jiang, X., and M. Overholtzer. V-ATPase and osmotic imbalances activate endolysosomal LC3 lipidation. Autophagy. 11, 88–99 (2015).

34. Medina, D.L., Di Paola, S., Peluso, I., Armani, A., De Stefani, D., Venditti, R., Montefusco, S., Scotto-Rosato, A., Prezioso, C., Forrester, A., et al., Lysosomal calcium signaling regulates autophagy through calcineurin and TFEB. Nat Cell Biol. 17, 288–99 (2015).

35. Johansen, T., and T., Lamark. Selective autophagy: ATG8 family proteins, LIR motifs and cargo receptors. J Mol Biol. 432, 80–103 (2020).

36. Dunlop, E.A., Seifan, S., Claessens, T., Behrends, C., Af Kamps, M., Rozycka, E., Kemp, A.J., Nookala, R.K., Blenis, J., Coull, B.J., et al., FLCN, a novel autophagy component, interacts with GABARAP and is regulated by ULK1 phosphorylation. Autophagy. 10, 1749–1760 (2014).

37. Hong, S.B., Oh, H.B., Valera, V.A., Baba, M., Schmidt, L.S., and W.M. Linehan. Inactivation of the FLCN tumor suppressor gene induces TFE3 transcriptional activity by increasing its nuclear translocation. PLoS One. 5, e15793 (2010).

38. Petit, C.S., Roczniak-Ferguson, A., and S.M. Ferguson. Recruitment of folliculin to lysosomes supports the amino acid-dependent activation of Rag GTPases. J Cell Biol. 202, 1107–1122 (2013).

39. Martina, J.A., and R., Puertollano. Rag GTPases mediate amino acid-dependent recruitment of TFEB and MITF to lysosomes. J Cell Biol. 200, 475–91 (2013).

40. Wirth, M., Zhang, W., Razi, M., Nyoni, L., Joshi, D., O’Reilly, N., Johansen, T., Tooze, S.A., and S. Mouilleron. Molecular determinants regulating selective binding of autophagy adapters and receptors to ATG8 proteins. Nat Comm. 10, 2055 (2019).

41. Meng, J., and S.M., Ferguson. GATOR1-dependent recruitment of FLCN-FNIP to lysosomes coordinates Rag GTPase heterodimer nucleotide status in response to amino acids. J Cell Biol. 217, 2765–2776 (2018).

42. Manifava, M., Smith, M., Rotondo, S., Walker, S., Niewczas, I., Zoncu, R., Clark, J., and N.T. Ktistakis. Dynamics of mTORC1 activation in response to amino acids. eLife. 5:e19960 (2016).

43. Lawrence, R.E., Cho, K.F., Rappold, R., Thrun, A., Tofaute, M., Kim, D.J., Moldavski, O., Hurley, J.H., and R. Zoncu. A nutrient-induced affinity switch controls mTORC1 activation by its Rag GTPase-Ragulator lysosomal scaffold. Nat Cell Biol. 20, 1052–1063 (2018).

44. Parminder, K., Tomechko, S.E., Kiselar, J., Shi, W., Deperalta, G., Wecksler, A.T., Gokulrangan, G., Ling, V., and M.R. Chance. Characterizing monoclonal antibody structure by carboxyl group footprinting. MAbs. 7, 540–52 (2015).

45. Zhang, H., Wen, J., Hunag, R. Y-C., Blankenship, R.E., and M.L. Gross. Mass spectrometry-based carboxyl footprinting of proteins: method evaluation. Int J Mass Spectrom. 312, 78– 86 (2012).

46. Noda, N.N., Ohsumi, T., and F. Inagaki. ATG8-family interacting motif crucial for selective autophagy. FEBS Lett. 584, 1379–1385 (2010).

47. Nezich, C.L., Wang, C., Fogel, A.I., and R.J. Youle. MiT/TFE3 transcription factors are activated during mitophagy downstream of Parkin and ATG5. J Cell Biol. 210, 435–50 (2015).

48. Heo, J-M., Ordureau, A., Swarup, S., Paulo, J.A., Shen, K., Sabatini, D.M., and J.W. Harper. RAB7A phosphorylation by TBK1 promotes mitophagy via the PINK-PARKIN pathway. Sci Adv. 4, eaav0443 (2018).

49. Lazarou, M., Sliter, D.A., Kane, L.A., Sarraf, S.A., Wang, C., Burman, J.L., Sideris, D.P., Fogel, A.I, and R.J. Youle. The ubiquitin kinase PINK1 recruits autophagy receptors to induce mitophagy. Nature. 524, 309–314 (2015).

50. Birmingham, C.L., Smith, A.C., Bakowski, M.A., Yoshimori, T., and J.H. Brumell. Autophagy controls *Salmonella* infection in response to damage to the *Salmonella*-containing vacuole. J Biol Chem. 281, 11374–11383 (2006).

51. Tsuboyama, K., Koyama-Honda, I., Sakamaki, Y., Koike, M., Morishita, H., and N. Mizushima. The ATG conjugation systems are important for degradation of the inner autophagosomal membrane. Science. 354, 1036–1041 (2016).

52. Schmidt, L.S., and W.M. Linehan. FLCN: The causative gene for Birt-Hogg-Dube syndrome. Gene. 640, 28–42 (2018).

53. Perera, R.M., Stoykova, S., Nicolay, B.N., Ross, K.N., Fitament, J., Boukhali, M., Lengrand, J., Deshpande, V., Selig, M.K., Ferrone, C.R., et al. Transcriptional control of autophagy-lysosome function drives pancreatic cancer metabolism. Nature. 524, 361–5 (2015).

54. Gupta, S., Yano, J., Htwe, H.H., Shin, H.R., Cakir, Z., Ituarte, T., Wen, K.W., Kim, G.E., Zoncu, R., Dawson, D.D., and R.M. Perera. Lysosomal retargeting of Myoferlin mitigates membrane stress to enable pancreatic cancer growth. bioRxiv. doi.org/10.1101/2021.01.04.425106 (2021).

55. Koster, S., Upadhyay, S., Chandra, P., Papavinasasundaram, K., Yang, G., Hassan, A., Grigsby, S.J., Mittal, E., Park, H.S., Jones, V., et al., Mycobacterium tuberculosis is protected from NADPH oxidase and LC3-associated phagocytosis by the LCP protein CpsA. Proc Natl Acad Sci.114, E8711–E8720 (2017).

56. Choy, A., Dancourt, J., Mugo, B., O’Connor, T.J., Isberg, R.R., Melia, T.J., and C.R. Roy. The Legionella effector RavZ inhibits host autophagy through irreversible Atg8 deconjugation. Science. 338, 1072–6 (2012).

57. Visvikis, O., Ihuegbu, N., Labed, S.A., Luhachack, L.G., Alves, A-M.F., Wollenberg, A.C., Stuart, L.M., Stormo, G.D., and J.E. Irazoqui. Innate host defense requires TFEB-mediated transcription of cytoprotective and antimicrobial genes. Immunity. 40, 896–909 (2014).

58. Sanjuan, M.A., Dillon, C.P., Tait, S.W.G., Moshiach, S., Dorsey, F., Connell, S., Komatsu, M., Tanaka, K., Cleveland, J.L., Withoff, S., and D.R. Green. Toll-like receptor signalling in macrophages links the autophagy pathway to phagocytosis. Nature. 450, 1253–7 (2007).

59. Martinez, J., Subbarao Malireddi, R.K., Lu, Q., Dias Cunha, L., Pelletier, S., Gingras, S., Orchard, R., Guan, J.L., Tan, H., Peng, J., et al. Molecular characterization of LC3-associated phagocytosis reveals distinct roles for Rubicon, NOX2 and autophagy proteins. Nat Cell Biol. 17, 893–906 (2015).

60. Napolitano, G., Esposito, A., Choi, H., Matarese, M., Benedetti, V., Di Malta, C., Monfregola, J., Medina, D.L., Lippincott-Schwartz, J., and A. Ballabio. mTOR-dependent phosphorylation controls TFEB nuclear export. Nat Comm. 9, 3312 (2018).

61. Heckmann, B.L., Teubner, B.J.W., Boada-Romero, E., Tummers, B., Guy, C., Fitzgerald, P., Mayer, U., Carding, S., Zakharenko, S.S., Wileman, T., and D.R. Green. Noncanonical function of an autophagy protein prevents spontaneous Alzheimer’s disease. Sci Adv. 6, eabb9036 (2020).

62. Bonam, S.R., Wang, F., and S. Muller. Lysosomes as a therapeutic target. Nat Rev Drug Disc. 18, 923–948 (2019).

## Supplemental References

63. Abu-Remaileh, M., et al., Lysosomal metabolomics reveals v-ATPase and mTOR-dependent regulation of amino acid efflux from lysosomes. Science. 358: 807–813 (2017).

64. Brumell, J.H., Rosenberger, C.M., Gotto, G.T., Marcus, S.L., and B.B., Finlay. SifA permits survival and replication of Salmonella typhimurium in murine macrophages. Cell Microbiol. 3, 75–84 (2001).

65. Kishi-Itakura, C., et al., Ultrastructural analysis of autophagosome organization using mammalian autophagy-deficient cells. J Cell Sci. 127, 4089–4102 (2014).

66. Kishi-Itakura, C., and F., Buss. The use of correlative light-electron microscopy (CLEM) to study PINK1/Parkin-mediated mitophagy. Methods Mol Biol. 1759, 29–39 (2018).

67. McCarthy, D.J., et al., Differential expression analysis of multifactor RNA-seq experiments with respect to biological variation. Nuc Acid Res. 40, 4288–97 (2012).

68. Bouziat, R., et al., Reovirus infection triggers inflammatory responses to dietary antigens and development of celiac disease. Science. 356, 44–50 (2017).

69. Bouziat, R., et al., Murine norovirus infection induces TH1 inflammatory responses to dietary antigens. Cell Host Microbe. 24, 677–688 (2018).

70. DeJesus, R., et al., Functional CRISPR screening identifies the ufmylation pathway as a regulator of SQSTM1/p62. eLife. 5, e17290 (2016).

71. Lystad, A.H., et al., Distinct functions of ATG16L1 isoforms in membrane binding and LC3B lipidation in autophagy-related processes. Nat Cell Biol. 21, 372–383 (2019).

72. Turco, E., et al., FIP200 Claw domain binding to p62 promotes autophagosome formation at ubiquitin condensates. Mol Cell. 74, 330–46 (2019).

73. Kabsch W. XDS. Acta Crystallogr D Biol Crystallogr. 66, 125–32. doi: 10.1107/S0907444909047337 (2010).

74. Evans, P.R., and G.N. Murshudov., How good are my data and what is the resolution? Acta Crystallogr D Biol Crystallogr. 69, 1204–14 (2013).

75. McCoy, A.J., et al., Phaser crystallographic software. J Appl Crystallogr. 40, 658–674 (2007).

76. Murshudov, G.N., et al., REFMAC5 for the refinement of macromolecular crystal structures. Acta Crystallogr D Biol Crystallogr. 2011 Apr;67, 355–67 (2011).

77. Adams, P.D., et al., PHENIX: a comprehensive Python-based system for macromolecular structure solution. Acta Crystallogr D Biol Crystallogr. 66, 213–21 (2010).

78. Emsley, P., et al., Features and development of Coot. Acta Crystallogr D Biol Crystallogr. 66, 486–501 (2010).

79. Nieba, L., et al., Competition BIAcore for measuring true affinities: large differences from values determined from binding kinetics. Analytical Biochemistry. 234, 155–165 (1996).

80. Lazar, G.A., et al., Engineered antibody Fc variants with enhanced effector function. Proc. Natl. Acad. Sci. 103, 4005–4010 (2006).

81. Walkup, W.G., et al., A model for regulation by synGAP-⍺1 of binding of synaptic proteins to PDZ-domain ‘Slots’ in the postsynaptic density. eLife. 5, e16813 (2016).

82. Szeto, J., et al., Salmonella-containing vacuoles display centrifugal movement associated with cell-to-cell transfer in epithelial cells. Infect Immun. 77, 996–1007 (2009)

